# AI-based mining of biomedical literature: Applications for drug repurposing for the treatment of dementia

**DOI:** 10.1101/2024.06.06.597745

**Authors:** Aliaksandra Sikirzhytskaya, Ilya Tyagin, S. Scott Sutton, Michael D. Wyatt, Ilya Safro, Michael Shtutman

**Affiliations:** Department of Department of Drug Discovery and Biomedical Sciences, College of Pharmacy, University of South Carolina; Department of Computer and Information Sciences, University of Delaware; Department of Clinical Pharmacy and Outcomes Sciences, College of Pharmacy, University of South Carolina

## Abstract

Neurodegenerative pathologies such as Alzheimer’s disease, Parkinson’s disease, Huntington’s disease, Amyotrophic lateral sclerosis, Multiple sclerosis, HIV-associated neurocognitive disorder, and others significantly affect individuals, their families, caregivers, and healthcare systems. While there are no cures yet, researchers worldwide are actively working on the development of novel treatments that have the potential to slow disease progression, alleviate symptoms, and ultimately improve the overall health of patients. Huge volumes of new scientific information necessitate new analytical approaches for meaningful hypothesis generation. To enable the automatic analysis of biomedical data we introduced AGATHA, an effective AI-based literature mining tool that can navigate massive scientific literature databases, such as PubMed. The overarching goal of this effort is to adapt AGATHA for drug repurposing by revealing hidden connections between FDA-approved medications and a health condition of interest. Our tool converts the abstracts of peer-reviewed papers from PubMed into multidimensional space where each gene and health condition are represented by specific metrics. We implemented advanced statistical analysis to reveal distinct clusters of scientific terms within the virtual space created using AGATHA-calculated parameters for selected health conditions and genes. Partial Least Squares Discriminant Analysis was employed for categorizing and predicting samples (122 diseases and 20889 genes) fitted to specific classes. Advanced statistics were employed to build a discrimination model and extract lists of genes specific to each disease class. Here we focus on drugs that can be repurposed for dementia treatment as an outcome of neurodegenerative diseases. Therefore, we determined dementia-associated genes statistically highly ranked in other disease classes. Additionally, we report a mechanism for detecting genes common to multiple health conditions. These sets of genes were classified based on their presence in biological pathways, aiding in selecting candidates and biological processes that are exploitable with drug repurposing.

**Author Summary:** This manuscript outlines our project involving the application of AGATHA, an AI-based literature mining tool, to discover drugs with the potential for repurposing in the context of neurocognitive disorders. The primary objective is to identify connections between approved medications and specific health conditions through advanced statistical analysis, including techniques like Partial Least Squares Discriminant Analysis (PLSDA) and unsupervised clustering. The methodology involves grouping scientific terms related to different health conditions and genes, followed by building discrimination models to extract lists of disease-specific genes. These genes are then analyzed through pathway analysis to select candidates for drug repurposing.

## Introduction

Over the past decade, advancements in analytical methods have opened dramatic new opportunities to unveil hidden connections among complex networks (1). The advancement of Artificial Intelligence (AI) techniques enables researchers to query and analyze massive datasets, simulate experiments virtually, and generate scientific hypotheses through advanced analysis. Such tools have been used extensively by the pharmaceutical industry to lower the costs of drug discovery (2, 3). The development of novel therapeutic agents from idea to FDA approval involves substantial commitments of time and money (4). The FDA strictly evaluates efficacy and safety when approving new therapeutics, but the costs of newly approved therapeutics are now under intense scrutiny as the costs of new drug development have risen. Repurposing existing FDA-approved drugs for new indications can alleviate drug development costs in part because the safety profile and clinical experience already exists for the drug (5). Repurposing significantly accelerates the entire process by taking advantage of crucial steps that already occurred in the original FDA approval process (6). The key to the initial steps of drug repurposing is to find a connection between an existing drug and a disease of interest that is worth exploring preclinically or clinically for a new therapeutic indication. In many cases the necessary knowledge may already be present in biomedical literature; however, the connections between various pieces of information may not be obvious. To determine the hidden connections, we developed AI-based literature analysis tools: MOLIERE (7), followed by Automatic Graph Mining And Transformer based Hypothesis Generation Approach (AGATHA) (8). The recent development of AI allows automated extraction of valuable information from unstructured text such as scientific abstracts or articles, thus enabling efficient and scalable processing of textual data that dramatically saves time and effort compared to manual processing (9). AI algorithms can identify themes, topics, and clusters within a collection of documents, thus helping researchers to overcome challenging, time-demanding analysis of literature (10). In vast databases like PubMed, search results often overwhelm users with excessive information that is hard to curate without detailed, time intensive assessment. Natural language processing (NLP) techniques in AI frameworks not only speed up the research but also enhance extracting valuable knowledge from massive datasets, which is challenging to achieve manually (11). Drug repurposing research benefits from these advantages in many ways (5, 12, 13). Studying disease at the gene level remains a challenging task despite recent advancements in genomics and technology. NLP methods were successfully used for a variety of gene-related tasks including but not limited to the identification of unique anticancer targets (2), predicting cognitive decline (14), interpreting microbial genes (15) and others. To achieve successful results in NLP calculations, it is imperative to have high-quality training datasets, conduct preprocessing procedures to normalize the data and reduce noise, choose appropriate model architecture (such as recurrent neural networks (RNNs), convolutional neural networks (CNNs), or transformers), and optimize hyperparameters such as learning rate, batch size, dropout rate, and model architecture configurations.

AGATHA is an effective literature-based discovery tool capable of extracting relevant information by sifting through immense scientific databases including PubMed, which expands annually by over one million papers (16). This tool analyzes the collection of lexical elements (e.g., words, phrases, and lemmas) within each research article abstract to identify possible hidden connections among terms specified by the user. The likelihood of the potential connection is estimated using a multi-headed self-attention mechanism accounting for the spatial relationships between individual terms in a latent vector space, which we will refer to as the “AGATHA space”.

The AGATHA system pipeline can roughly be split into two stages: (1) semantic knowledge network construction and its embedding into a low-dimensional vector space and (2) transformer-based predictor training. These two stages can be used independently from each other, which we leverage in the current work here. The first stage results in a large multi-layered knowledge network, which connects individual units of information (such as the UMLS terms, semantic predicates or entities) with their corresponding literature sources. For example, a term representing “breast cancer” is connected to all sentences and semantic predicates containing this term. After construction, we perform network embedding, such that each node is assigned a learned vector representation (or coordinates) of 512 dimensions. This allows us to establish spatial relationships between individual concepts by computing distances between them (8). Terms that are logically connected, such as different stages of the same health condition or its type, tend to cluster together in AGATHA space. Conversely, terms that are logically distant from each other are positioned relatively far apart. This approach aims to facilitate an intuitive understanding of the relationships and connections between the many scientific terms and concepts within scientific literature.

The second stage results in a transformer-based predictor model, which is trained to prioritize meaningful associations between biomedical concepts above random noise. For each term-term association, it outputs a score within a unit interval indicating the likelihood of this association being biomedically relevant, based on the insights learned from scientific abstracts. When we use the AGATHA predictor in a one-to-many setting (like in this work), we obtain the probability distribution over a range of pairs, where the source term is fixed (e.g., a particular disease) and target terms represent a group of concepts of similar semantic type (e.g., list of different genes). Therefore, we can identify what genes are more likely to be associated with a particular disease and select the most prominent candidates for further downstream analysis.

To classify AGATHA outcomes, we applied multivariant statistical methods, including Partial Least Squares Discriminant Analysis (PLSDA) (17) (18). This method helps categorize and predict samples (diseases/genes) belonging to specific classes. PLSDA has advantages compared to other discriminant methods due to its ability to handle data with high variability and a power to reduce dataset dimensionality through the utilization of latent variables (19). In addition to the classification analysis, unsupervised clustering was utilized to unveil latent relationships that cannot be directly measured within the multivariate data. The combination of these steps followed by comprehensive pathway analysis helps to explain the biological significance of the classification outcomes and produces a final list of genes as candidates for drug repurposing.

In this work, we focused on application of our methodology to identify candidate drugs suitable to be repurposed as treatments for neurodegenerative diseases (20), which pose major challenges in healthcare as the seventh leading cause of death in the world (21). The term “neurodegenerative” covers a wide spectrum of neurocognitive conditions that despite their different pathologies often share common symptoms, in which dementia is a major outcome (22). Therefore, the analysis included a broad spectrum of neurodegeneration in the dementia domain to search for common themes and pathways.

Initially, the classification model facilitated the extraction of “dementia” genes, which were subsequently analyzed within the context of the pathways in which they participate. Once it was confirmed that the proposed method effectively extracts the necessary data, the same procedure was employed on the remaining non-neurodegenerative disease classes to obtain specific genes for each group. Next, after obtaining a list of genes that have a high likelihood to be associated with each disease, they were mapped in the Dementia class to assess their places on a probability scale. Genes for which known small molecules interact with the pathways of interest were prioritized to select FDA-approved medications or medications in experimental or investigational status. We chose a total of 38 drugs for potential repurposing with a focus on six of them, all of which have demonstrated effectiveness in treating diseases unrelated to central nervous system function.

## Results

### Data description

Diseases and conditions of interest were selected from the Disease Database provided by the Unified Medical Language System (UMLS) (23, 24) and combined into the Health Conditions Data Set (HCDS) comprising a total of 122 terms, which are categorized into seven groups: Dementia (24 conditions), Diabetes (12 conditions), Arthritis (9 conditions), Heart Conditions/Diseases (14 conditions), Hypertension (11 conditions), Cancer (12 conditions), and Substance Use Disorders (SUD) (40 conditions/substances). All the selected terms are formal names for diseases and health conditions and the last group (SUD) additionally contains the most common substances of abuse (**Table 1**).

**Table 1.** Health Condition Data Set. Diseases, conditions, and substances are grouped by the type and color coded regarding their attribution.

| # | Name | UMLS code | # | Name | UMLS code | # | Name | UMLS code | # | Name | Code | # | Name | UMLS code |
| --- | --- | --- | --- | --- | --- | --- | --- | --- | --- | --- | --- | --- | --- | --- |
| **Dementia** |  |  | 26 | Monogenic Diabetes | c3888631 | 52 | Atrial Fibrillation | c0004238 | 77 | Osteosarcoma | c0029463 | 103 | Methadone | c0025605 |
| 1 | Aids Dementia Complex | c0001849 | 27 | Steroid-Induced Diabetes | c0342269 | 53 | Cardiac Arrhythmia | c0003811 | 78 | Pancreas Carcinoma | c0235974 | 104 | Hydrocodone | c0020264 |
| 2 | Alcohol Amnestic Disorder | c0001940 | 28 | Type 1 Diabetes Mellitus | c0011854 | 54 | Rheumatic Heart Disease | c0035439 | 79 | Prostate Cancer | c0376358 | 105 | Meperidine | c0025376 |
| 3 | Alcohol-Induced Mental Disorders | c0033936 | 29 | Type 2 Diabetes Mellitus | c0011860 | 55 | Myocardial Infarction | c0027051 | 80 | Melanoma | c0025202 | 106 | Oxycodone | c0030049 |
| 4 | Alzheimer Disease, Early Onset | c0750901 | 30 | Maturity Onset Diabetes Mellitus In Young | c0342276 | 56 | Cardiomyopathies | c0878544 | **Substance abuse disorder** |  |  | 107 | Oxymorphone | c0030073 |
| 5 | Alzheimer Disease, Late Onset | c0494463 | 31 | Cystic Fibrosis Related Diabetes | c2242728 | 57 | Atrial Septal Defects | c0018817 | 81 | Alcohol Abuse | c0085762 | 108 | Diacetylmorphine | c001189 |
| 6 | Amyotrophic Lateral Sclerosis | c0002736 | 32 | Gestational Diabetes | c0085207 | **Hypertension** |  |  | 82 | Ethanol | c0001962 | 109 | Mitragynine | c0066619 |
| 7 | Corticobasal Degeneration | c0393570 | 33 | Wolfram Syndrome 1 | c4551693 | 58 | High Blood Pressure | c0020538 | 83 | Amphetamine | c0002658 | 110 | Hydromorphone | c0012306 |
| 8 | Dementia Associated with Alcoholism | c0236656 | **Arthritis** |  |  | 59 | Chronic Hypertension | c0745114 | 84 | Cannabicyclohexanol | c3492502 | 111 | Loperamide Hydrochloride | c0282221 |
| 9 | Dementia, Vascular | c0011269 | 34 | Arthritis | c0003864 | 60 | Essential Hypertension | c0085580 | 85 | Synthetic Cannabinoids | c0006864 | 112 | Delysid | c0024334 |
| 10 | Frontotemporal Dementia | c0338451 | 35 | Ankylosing Spondylitis | c0038013 | 61 | Malignant Hypertension | c0020540 | 86 | Cannabidiol | c0936079 | 113 | Ketamine | c0022614 |
| 11 | Hand | c4285693 | 36 | Psoriatic Arthritis | c0003872 | 62 | Hypertensive Crisis | c0020546 | 87 | Central Nervous System Depressants | c0007681 | 114 | Mescaline | c0025460 |
| 12 | Hiv Encephalopathy | c0276548 | 37 | Pyogenic Arthritis | c0003869 | 63 | Isolated Systolic Hypertension | c0745133 | 88 | Dextromethorphan | c0011816 | 115 | Ecstasy Abuse | c0743247 |
| 13 | Huntington's Disease | c0020179 | 38 | Rheumatoid Arthritis | c0003873 | 64 | Resistant Hypertension | c0745130 | 89 | Barbiturates, Benzodiazepines | c0004745 | 116 | Psilocybin | c0033850 |
| 14 | Lewy Body Disease | c0752347 | 39 | Fibromyalgia | c0016053 | 65 | Gestational Hypertension | c0852036 | 90 | Flunitrazepam | c0016296 | 117 | Salvia | c3668955 |
| 15 | Major Neurocognitive Disorder | c4087461 | 40 | Reactive Arthritis | c0085435 | 66 | Secondary Hypertension | c0155616 | 91 | Rohypnol | c0699927 | 118 | Phencyclidine | c0031381 |
| 16 | Mild Cognitive Disorder | c1270972 | 41 | Gout | c0018099 | 67 | Cerebrovascular Accident | c0038454 | 92 | Opium | c0029112 | 119 | Nandrolone | c0027368 |
| 17 | Mild Neurocognitive Motor Disorder | c2609165 | 42 | Degenerative Polyarthritis | c0029408 | 68 | Transient Ischemic Attack | c0007787 | 93 | Sodium Oxybate | c0037537 | 120 | Oxandrolone | c0029995 |
| 18 | Parkinson's Disease | c0030567 | **Heart disease/condition** |  |  | **Cancer** |  |  | 94 | Cathine | c0069021 | 121 | Oxymetholone | c0030072 |
| 19 | Presenile Dementia | c0011265 | 43 | Angina Pectoris | c0002962 | 69 | Breast Cancer | c0006142 | 95 | Tobacco | c0040329 | 122 | Testosterone Cypionate | c0076181 |
| 20 | Prion Disease | c0162534 | 45 | Heart Failure | c0018801 | 70 | Cervical Cancer | c0007847 | 96 | Nicotine | c0028040 |  |  |  |
| 21 | Senile Dementia | c0002395 | 46 | Congenital Heart Defects | c0018798 | 71 | Colorectal Cancer | c0009402 | 97 | Khat | c1386575 |  |  |  |
| 22 | Senile Psychosis | c1457889 | 47 | Heart Valve Disease | c0018824 | 72 | Leukemia | c0023418 | 98 | Benzoylmethylecgonine | c0009170 |  |  |  |
| 23 | Subcortical Vascular Dementia | c0393561 | 48 | Coronary Artery Disease | c1956346 | 73 | Lymphoma | c0024299 | 99 | Methylphenidate | c0025810 |  |  |  |
| **Diabetes** |  |  | 49 | Peripheral Artery Disease | c1704436 | 74 | Malignant Lung Neoplasm | c0242379 | 100 | Codeine | c0009214 |  |  |  |
| 24 | Prediabetes Syndrome | c0362046 | 50 | Coronary Arteriosclerosis | c0010054 | 75 | Malignant Neoplasm Of Skin | c0007114 | 101 | Morphine | c0026549 |  |  |  |
| 25 | Latent Autoimmune Diabetes in Adults | c1739108 | 51 | Disorder Of Pericardium | c0265122 | 76 | Neuroblastoma | c0027819 | 102 | Loperamide | c0023992 |  |  |  |

We hypothesized that there are spatial clusters within the AGATHA space that correspond to different groups of health conditions (**Table 1**). This implies that by mapping genes within the AGATHA space and analyzing their positions relative to disease groups, we can uncover previously unrecognized links between specific gene sets and health conditions. These genes are specifically evaluated for potential drug repurposing opportunities.

A general logical workflow is represented in ***Scheme 1*** below. Disease-categorized data from a variety of databases was mapped to the AGATHA space for further characterization. Then, classification methods were used to build a discrimination model that extracted four lists of genes: 1) genes specific for each disease class; 2) Dementia genes, highly ranked in other disease classes; 3) Disease genes, highly ranked in Dementia class, and 4) genes common for all diseases. These groups of genes were used for pathway analysis performed using **g:profiler** tool (25), which helped to select the candidates for drug repurposing evaluation.

**Scheme 1.**
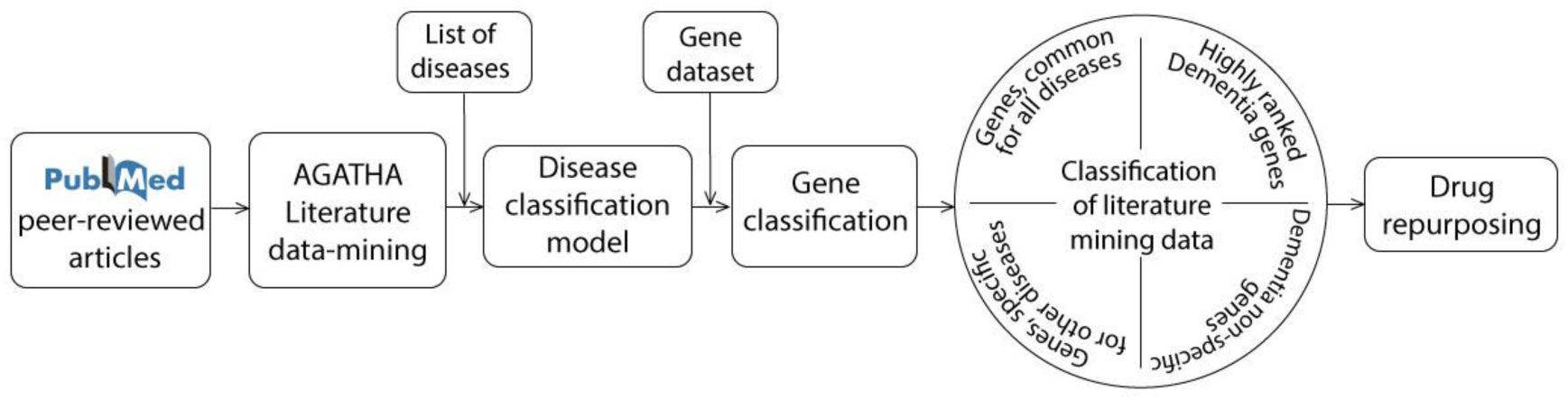
Workflow of gene selection for drug repurposing based on literature-mining strategy.

### Exploratory analysis of semantic links revealed by the AGATHA embedding space

The complex relationships between the selected health condition terms (26, 27) (**Table 1**), described across multiple diverse scientific articles, were assessed using AGATHA text mining software. Following semantic embedding, these health condition terms are represented as points within a high-dimensional latent space, the embedding space we named earlier as AGATHA space (see the first paragraph of the **Results** section). The coordinates of these terms are calculated to reflect their semantic properties in such a way that words or phrases with similar meanings are represented by points that are closer to each other within the space. The number of dimensions in an embedding space is typically influenced by the volume and complexity of the text data being analyzed. However, it is also determined by the specific requirements of the model and the task at hand. While a larger and more complex dataset might benefit from a higher-dimensional space to capture more nuanced semantic relationships, the choice of dimensionality also depends on computational constraints and the desired balance between detail and efficiency. In our case, preliminary studies indicated that an efficient embedding of the information contained within the PubMed database of scientific abstracts is achieved by using an embedding space with 512 dimensions (8). In **Figure 1**, we see the relative positions of HCDS groups as visualized in 3D space. This visualization is the result of condensing the original 512-dimensional data into a more comprehensible three-dimensional space using Principal Component Analysis (PCA). Two distinct clusters are formed by two non-overlapping sets of health conditions: SUD (green diamonds) and Dementia (red diamonds). The five remaining sets — Diabetes, Arthritis, Heart Diseases, Hypertension, and Cancer — form a tight spatial cluster that is separated from both the SUD and Dementia clusters. Subclusters corresponding to these five groups of health conditions remain distinguishable. However, they are positioned close to each other resulting in the overlap of certain groups. These observations have led us to hypothesize that the AGATHA space contains a spatial pattern characteristic of the health condition groups. In further sections we implement advanced statistics to identify and characterize such patterns.

**Figure 1.**
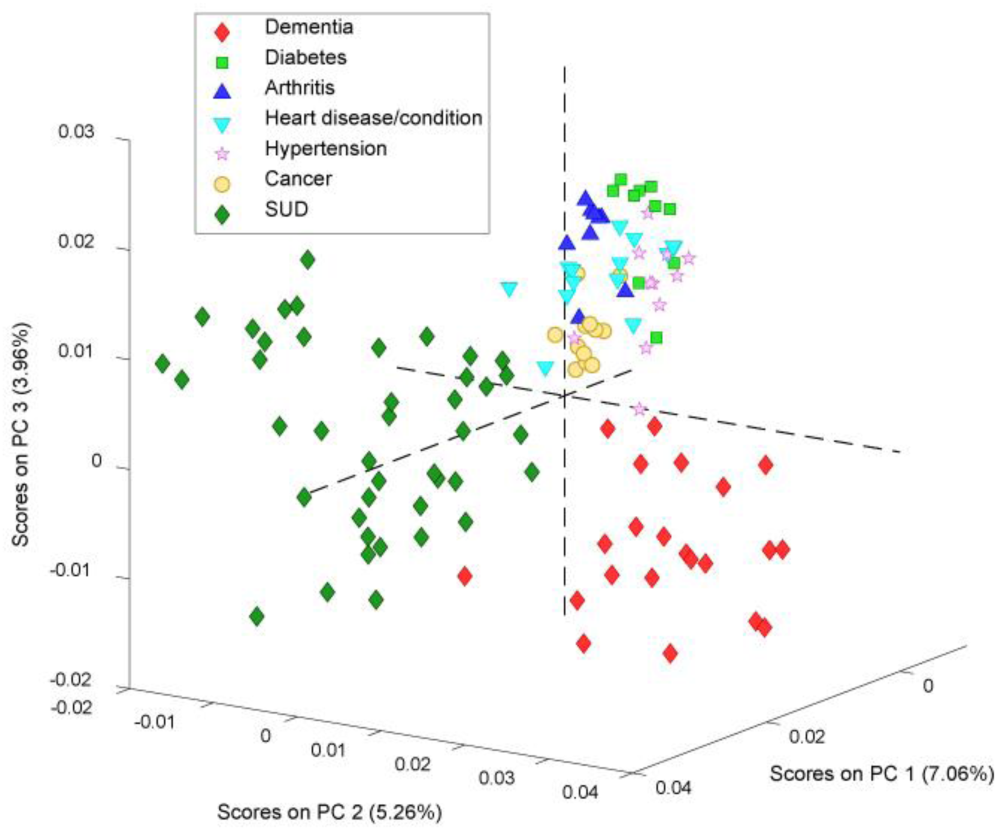
Principal Component Analysis of the Health Condition Data Set (HCDS).

### Classification analysis of health condition groups mapped to the AGATHA space

The validity of spatial patterns in the AGATHA space associated with health condition groups was tested using PLSDA, a standard partial least square classification approach. In this study, we leverage both the interpretability of the multiclass PLSDA models and their capability to effectively handle collinear data. The strong predictive performance of the PLSDA models was subsequently employed to investigate gene/health condition associations. PLSDA classification has been demonstrated to be a successful method for addressing multivariate data, offering tunable model complexity (18). We used PLSDA to build a supervised classification model (**Figure 2**) with classes defined by health condition groups (**Table 1**). Extensive preliminary classification trials (not reported here) enabled the identification of optimal data preprocessing and classification parameters. For the final classification model, the input matrix containing coordinates in 512-dimensional AGATHA space for all health conditions was preprocessed using normalization by the total area and auto-scaling. Despite the high dimensionality of the input data generated through complex algorithms implemented in the AGATHA text mining software, four latent variables were sufficient to produce a robust classification of health conditions. Generally, latent variables are calculated so that each subsequent latent variable captures the shared variance remaining after the extraction by the previously calculated variables. A total of 16.31% of the data was covered by the first four latent variables.

**Figure 2.**
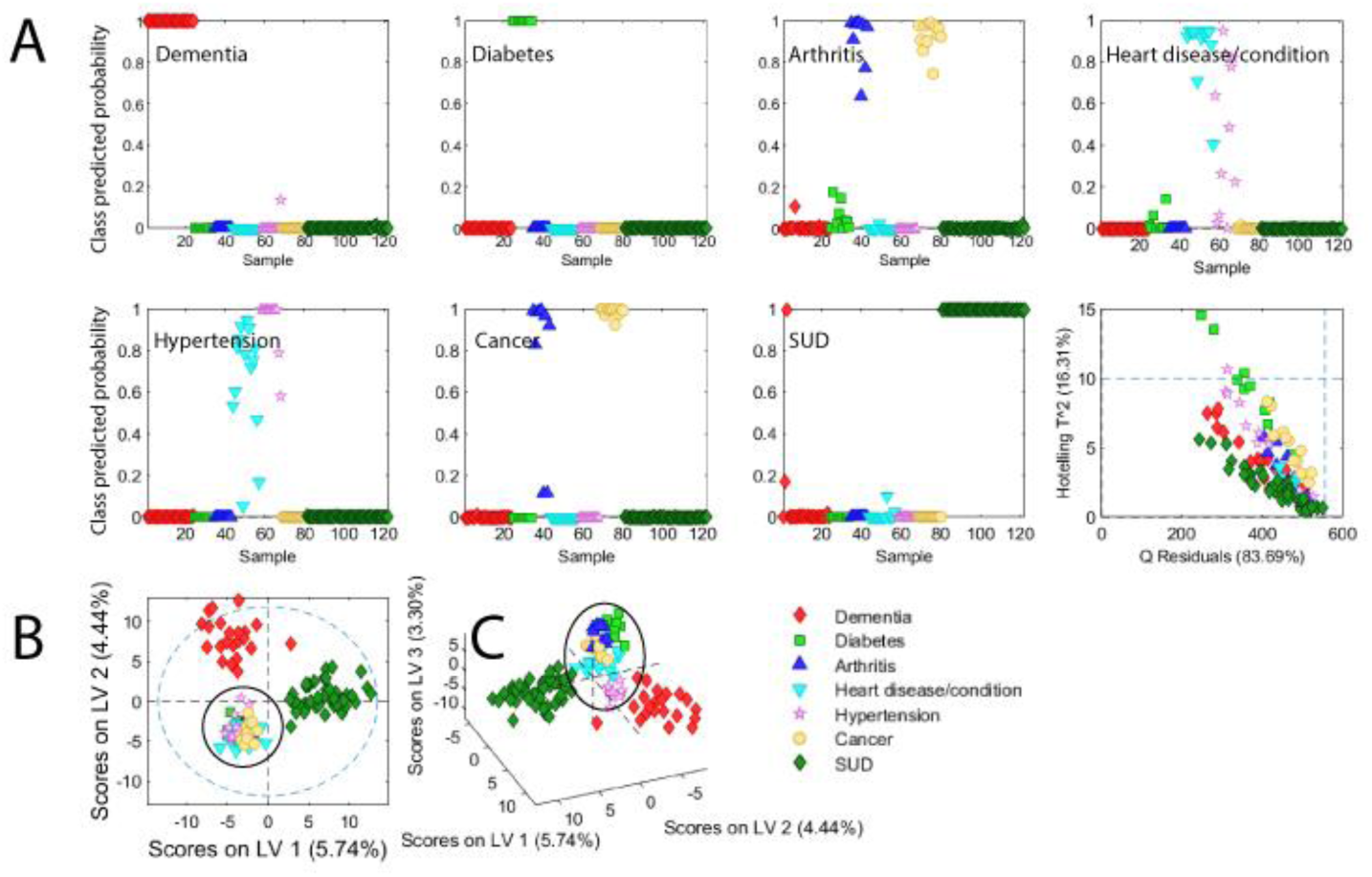
Results of cross-validated PLSDA classification analysis. **A** – Probabilities for each disease to be classified as an assigned class, Q Residuals vs T^2^ Hoteling plot error plot. PLSDA plots represent diseases separation for the first two (**B**) and three (**C**) latent variables.

The stability of the PLSDA classification model was verified using the Venetian blinds cross-validation approach, which involves dividing the data into ten equally sized folds. The final classification model effectively categorizes health conditions into seven predefined groups, as shown in **Figure 2.A**, demonstrating cross-validated sensitivity and specificity parameters within the range of 0.786 to 0.990. As expected from the exploratory analysis (**Figure 1**), the Dementia and SUD classes exhibited the best classification performance. The Dementia panel in **Figure 2.A** reveals that all health conditions initially selected for the Dementia group have a probability close to 100% of being classified as part of the Dementia class. Note that the 0-1 range on the Y-axis in the panel corresponds to a 0-100% range of probabilities. These observations suggest that all the health conditions we originally selected for the Dementia group constitute a distinct spatial cluster in the AGATHA space. Furthermore, should a small portion (one-tenth) of the Dementia set be omitted as ’unknown’ health conditions during the training phase, these ’unknowns’ are likely to be accurately classified in subsequent classification analysis. Interestingly, this observation holds true not only for Dementia and SUD, but also for Diabetes. In **Figure 1**, Diabetes is the most distant from the Dementia and SUD groups, neighboring but not overlapping with the other groups. Discriminating between the Arthritis, Cancer, Heart Disease/Condition, and Hypertension groups is also achievable, as shown in the **Discussion** section, despite overlapping regions. Two distinct pairs of groups can be identified: the Heart Disease/Condition and Hypertension pair, and the Arthritis and Cancer pair (**Figure 2.A** corresponding panels). Health conditions originally selected for these four groups overlap and show a non-zero probability of being assigned to another class of the pair. The proximity and overlap of the Heart Disease/Condition and Hypertension groups can be explained by the shared physiological characteristics of these disorders(28). There are also certain connections between Cancer and Arthritis, such as associations with chronic inflammation and paraneoplastic arthritis(29). While considering the physiological origins of connections within these two pairs of health condition groups is beyond the scope of this proof-of-concept study, we will later demonstrate that, upon more detailed analysis of the overlapping groups (see black circles in **Figures 2.B** and **C**), it is possible to build a robust classification model for discrimination of all groups.

### Assigning human genes to health condition groups using AGATHA latent space and advanced statistical methods

Text mining algorithms provide a unique opportunity to connect scientific concepts using lexical context. As demonstrated above, the AGATHA algorithm successfully condensed scientific information within the PubMed database, capturing lexical context characteristics of the health condition groups we selected for this proof-of-concept study. In this section, we explore the ability of the AGATHA system to uncover hidden connections between genes and health conditions. This was achieved by mapping all human genes to the AGATHA space and categorizing them into health condition groups using the PLSDA classification model. This step was followed by an in-depth analysis of the identified gene clusters in the context of diseases, physiological pathways, and drugs known to interact with these pathways.

The complete list of human genes, mapped to the AGATHA space as a matrix with 20,889 rows and 512 columns, was analyzed using the PLSDA model developed for HCDS. **Figure 3** illustrates the distribution of genes among all disease classes, with the color bar showing their attribution to the Dementia class in each category.

**Figure 3.**
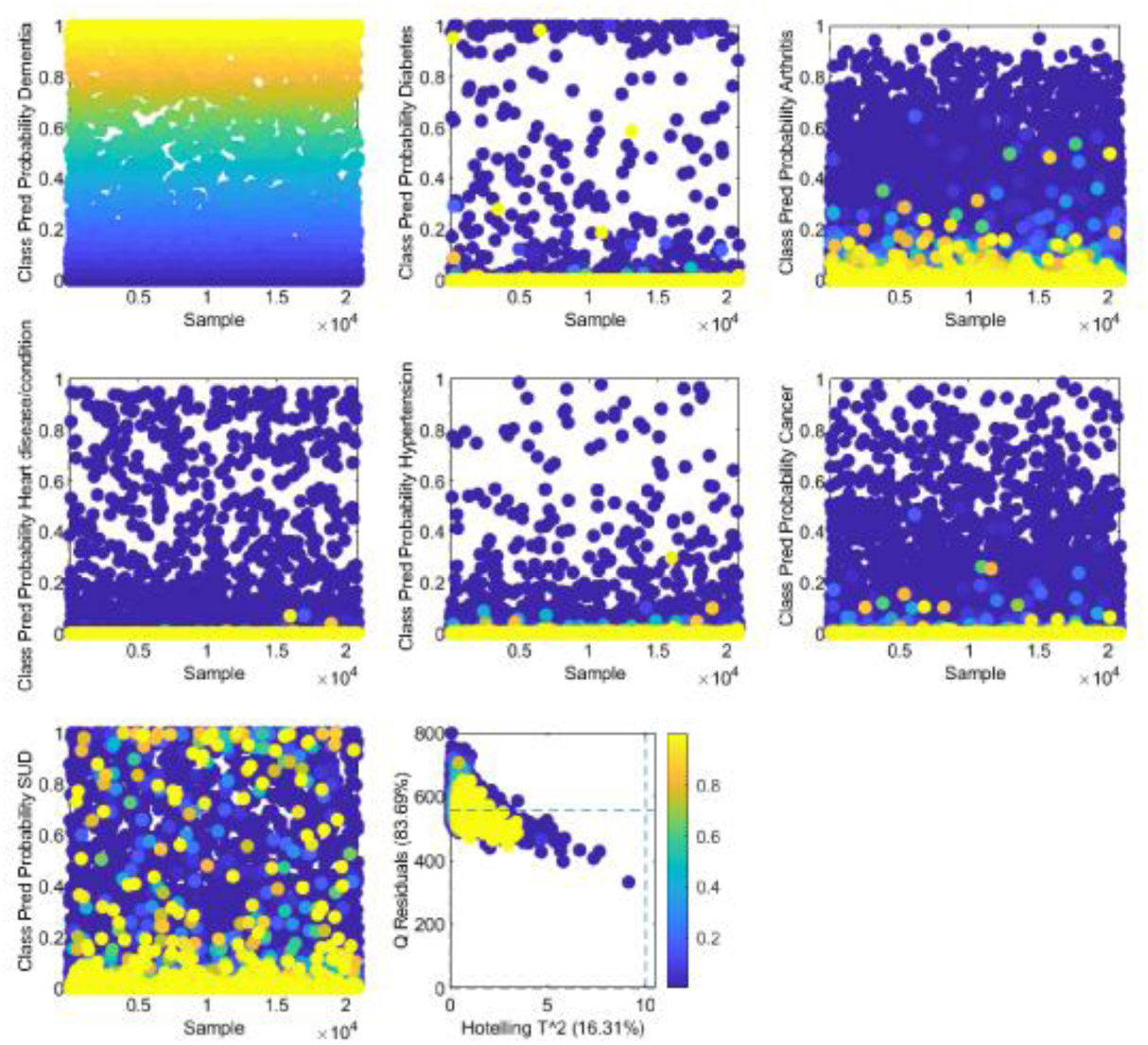
PLSDA prediction of gene distribution among diseases A – Dementia, B – Diabetes, C – Arthritis, D – Heart condition/disease, E – Hypertension, F – Cancer, G – SUD, H – Q residuals vs Hotelling T^2 plot. Genes colored by the prediction probability to be assigned to Dementia.

As seen in the figure above, the distribution of Dementia genes does not follow the same pattern across all other classes. At this point, the evaluation of gene distribution in the Dementia class is necessary to show that the calculated model is coherent from the biological point of view. To achieve this, genes with a probability exceeding 80% were analyzed using hierarchical clustering. This approach aided in investigating the internal structure of the data, followed by the pathway analysis of the calculated clusters.

A total of 1079 genes with high probability to be associated with dementia were identified by the classification model and further subjected to unsupervised cluster analysis using agglomerative Ward’s method with a total of four principal components and Mahalanobis distance that accounts for the variations of multivariate data (**Figure 4**).The selected threshold allowed for gene separation into four distinct clusters that were further subjected to a pathway analysis to justify the biological meaning of data distribution.

**Figure 4.**
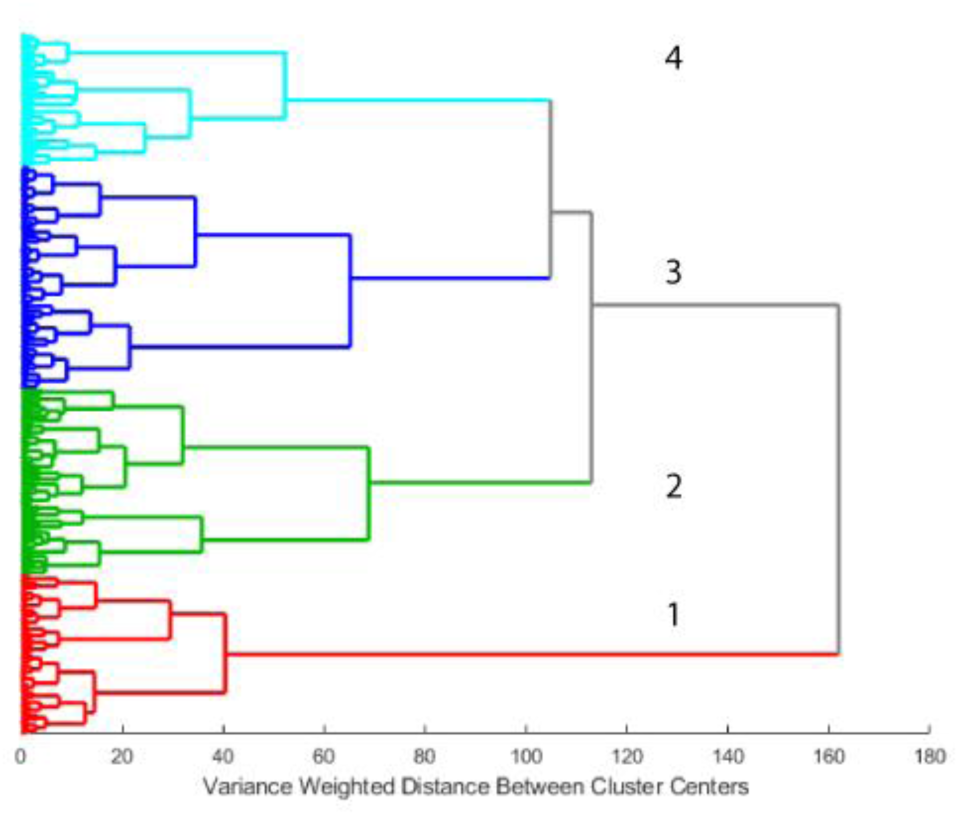
Hierarchical clustering analysis using Ward’s method. Different colors illustrate gene groupings characterized by various disease markers.

The dendrogram in Figure 4 shows four well-separated gene clusters, each defined by specific biological processes and mechanisms. These assignments were determined through pathway analysis, which can be summarized as follows:

Compared with **Clusters 2-4, Cluster 1** is separated from the remaining data at the initial threshold level, indicating its unique characteristics. Pathway analysis revealed that the processes within Cluster 1 do not show direct connections to specific physical or behavioral anomalies and cannot be linked to any specific disease category. However, this information is still useful when genes from this cluster are mapped as high-ranked in other disease categories. **Cluster 2** has a strong connection to several kinds of pathways known to be altered in neurodegenerative conditions including but not limited to Alzheimer’s disease, Amyotrophic lateral sclerosis, Parkinson’s disease, and Apoptosis - multiple species as labeled by g-profiler. **Cluster 3** has characteristics analogous to Cluster 2, such as nervous system development, presynaptic endocytosis, neuron projection organization, visual perception, regulation of neuron projection development, dendrite morphogenesis, and regulation of cell projection organization, and neuron projection. Some of the pathways discussed here are relevant to conditions such as Parkinsonism, disturbances in higher cognitive functions, central motor function disruptions, Ataxia, speech impairments related to the nervous system, and the life cycle of the HIV-1 virus. **Cluster 4** is different from the other three by having pathways related to substance abuse. It includes nicotine, cocaine, amphetamine addictions, alcoholism, and some pathways connected to the nervous system such as neuroactive ligand-receptor interaction, dopaminergic synapse, retrograde endocannabinoid signaling, axon guidance, and many more (***Supporting Table 1. Summary of pathways and genes for the Dementia class***).

As a result, we acquired a list of genes with a high probability of being connected to Dementia as well as being simultaneously highly ranked in the remaining six classes. After evaluation by the GeneCards database, these genes were separated into one specific group (***Supporting Table 2. Dementia non-specific genes, highly ranked in other disease groups***). Subsequently, highly ranked genes from the Diabetes, Arthritis, Heart, Hypertension, Cancer, and SUD classes were extracted for each disease/condition and were mapped in the Dementia group (**Figure 5**). For most of the classes, they were not presented at the top of the probability scale, so only the ones with the highest likelihood to be connected to Dementia were combined. (***Supporting Table 3. Disease-specific genes, highly ranked in Dementia***).

**Figure 5.**
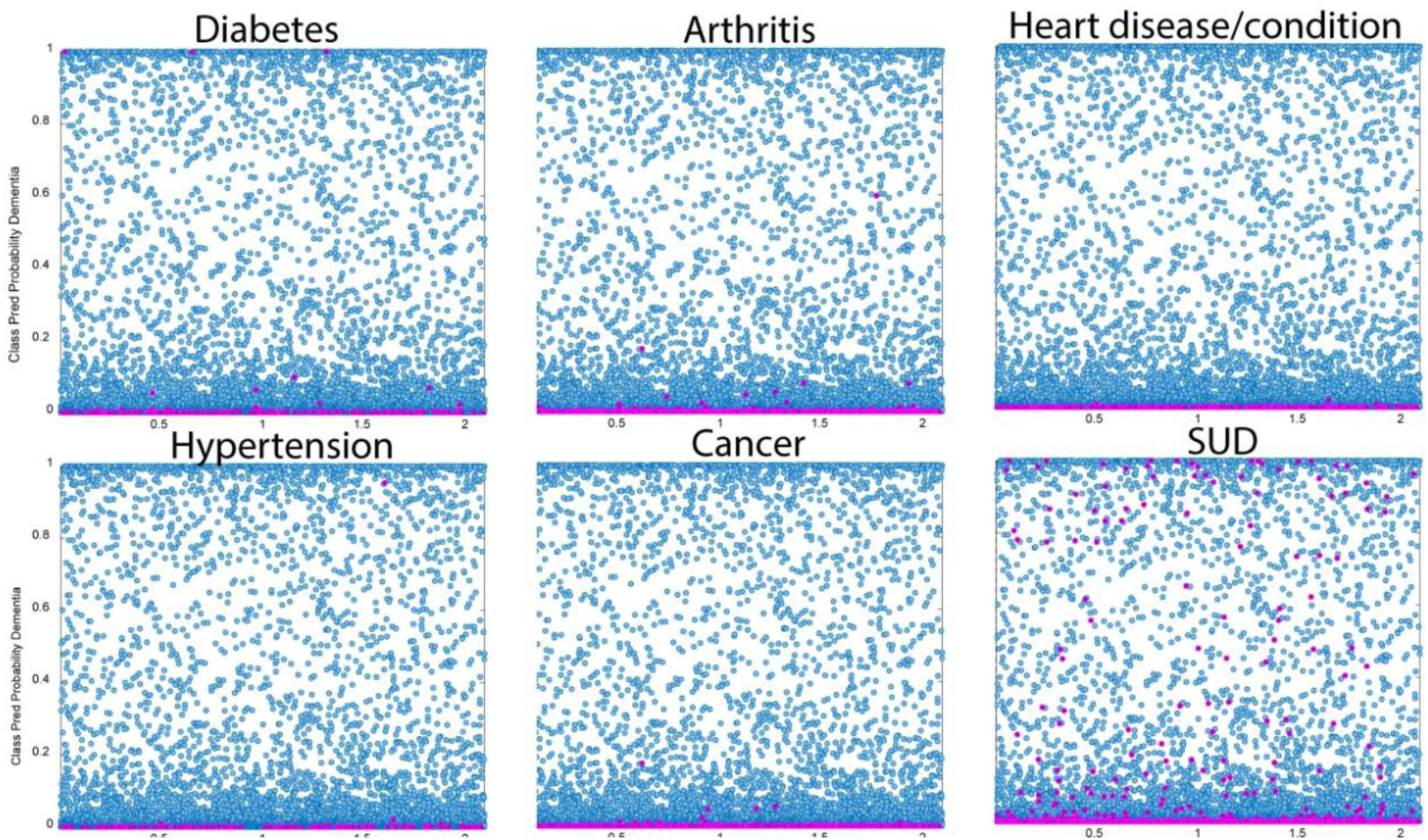
PLSDA prediction. Genes highly ranked for every disease (pink markers) mapped by their prediction probability for the Dementia group.

In addition to the described analyses, we followed the same procedure and extracted the top 1079 genes for each disease resulting in six lists: Diabetes, Arthritis, Heart conditions/diseases, Hypertension, Cancer, and SUD. These lists were reduced by retaining only genes distinct to each specific class of disorders to remove excessive overlapping of biological information. Based on these results, pathway analysis was performed for the specific gene lists and summarized in **Table 2**.

**Table 2.** Main pathways identified for the lists of specific genes for all disease classes.

| Disease | Number of genes | Significant pathways |
| --- | --- | --- |
| Dementia | 518 | Glutamatergic synapse pathway, neuron development, axon development, Alzheimer disease, Amyotrophic lateral sclerosis, Dementia, Brain atrophy, Frontotemporal dementia, Parkinsonism, regulation of synaptic plasticity |
| Diabetes | 109 | Insulin and sugar metabolism-related pathways, glucose transmembrane transporter activity, insulin secretion, monosaccharide transmembrane transport, Type II diabetes mellitus, Abnormal hemoglobin |
| Arthritis | 32 | Immune response-activating cell surface receptor signaling pathway, activation of immune response, regulation of B cell receptor signaling pathway, protein deglutamylation, Abnormal lymphocyte proliferation |
| Heart | 67 | Blood circulation, regulation of blood circulation, regulation of heart contraction, regulation of heart rate, heart development |
| Hypertension | 0 | N/A |
| Cancer | 18 | DNA damage sensor activity, Homologous recombination, RAD51B-RAD51C-RAD51D-XRCC2-XRCC3 complex |
| SUD | 397 | Nicotine addiction, morphine addiction, cocaine addiction, Common pathways underlying drug addiction, Drug metabolism - cytochrome P450, drug adme, Steroid hormone biosynthesis, dopamine neurotransmitter receptor activity, dopamine secretion, modulation of chemical synaptic transmission, Retrograde endocannabinoid signaling |

### Pathway overview

Pathways specific for **Dementia** class based on selected genes are described above.

### Diabetes (109 genes)

The diabetes group included insulin and sugar metabolism-related pathways, glucose transmembrane transporter activity, insulin secretion, monosaccharide transmembrane transport, Type II diabetes mellitus, abnormal hemoglobin, and others, as well as general pathways that can be assigned to a variety of conditions.

### Arthritis (32 genes)

The arthritis group included a variety of autoimmune and inflammatory diseases, which is reflected by the list of different characteristic pathways: immune response-activating cell surface receptor signaling pathway, activation of immune response, regulation of B cell receptor signaling pathway, protein deglutamylation, abnormal lymphocyte proliferation, and others.

### Heart condition/disease (67 genes)

The group included pathways such as blood circulation, regulation of blood circulation, regulation of heart contraction, regulation of heart rate, heart development, and others. By nature, some of these pathways are related to hypertension which can explain an absence of specific genes in the Hypertension cluster.

### Hypertension (0 genes)

There were no specific genes identified for the current group. Hypertension can be caused by various factors. It is related to the function of the heart, diabetes due to damaged arteries, excessive alcohol consumption, and others. Some of these causes were considered by the rest of the six classes, so the absence of hypertension-specific genes is not surprising and can be investigated further.

### Cancer (18 genes)

The cancer group can be described by several pathways, related to DNA damage sensing activity(30). This class includes the RAD51B-RAD51C-RAD51D-XRCC2-XRCC3 complex, in which inactivating mutations predispose to breast, ovarian and prostate cancers(31).

### SUD (397 genes)

The selected SUD group was composed of various addictions and individual substances. This was reflected in identified pathways that included chemical carcinogenesis - DNA adducts, nicotine addiction, morphine addiction, cocaine addiction, Common pathways underlying drug addiction, Drug metabolism - cytochrome P450, drug ADME (absorption, distribution, metabolism and excretion), steroid hormone biosynthesis, dopamine neurotransmitter receptor activity, dopamine secretion, modulation of chemical synaptic transmission, retrograde endocannabinoid signaling and many others.

### Drug repurposing

The classification model predicted the list of genes with high potential to be associated with Dementia. These hidden connections are selected based on the learned patterns and relationships in the data indirectly revealing acquaintances between terms. Further developments in understanding these relationships will require additional interpretation and analysis beyond the model itself. We selected a list of potential drugs for repurposing analysis based on the presence of genes specific to six groups of diseases within the Dementia class using GeneCards and Comparative Toxicogenomics Database (CTD) (32). Chosen medications were additionally verified by the DrugBank database (33) to track their approval status as well as the stage of Clinical trials (**Supporting Table 4**). Finally, **Table 3** summarizes the most significant candidates for drug repurposing based on the combination of statistically predicted gene/disease connections discovered by the classification model and gene/drug connections identified using databases listed above. Specifically, Bosentan, Mecamylamine, and Methylphenidate are the most compelling candidates, ranking at the top of the lists of small molecules for their predicted targets (EDNRB, CHRNA4, SLC6A3). Every drug was selected from the list of FDA-approved medications that target specific genes predicted by the classification model. We illustrate the pharmacological use of these drugs, and the level of clinical data as well as provide a literature-based explanation of their potency.

**Table 3.**
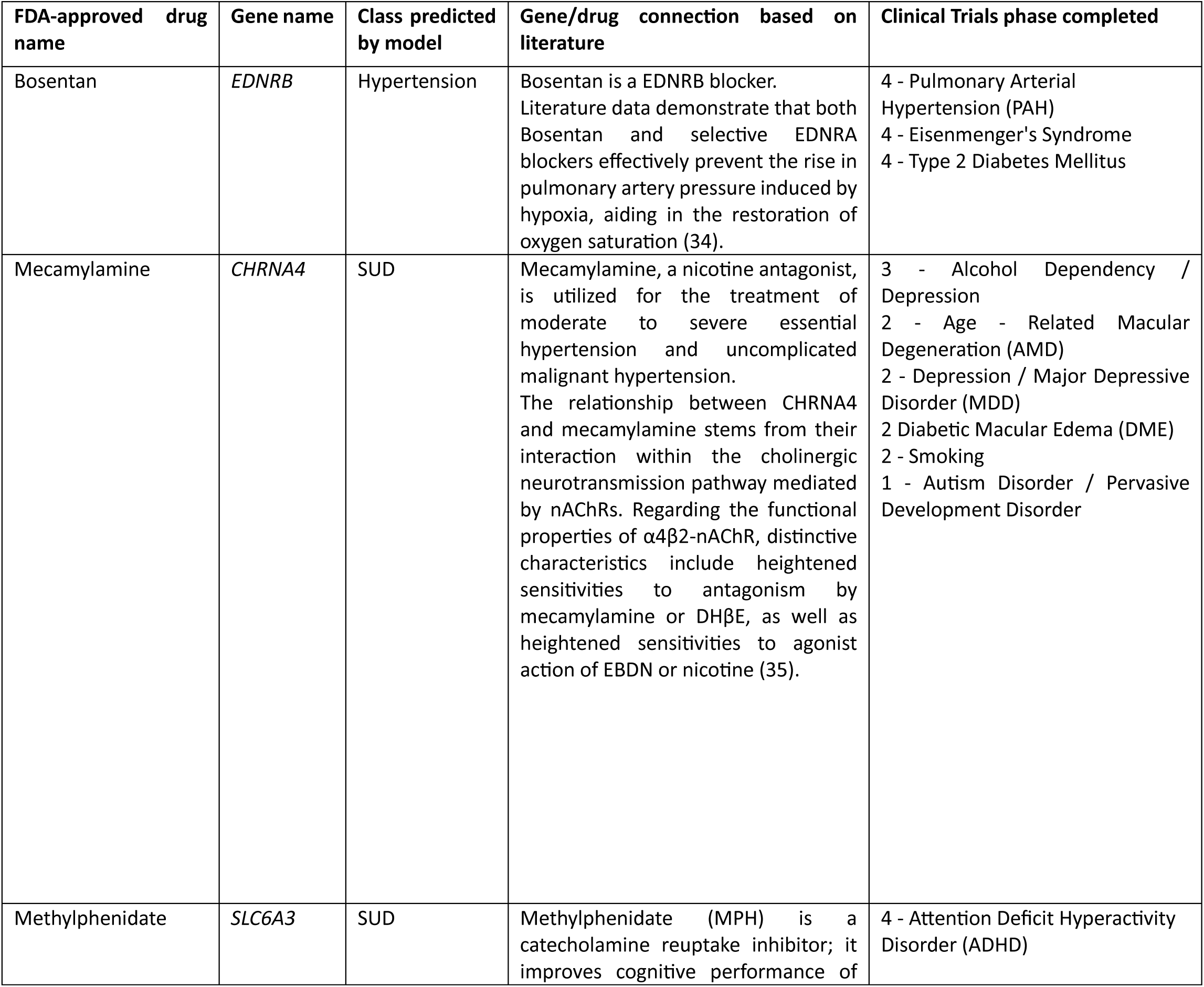
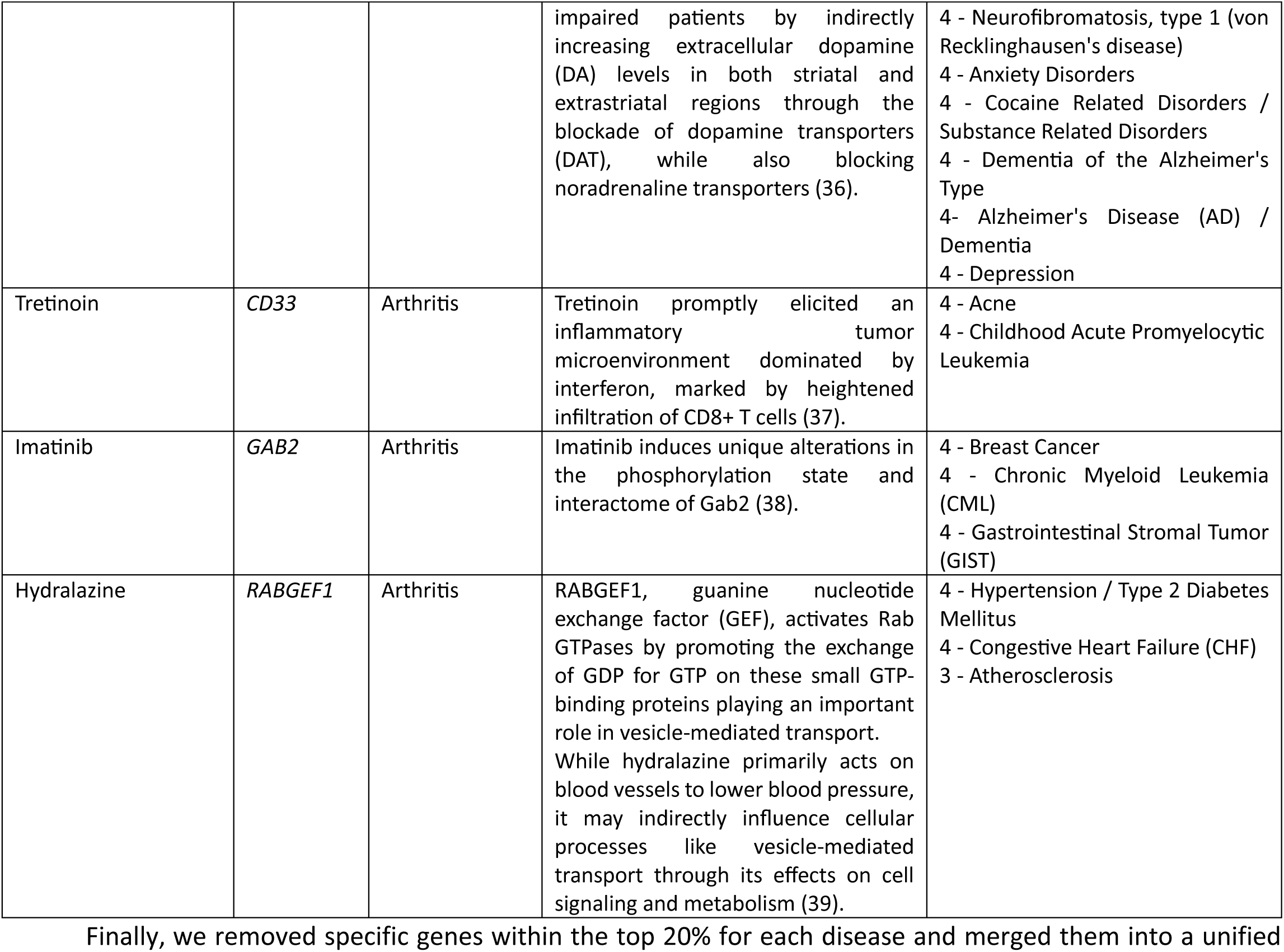
Drugs for repurposing for dementia treatment.

Finally, we removed specific genes within the top 20% for each disease and merged them into a unified list of common genes, totaling 1616 items (**Supporting table 5. Common genes and their pathways**). Excluding specific information allowed for the identification of general biological processes that can be applicable across various diseases or conditions, such as protein binding, catalytic activity, response to stimuli, biological regulation, cell communication, membrane involvement, cytoplasmic activities, and more. As of now, this list may not serve as a basis for future target selection, but it effectively illustrates the shared nature of diseases.

## Discussion

Over the last decade, various literature-mining methods were introduced for biological analysis. AI technology provides researchers with an opportunity to perform experiments with biomedical entity normalization applied to multiple datasets (40). This facilitates the identification of intricate gene citations in scientific articles and books (41) and aids drug repurposing efforts (42, 43). Previously we introduced literature mining methods (7, 8, 44) and demonstrated their potential application in many areas including the introduction of a new use for existing approved drug therapies (45). As a result of this project, a list of potential drugs for dementia treatment was extracted by AGATHA and advanced statistical analysis. The method identified hidden connections and pathways related to different diseases and neurodegeneration specifically. AGATHA-calculated variables for 122 diseases were separated into seven classes to calculate the PLSDA classification model. Initial discrimination showed that Dementia and SUD were separated from the rest of the group, agglomerating well-defined clusters when other diseases stay in a uniform cloud. However, when these two classes are excluded from the dataset, the rest of the diseases separate without any overlap (**Figure 6**). This illustrates the potential of the method to be applied for future projects studying other diseases or gene combinations.

**Figure 6.**
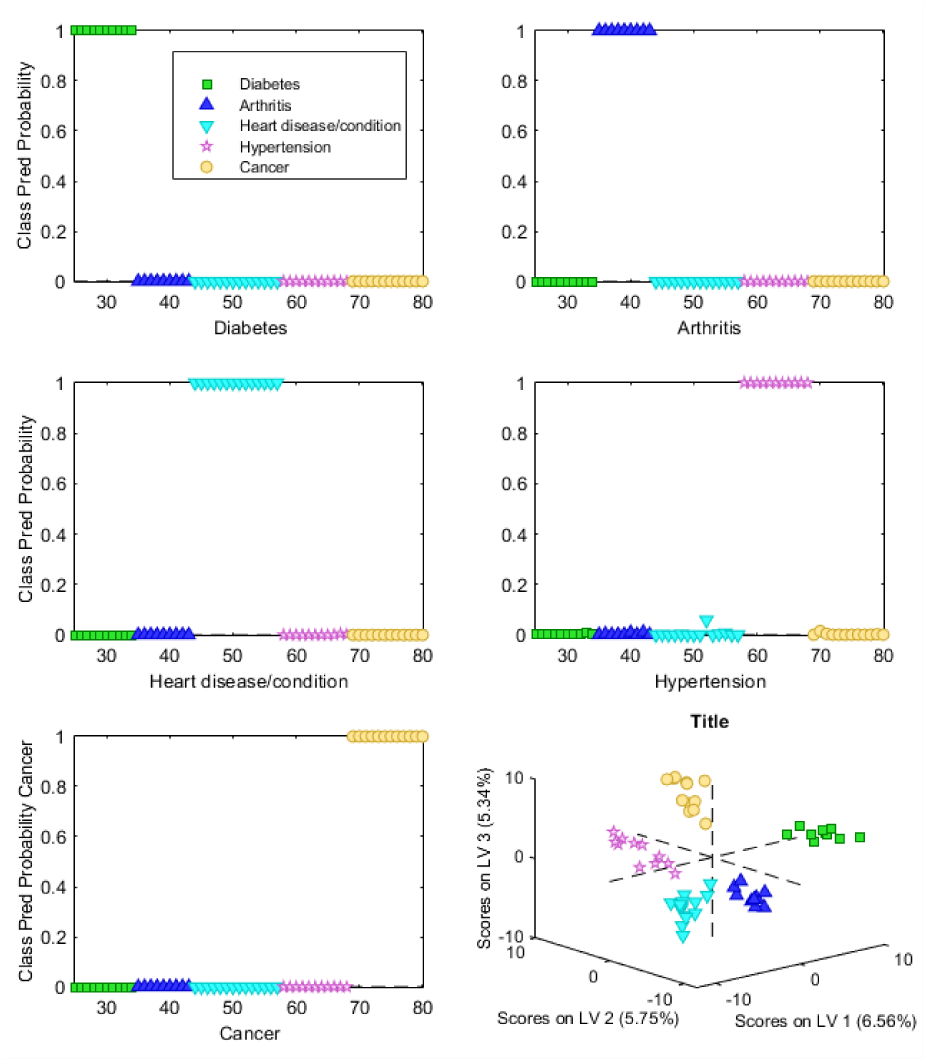
Class predicted probability for Diabetes, Arthritis, Heart disease/condition, Hypertension and Cancer groups.

In the next step 20889 genes were classified by the PLSDA model, which revealed different distribution patterns among the classes (**Figure 3**). It appears that in most cases Dementia genes are present at the bottom of the probability scale. It was noted that certain Dementia genes within the top 20% have a probability of being associated with SUD, with only three showing a connection to Diabetes. These genes are not prevalent in the top tier of the other four classes (**Figure 5**). As a result, a close look at the genes on top of the Dementia class showed elements of neurodegeneration as well as substance abuse. A total of 1079 genes (top 20%, **Figure 2A, Dementia plot**) were subjected to the pathway analysis which proved their belonging to that class, since they play crucial roles in numerous vital biological processes. Notably, they are involved in the Glutamatergic synapse pathway, which contributes to ensuring proper brain function. Disturbances in glutamate transmission or the improper regulation of glutamate receptors have been linked to various neurological disorders, such as epilepsy, Alzheimer’s disease, and schizophrenia. On the other hand, it has been shown that changes in metaplasticity of glutamatergic synapses play a significant role in the development of chronic SUD (46). In addition, it is known that tryptophan metabolism can have implications in the context of substance abuse due to its role in the production of neurotransmitters, including serotonin (47), which was shown in studies of patients with alcohol use disorder (48, 49). The same pathway leads to development of Alzheimer’s disease due to the inhibition of various enzymes responsible for the biosynthesis of β-amyloid (50). Thus, the genes that have a higher probability of being associated with Dementia can serve as potential targets for future drug repurposing due to their shared nature between SUD and Dementia based on the discovered pathways. As the result of mapping statistically allocated Dementia genes in the remaining classes, we obtained a list of genes highly ranked in other diseases.

A selection procedure was performed for the Diabetes, Arthritis, Heart conditions/diseases, Hypertension, Cancer, and SUD classes extracting the same number of genes as was performed for Dementia. Highly ranked genes in every group were then mapped in the Dementia class to evaluate their positions. This resulted in a separate list of genes that are not necessarily specific for any of the selected types of neurodegenerative disorders but have higher scores in general. Based on acquired information, the list of potential drugs for repurposing was created using GeneCards and CTD (**Table 3**). The suggested method enabled us to explore textual data from various angles. Apart from examining the interconnection of genes, it facilitates the identification of genes unique to each type of disease (**Table 2**).

The exploration of genes common for all 122 groups revealed their tendency to be present in pathways included in many biological processes simultaneously, proving the accuracy of the proposed method. The pathways disclosed in this list have a wide range of meanings and can be attributed to many processes or disorders. These similarities could be potentially used in future steps of the research project to discover new hidden connections. To summarize, the combination of the literature-mining method AGATHA, coupled with advanced statistical analysis allowed for the separation of the different lists of genes: Dementia genes, highly ranked in other disease classes, Disease genes, highly ranked in Dementia class, genes specific for every disorder, genes common for all diseases. This information was used for the selection of potential drugs for repurposing and has the potential of being used for future experiments involving finding new common pathways, selecting specific genes within the same group of diseases, or creating a robust automatic prediction method for the different inquiries.

## Materials and methods

AGATHA is an open-source algorithm available at: https://github.com/IlyaTyagin/AGATHA-C-GP. The operational principles of AGATHA are detailed in the **Supporting Information** section. Statistical analysis was performed on a multidimensional dataset using MATLAB R2023a software from MathWorks (Natick, MA) and the PLS Toolbox from Eigenvector Research, Inc. (Wenatchee, WA). Pathway analysis was conducted using g:profiler tool (25). Gene characterization and selection of potential drugs were achieved with the help of GeneCards (51) and CDT(32) databases.

Principal Component Analysis (PCA) (52) was primarily used for dimensionality reduction and preliminary data structure analysis. This widely-employed method is based on transforming the introduced data into a set of principal components that describe the variance of the data. It involves a series of mathematical steps, including calculating the covariance matrix of the data, computing its eigenvalues, and subsequently reducing the dimensionality of the data. The prepared dataset was normalized by the total area, and auto-scaled by the division of each of the 512-column in the calibration matrix by its standard deviation. Subsequently, cross-validation was performed using the Venetian Blinds approach, consisting of 10 splits with a blind thickness of 1. The achieved model showed a clear separation between the Dementia and SUD classes with the rest of the data falling into a single cohesive group. However, the cross-validated Root Mean Square Error for this model was extremely low (0.00231327), which indicates a good accuracy of the model.

The classification model was calculated using the PLSDA method (17), which can deal with heterogeneous data and describe it by only a few Latent Variables (LV). LVs are calculated using regression coefficients, determined for each component, and followed by estimating their positions in the PLSDA space. To moderate the risk of overfitting, Venetian-blinds cross-validation was employed. This method involves partitioning the data into k equal-sized segments and alternately using them as training and validation sets. Alongside dimensionality reduction, PLSDA ensures that the calculated components possess unique information by being orthogonally opposed to each other.

Unsupervised hierarchical clustering analysis was applied on part of the Dementia-classified data to group similar values into clusters based on their common characteristics. In cluster analysis, pairs of samples with the smallest distance between them are identified and merged without knowledge of class origin. These similar clusters are then grouped together in dendrogram visualization to provide a clearer representation. Ward’s method was used to minimize the variance within each cluster by evaluating the differences between merging two groups of data (53). This approach is especially efficient when handling high-dimensional data, such as our disease/coordinate or gene/coordinate sets, or when clusters are more likely to exhibit equal variance within them. These differences were estimated by the sum of squared deviations from the mean (variance) after merging the clusters (Mahalanobis distance).

## Competing interests

The authors have declared that no competing interests exist.

## Funding information

This project was supported by awards from NIH R01DA054992 (MS, MDW).

## Acknowledgments

We acknowledge the support of NIH for their financial assistance, and the College of Pharmacy for providing the necessary infrastructure and resources. Also, we would like to thank Dr. Vitali Sikirzhytski for the insightful feedback and suggestions, which significantly improved the quality of this manuscript.

## Notes

### Competing Interest Statement

The authors have declared no competing interest.

