## Supporting material. Methods for "AI-based mining of biomedical literature: Applications for drug repurposing for the treatment of dementia"

**S1. AGATHA system**

AGATHA stands for Automatic Graph mining and Transformer-based Hypothesis generation Approach, and it is meant to provide the domain experts in biology with insightful information about potential connections between biomedical terms. It does so by analyzing the large corpus of scientific literature in fully automatic manner. We present a brief description of the system below.

AGATHA system needs a textual corpus to get started which is obtained from MEDLINE database in a form of records, containing scientific abstracts from biomedical domain. These abstracts are then pre-processed with a list of NLP tools, such as the NLM-developed tool SemRep (1) and SciSpacy (2), to first split them into sentences and extract meaningful information from them, such as semantic predicates, lemmas, phrases and UMLS terms.

All this information is used to construct a semantic graph, which is composed of nodes of different types and edges between them. Edges connect sentences within the same abstract if they are consecutive or semantically relevant based on their text embeddings. There are also edges between sentences and the components extracted from them that were mentioned earlier.

This network is embedded with Pytorch BigGraph framework (3), such that each node is assigned a vector representation of dimensionality $d=512$. The goal is to have nodes sharing similar neighbors to be closer together in a vector space. These quantitative or functional relationships can be evaluated by the similarity measure S parameter and expressed as a biased transformed dot product:

$$S\left( a,b \right)= \hat{a}_{1}+\hat{b}_{1}+ T_{1}^{uv}+\sum_{i=2}^{d} \hat{a}_{i}(\hat{b}_{i}+ T_{1}^{uv})$$

where $d$ – number of dimensions, $\hat{a},\hat{b}$ – node embeddings and $T^{uv}$ – relation transformation vector.

The node embeddings are then used to train the predictor model, which is based on transformer architecture. The training goal is to learn a scoring function, which ranks a pair of biomedically relevant UMLS concepts above a pair with no evident biomedical relation. This is achieved by using published subject-object pairs as a positive training set and composing negative samples from the graph, similar to link prediction algorithms.

The objective function is margin ranking loss, which is meant to emphasize the scoring difference between positive and negative samples, making the model assign higher scores to positive samples compared to negatives:

$$L\left( a,b \right)=\sum_{n\in N} \max(0, m-H\left( a,b \right)+H(n))$$

where a,b – a pair of published subject-object terms, N – set of generated negative samples, H – predictor model, m – margin between positive and negative samples and n – current negative sample.

More detailed information about the model you can find in the original publication (4).
