## Supporting Table 1 for "AI-based mining of biomedical literature: Applications for drug repurposing for the treatment of dementia"

*Supporting Table 1. Summary of pathways and genes for the Dementia class*

| **Pathways** | **Genes** |
| --- | --- |
| Alcoholism | NTRK2, GRIN3B, DDC, MAOA, BDNF, GRIN3A, MAOB, SLC6A3, GRIN2A, PPP1R1B |
| Amphetamine addiction | ARC, GRIN3B, DDC, MAOA, GRIA4, CAMK2A, GRIN3A, MAOB, SLC6A3, GRIN2A, PPP1R1B, GRIA2 |
| cellular response to cocaine | SLC1A1, SLC1A3, SLC1A2, PPP1R1B |
| Cocaine addiction | GRIN3B, DDC, MAOA, GRM3, BDNF, GRIN3A, MAOB, SLC6A3, GRIN2A, PPP1R1B, GRIA2 |
| Common pathways underlying drug addiction | GRM5, GRIA4, CAMK2A, GRIN2A, GRIA2 |
| dopamine catabolic process | COMT, MAOA, MAOB, SLC6A3 |
| dopamine metabolic process | DRD3, DDC, COMT, MAOA, SLC1A1, DAO, MAOB, SLC6A3, GRIN2A |
| Dopamine metabolism | DDC, COMT, MAOA, MAOB |
| Dopaminergic synapse | DRD3, DDC, COMT, MAOA, GRIA4, CAMK2A, MAOB, SLC6A3, GRIN2A, PPP1R1B, GRIA2, GNAL |
| Enzymatic degradation of Dopamine by monoamine oxidase | COMT, MAOA |
| GABA receptor signaling | GABRB2, SLC6A11, SLC6A1, GABRG3, SLC32A1 |
| GABA synthesis release reuptake and degradation | SLC6A11, SLC6A1, SLC32A1, ALDH5A1 |
| GABAergic neuron differentiation | DLX1, GSX2, FEZF2, PRDM13 |
| GABAergic synapse | GABRR3, GABRB2, SLC6A11, SLC6A1, GABRG3, SLC32A1, GLUL |
| GABA-ergic synapse | NLGN3, DRD3, NLGN4Y, DISC1, NRXN3, LRRTM1, SLITRK2, GLRB, MDGA1, NLGN4X, GABRB2, SLC6A11, NBEA, NRXN1, NLGN2, SLC6A1, GABRG3, SLC32A1, SLITRK1, SLC6A17, LRRTM2 |
| Nicotine addiction | GRIN3B, GABRR3, SLC17A7, SLC17A6, GABRB2, GRIA4, GRIN3A, SLC17A8, GRIN2A, GABRG3, SLC32A1, GRIA2 |
| response to cocaine | DRD3, SLC1A1, SLC1A3, SLC6A1, SLC6A3, SLC1A2, PPP1R1B, ST8SIA2 |
| response to external stimulus | DRD3, UNC5D, NTN1, ADNP, NTF3, CACNG2, POU4F3, NEUROG1, UNC5C, GLRA1, NRXN3, LGI1, CNTF, COMT, LMX1A, NTN5, FEZF2, GRM6, SLC1A1, NPAS1, SLC1A3, TBR1, GLRB, PLXNA2, GRIK2, MECP2, KIAA0319, SHANK2, BDNF, GRIN3A, NRXN1, CHD8, EPHA4, MAOB, NRG3, SLC6A3, GRIN2A, CNTNAP2, MAG, TUBB2B, NTRK1, MAP1B, SLC1A2, GRID2, VEGFB, SHANK1, SEMA3A, NTRK3, GLUL |
| Retrograde endocannabinoid signaling | GABRR3, SLC17A7, SLC17A6, GABRB2, GRM5, GRIA4, SLC17A8, GABRG3, SLC32A1, GRIA2 |
| Abnormal nervous system electrophysiology | NLGN3, DRD3, NEUROD2, NTNG2, NTRK2, LGI1, COMT, NTNG1, CHAT, NLGN4X, NEXMIF, GABRB2, MECP2, ASPA, SYNGAP1, MPZ, NBEA, CAMK2A, GALC, SLC6A1, GRIN2A, CNTNAP2, DAOA, NTRK1, MAP1B, SLC1A2, IQSEC2, ACTL6B, LAMC3, ALDH5A1 |
| Abnormal nervous system physiology | DLG3, NLGN3, DRD3, AUTS2, NTN1, ADNP, MYT1L, CACNG2, NEUROD2, NTNG2, IL1RAPL1, GSX2, GLRA1, NTRK2, LGI1, DDC, COMT, ZNF462, DCDC2, MAOA, SLC1A1, PRDM13, SLC1A3, TBR1, NTNG1, GLRB, CHAT, SLC6A9, GRIK2, NLGN4X, ZSWIM6, SHANK3, NEXMIF, GABRB2, SLC6A5, GCDH, MECP2, SPTBN2, ASPA, SYNGAP1, MPZ, POU4F1, GFAP, GRIA4, NBEA, CNGB3, BDNF, CAMK2A, NRXN1, CHD8, DAO, YIF1B, GALC, EPHA4, SLC6A1, PRDM8, RLBP1, SLC6A3, NDNF, GRIN2A, NRTN, CNTNAP2, MAG, DAOA, TUBB2B, GRIP1, FOXP2, NTRK1, MAP1B, SLC1A2, SYN1, GRID2, IQSEC2, SLITRK1, PTCHD1, SLC6A17, NAT8L, ACTL6B, LAMC3, GRIA2, SEMA3A, GNAL, SLC19A3, ALDH5A1, GLUL, |
| Abnormality of central nervous system electrophysiology | NLGN3, DRD3, NEUROD2, NTNG2, NTRK2, LGI1, COMT, NTNG1, CHAT, NLGN4X, NEXMIF, GABRB2, MECP2, ASPA, SYNGAP1, NBEA, CAMK2A, GALC, SLC6A1, GRIN2A, CNTNAP2, DAOA, MAP1B, SLC1A2, IQSEC2, ACTL6B, LAMC3, ALDH5A1 |
| Abnormality of the nervous system | DLG3, NLGN3, DRD3, AUTS2, NTN1, ADNP, MYT1L, CACNG2, NEUROD2, DISC1, NTNG2, IL1RAPL1, GSX2, GLRA1, NTRK2, LGI1, DDC, COMT, ZNF462, DCDC2, MAOA, SLC1A1, PRDM13, SLC1A3, TBR1, NTNG1, GLRB, CHAT, SLC6A9, GRIK2, NLGN4X, ZSWIM6, SHANK3, NEXMIF, GABRB2, SLC6A5, GCDH, MECP2, SPTBN2, ASPA, SYNGAP1, MPZ, POU4F1, GFAP, GRIA4, NBEA, CNGB3, BDNF, CAMK2A, NRXN1, CHD8, DAO, YIF1B, GALC, EPHA4, SLC6A1, PRDM8, RLBP1, SLC6A3, NDNF, GRIN2A, NRTN, CNTNAP2, MAG, DAOA, TUBB2B, GRIP1, FOXP2, NTRK1, MAP1B, SLC1A2, SYN1, GRID2, IQSEC2, SLITRK1, PTCHD1, SLC6A17, NAT8L, ACTL6B, LAMC3, GRIA2, SEMA3A, GNAL, SLC19A3, ALDH5A1, GLUL |
| Activation of NMDA receptors and postsynaptic events | DLG3, GRIN3B, NRGN, GRIA4, NBEA, CAMK2A, GRIN3A, GRIN2A, TUBB2B, GRIA2 |
| adenylate cyclase-inhibiting G protein-coupled receptor signaling pathway | DRD3, GRM4, GRM8, GRM6, GRM3, GRM5, CHRM1, GRIK3 |
| AMPA glutamate receptor complex | DLG3, SHISA6, OLFM3, CACNG2, GRIA4, GRIA2 |
| astrocyte differentiation | HES5, CNTF, HES1, GFAP, EPHA4, MAG, LAMC3, NTRK3 |
| astrocyte projection | GRM3, GRM5, GFAP, SLC17A8, SLC1A2 |
| Astrocytic Glutamate-Glutamine Uptake And Metabolism | SLC1A3, SLC1A2, GLUL |
| asymmetric synapse | NLGN3, DRD3, DLGAP2, SHISA6, NLGN4Y, ARC, CACNG2, DISC1, FXR2, DLGAP3, NTRK2, NETO1, ZDHHC15, SORCS2, SLC1A1, SLC6A9, GRM3, GRIK2, NLGN4X, SHANK3, NRGN, SYNGAP1, GRM5, SHANK2, CAMK2A, GRIN3A, EPHA4, LRRC4, GRIN2A, CHRM1, GRIP1, MAP1B, SYN1, GRID2, SORCS3, LZTS3, SLITRK1, DLGAP1, SHANK1, GRIA2, LRRTM2 |
| asymmetric glutamatergic excitatory synapse | NLGN3, SHISA6, NLGN4Y, NLGN4X |
| Ataxia | NEUROD2, GLRA1, NTRK2, PRDM13, SLC1A3, NTNG1, GLRB, CHAT, GRIK2, ZSWIM6, NEXMIF, GABRB2, SLC6A5, GCDH, MECP2, SPTBN2, SYNGAP1, MPZ, POU4F1, GFAP, GALC, SLC6A1, PRDM8, NDNF, GRIN2A, CNTNAP2, MAG, TUBB2B, SLC1A2, GRID2, NAT8L, ACTL6B, GRIA2, SEMA3A, SLC19A3, ALDH5A1 |
| axon | CPNE6, CPLX3, MBP, AUTS2, ADNP, NTF3, CACNG2, DISC1, NTNG2, IL1RAPL1, FXR2, UNC5C, NTRK2, LGI1, CNTF, LRRTM1, DDC, COMT, NETO1, SLC1A1, GRM3, NRGN, SHANK2, ZNF804A, BDNF, THY1, NRXN1, SLC17A8, EPHA4, SLC6A1, SLC6A3, NRTN, CNTNAP2, MAG, CHRM1, GRIK3, NTRK1, MAP1B, SLC1A2, SYN1, SLC32A1, SYAP1, SEMA3A, NTRK3, GLUL |
| axon development | NLGN3, UNC5D, MBP, AUTS2, NTN1, ADNP, DISC1, POU4F3, NTNG2, UNC5C, NTRK2, NRXN3, LGI1, CNTF, LMX1A, NTN5, FEZF2, SLITRK2, TBR1, NTNG1, PLXNA2, SYNGAP1, KIAA0319, POU4F1, BDNF, THY1, NRXN1, EPHA4, MAG, TUBB2B, NTRK1, MAP1B, SLITRK1, SEMA3A |
| axon guidance | UNC5D, NTN1, POU4F3, UNC5C, NRXN3, LGI1, LMX1A, NTN5, FEZF2, TBR1, PLXNA2, BDNF, NRXN1, EPHA4, TUBB2B, NTRK1, SEMA3A |
| Axon guidance | UNC5D, NTN1, NTNG2, UNC5C, NTNG1, PLXNA2, CAMK2A, EPHA4, LRRC4, SEMA3A |
| axon terminus | CPLX3, NTRK2, SLC1A1, SLC17A8, EPHA4, SLC6A3, CHRM1, GRIK3, SLC32A1, GLUL |
| BDNF activates NTRK2 (TRKB) signaling | NTRK2, BDNF |
| central nervous system development | DRD3, DLX1, MBP, GRIK1, OLIG3, NEUROD2, DISC1, GSX2, UNC5C, HES5, NTRK2, CNTF, LMX1A, FOXN4, SLC17A7, FEZF2, SLC1A1, NPAS1, PRDM13, OLIG1, TBR1, MDGA1, PLXNA2, NLGN4X, ZSWIM6, SHANK3, MECP2, SPTBN2, HES1, ASPA, NRGN, LHX5, POU4F1, GFAP, SLC6A11, SHANK2, NRXN1, CHD8, NLGN2, SLC17A8, CEND1, EPHA4, PRDM8, MAOB, NRG3, SLC6A3, NDNF, GRIN2A, CNTNAP2, MAG, ZIC4, TUBB2B, FOXP2, SLC1A2, SLC32A1, SOX1, GRID2, EMX1, PTCHD1, SLC6A17, ACTL6B, LAMC3, SEMA3A, NTRK3, OLIG2, ALDH5A1 |
| central nervous system neuron differentiation | DLX1, OLIG3, DISC1, GSX2, HES5, NTRK2, LMX1A, FOXN4, FEZF2, TBR1, MDGA1, ZSWIM6, SHANK3, HES1, LHX5, POU4F1, NRXN1, CEND1, EPHA4, NDNF, SOX1, GRID2, EMX1, OLIG2 |
| cognition | NLGN3, DRD3, NLGN4Y, ADNP, ARC, NTF3, NEUROD2, NEUROG1, NTRK2, NRXN3, LMX1A, NETO1, SLC1A1, TBR1, NLGN4X, SHANK3, MECP2, NRGN, SYNGAP1, GRM5, SHANK2, BDNF, NRXN1, SLC6A1, GRIN2A, CNTNAP2, CHRM1, NTRK1, PPP1R1B, SORCS3, PTCHD1, SHANK1 |
| dendrite development | NTN1, ARC, DISC1, IL1RAPL1, DCDC2, NTN5, FEZF2, ZDHHC15, SHANK3, MECP2, SYNGAP1, KIAA0319, CAMK2A, GRIN3A, NLGN2, EPHA4, GRIP1, MAP1B, LZTS3, ACTL6B, SHANK1, SEMA3A |
| dendrite morphogenesis | ARC, IL1RAPL1, DCDC2, ZDHHC15, SHANK3, EPHA4, LZTS3, SHANK1, SEMA3A |
| dendritic spine development | ARC, DISC1, ZDHHC15, SHANK3, CAMK2A, GRIN3A, NLGN2, EPHA4, LZTS3, SHANK1 |
| dendritic spine morphogenesis | ARC, ZDHHC15, SHANK3, EPHA4, LZTS3, SHANK1 |
| Encephalopathy | NEUROD2, NTRK2, SLC6A9, GRIK2, NEXMIF, GABRB2, GCDH, MECP2, SYNGAP1, NRXN1, GALC, SLC6A1, GRIN2A, SLC1A2, ACTL6B, SLC19A3, GLUL |
| excitatory synapse | NLGN3, SHISA6, NLGN4Y, SLC17A7, NETO1, ELFN1, SLC17A6, SLC6A9, NLGN4X, SPTBN2, NRXN1, NLGN2, SLC17A8, LRRC4, GRID2, SHANK1, GRIA2, LRRTM2 |
| excitatory synapse assembly | NRXN3, SHANK3, NRXN1, NLGN2, LRRC4, GRID2, NTRK3, LRRTM2 |
| forebrain generation of neurons | DLX1, DISC1, HES5, FEZF2, TBR1, ZSWIM6, SHANK3, HES1, LHX5, NDNF, SOX1 |
| G protein-coupled glutamate receptor signaling pathway | GRM4, GRM8, GRM6, GRM3, GRM5, GRIK3 |
| generation of neurons | NLGN3, CPNE6, DLX1, UNC5D, MBP, AUTS2, NTN1, ADNP, OLIG3, OLFM3, MYT1L, ARC, NEUROG2, NTF3, NEUROD2, DISC1, POU4F3, NTNG2, IL1RAPL1, GSX2, NEUROG1, ASTN2, UNC5C, HES5, NTRK2, NRXN3, LGI1, CNTF, LMX1A, FOXN4, DCDC2, NTN5, FEZF2, ZDHHC15, PRDM13, OLIG1, SLC1A3, SLITRK2, TBR1, NTNG1, MDGA1, PLXNA2, NLGN4X, ZSWIM6, SHANK3, NEXMIF, GABRB2, MECP2, HES1, CPNE5, SYNGAP1, LHX5, KIAA0319, POU4F1, GFAP, ZNF804A, BDNF, CAMK2A, GRIN3A, THY1, NRXN1, NLGN2, CEND1, EPHA4, NRG3, NDNF, NRTN, CNTNAP2, MAG, TUBB2B, GRIP1, NTRK1, MAP1B, SYN1, SOX1, ST8SIA2, GRID2, EMX1, LZTS3, SLITRK1, ACTL6B, SHANK1, SEMA3A, NTRK3, OLIG2, |
| glial cell development | HES5, NTRK2, CNTF, OLIG1, ASPA, GFAP, PRDM8, MAG, LAMC3 |
| glial cell differentiation | DRD3, DLX1, GSX2, HES5, NTRK2, CNTF, OLIG1, HES1, ASPA, GFAP, EPHA4, PRDM8, MAG, SOX1, EMX1, LAMC3, NTRK3, OLIG2 |
| glial cell projection | SLC1A1, GRM3, GRM5, GFAP, SLC17A8, SLC1A2, GLUL |
| glial cell proliferation | NTN1, SHANK3, MECP2, HES1, GFAP, CHRM1 |
| gliogenesis | DRD3, DLX1, NTN1, DISC1, GSX2, HES5, NTRK2, CNTF, OLIG1, SHANK3, MECP2, HES1, ASPA, GFAP, EPHA4, PRDM8, MAG, CHRM1, SOX1, EMX1, LAMC3, NTRK3, OLIG2 |
| Glutamate Neurotransmitter Release Cycle | SLC17A7, SLC1A1, SLC1A3, SLC1A2, SLC1A6 |
| glutamate receptor activity | GRIK1, GRM4, GRM8, GRIN3B, GRM6, GRM3, GRIK2, GRM5, GRIA4, GRIN3A, GRIN2A, GRIK3, GRID2, GRIA2 |
| glutamate receptor binding | SHISA6, CACNG2, NETO1, SHANK3, SHANK2, CAMK2A, SHANK1 |
| glutamate receptor signaling pathway | GRIK1, GRM4, GRM8, GRIN3B, GRM6, SLC1A1, GRM3, GRIK2, SHANK3, GRM5, GRIA4, GRIN3A, GRIN2A, GRIK3, GRID2, GRIA2 |
| hippocampal mossy fiber to CA3 synapse | CACNG2, SLC6A9, GRIK2, SHANK2, LRRTM2 |
| learning or memory | NLGN3, DRD3, NLGN4Y, ADNP, ARC, NTF3, NEUROD2, NEUROG1, NTRK2, NRXN3, LMX1A, NETO1, SLC1A1, TBR1, NLGN4X, SHANK3, MECP2, NRGN, SYNGAP1, GRM5, SHANK2, BDNF, NRXN1, SLC6A1, GRIN2A, CNTNAP2, NTRK1, PPP1R1B, SORCS3, PTCHD1, SHANK1 |
| memory | ADNP, ARC, NTF3, LMX1A, NETO1, SLC1A1, SHANK3, MECP2, BDNF, SLC6A1, GRIN2A, SORCS3, PTCHD1, SHANK1 |
| negative regulation of glial cell differentiation | DRD3, DLX1, HES5, HES1, NTRK3 |
| negative regulation of nervous system development | DRD3, DLX1, MBP, NTN1, HES5, HES1, SYNGAP1, KIAA0319, THY1, MAG, SEMA3A, NTRK3 |
| negative regulation of neurogenesis | DRD3, DLX1, MBP, NTN1, HES5, HES1, SYNGAP1, KIAA0319, THY1, MAG, SEMA3A, NTRK3 |
| negative regulation of neuron apoptotic process | DLX1, ADNP, NTF3, NTRK2, CNTF, SLC1A1, GRIK2, GABRB2, MECP2, SYNGAP1, POU4F1, BDNF, NDNF, MAG, NTRK1, VEGFB |
| negative regulation of neuron death | DLX1, ADNP, NTF3, NTRK2, CNTF, SLC1A1, GRIK2, GABRB2, MECP2, SYNGAP1, POU4F1, BDNF, NDNF, MAG, NTRK1, VEGFB |
| negative regulation of neuron differentiation | DLX1, HES5, CNTF, LMX1A, FEZF2, HES1, MAG, OLIG2 |
| Neuroinflammation and glutamatergic signaling | GRIK1, ARC, DISC1, GRM4, GRM8, CNTF, GRIN3B, SLC17A7, SLC17A6, SLC1A1, SLC1A3, SLC6A9, GRIK2, GRM5, GFAP, GRIA4, BDNF, CAMK2A, GRIN3A, DAO, GRIN2A, GRIK3, SLC1A2, SLC1A6, GRIA2, GLUL |
| neuron apoptotic process | DLX1, ADNP, NTF3, POU4F3, GRM4, NTRK2, CNTF, SLC1A1, GRIK2, GABRB2, MECP2, SYNGAP1, POU4F1, BDNF, NDNF, MAG, NTRK1, GRID2, VEGFB |
| neuron death | DLX1, ADNP, NTF3, POU4F3, GRM4, NTRK2, CNTF, SLC1A1, GRIK2, GABRB2, MECP2, SYNGAP1, POU4F1, BDNF, NDNF, MAG, NTRK1, ST8SIA2, GRID2, VEGFB |
| neuron differentiation | NLGN3, CPNE6, DLX1, UNC5D, MBP, AUTS2, NTN1, ADNP, OLIG3, OLFM3, MYT1L, ARC, NEUROG2, NTF3, NEUROD2, DISC1, POU4F3, NTNG2, IL1RAPL1, GSX2, NEUROG1, UNC5C, HES5, NTRK2, NRXN3, LGI1, CNTF, LMX1A, FOXN4, DCDC2, NTN5, FEZF2, ZDHHC15, PRDM13, OLIG1, SLC1A3, SLITRK2, TBR1, NTNG1, MDGA1, PLXNA2, NLGN4X, ZSWIM6, SHANK3, GABRB2, MECP2, HES1, CPNE5, SYNGAP1, LHX5, KIAA0319, POU4F1, GFAP, ZNF804A, BDNF, CAMK2A, GRIN3A, THY1, NRXN1, NLGN2, CEND1, EPHA4, NDNF, NRTN, CNTNAP2, MAG, TUBB2B, GRIP1, NTRK1, MAP1B, SYN1, SOX1, ST8SIA2, GRID2, EMX1, LZTS3, SLITRK1, ACTL6B, SHANK1, SEMA3A, NTRK3, OLIG2 |
| Neurotransmitter uptake and metabolism In glial cells | SLC1A3, SLC1A2, GLUL |
| NMDA glutamate receptor activity | GRIN3B, GRIN3A, GRIN2A |
| Activation of Ca-permeable Kainate Receptor | DLG3, GRIK1, GRIK2, GRIK3 |
| Activation of Na-permeable kainate receptors | GRIK1, GRIK2 |
| Brain-derived neurotrophic factor (BDNF) signaling pathway | NTF3, NTRK2, BDNF, CAMK2A, GRIP1, NTRK1, SYN1, GRIA2, NTRK3 |
| main axon | MBP, LGI1, THY1, CNTNAP2, MAG, MAP1B, SLC1A2 |
| nervous system process | DLGAP4, NLGN3, DRD3, DLGAP2, CPLX3, MBP, SHISA6, NLGN4Y, ADNP, ARC, NTF3, CACNG2, NEUROD2, POU4F3, GSX2, NEUROG1, GLRA1, DLGAP3, GRM8, NTRK2, NRXN3, GABRR3, LMX1A, NETO1, DCDC2, BEGAIN, GRM6, SLC1A1, NPAS1, SLC1A3, TBR1, GLRB, GRIK2, NLGN4X, SHANK3, GABRB2, MECP2, NRGN, SYNGAP1, GRM5, POU4F1, SHANK2, CNGB3, BDNF, GRIN3A, NRXN1, CHD8, NLGN2, SLC17A8, SLC6A1, RLBP1, SLC6A3, GRIN2A, CNTNAP2, MAG, CHRM1, GABRG3, NTRK1, PPP1R1B, GRID2, CELF4, SORCS3, PTCHD1, DLGAP1, LAMC3, SHANK1, GNAL |
| neuron to neuron synapse | NLGN3, DRD3, DLGAP2, SHISA6, NLGN4Y, ARC, CACNG2, DISC1, FXR2, DLGAP3, NTRK2, NETO1, ZDHHC15, SORCS2, SLC1A1, SLC6A9, GRM3, GRIK2, NLGN4X, SHANK3, NRGN, SYNGAP1, GRM5, SHANK2, CAMK2A, GRIN3A, NRXN1, NLGN2, EPHA4, LRRC4, GRIN2A, CHRM1, GRIP1, MAP1B, SYN1, GRID2, SORCS3, LZTS3, SLITRK1, DLGAP1, SHANK1, GRIA2, LRRTM2 |
| neuronal cell body | CPNE6, MBP, ADNP, RBFOX3, ARC, NEUROG1, ASTN2, FXR2, UNC5C, GLRA1, GRIN3B, DDC, SORCS2, SLC1A1, SLC1A3, GRIK2, SPTBN2, NRGN, CPNE5, GRIA4, SHANK2, ZNF804A, GRIN3A, THY1, NRXN1, SLC17A8, EPHA4, SLC6A1, SLC6A3, CNTNAP2, GRIP1, GRIK3, NTRK1, MAP1B, PPP1R1B, SYAP1, GRIA2, GLUL |
| Neuronal System | DLGAP4, DLG3, NLGN3, DLGAP2, NLGN4Y, GRIK1, CACNG2, IL1RAPL1, GLRA1, DLGAP3, NRXN3, GRIN3B, LRRTM1, COMT, GABRR3, SLC17A7, MAOA, BEGAIN, SLC1A1, SLC1A3, SLITRK2, GLRB, CHAT, GRIK2, NLGN4X, GABRB2, NRGN, GRM5, GRIA4, SLC6A11, SHANK2, NBEA, CAMK2A, GRIN3A, NRXN1, NLGN2, SLC6A1, SLC6A3, GRIN2A, TUBB2B, GRIP1, GABRG3, GRIK3, SLC1A2, SYN1, SLC32A1, SLITRK1, DLGAP1, SLC1A6, SHANK1, GRIA2, GNAL, NTRK3, LRRTM2, ALDH5A1, GLUL |
| Neurotransmitter disorders | DDC, COMT, MAOA, SLC6A3 |
| perikaryon | CPNE6, RBFOX3, NEUROG1, ASTN2, GLRA1, SORCS2, SLC1A1, CPNE5, SLC17A8, EPHA4, CNTNAP2, GRIP1, GRIK3, MAP1B, SYAP1, GLUL |
| Tryptophan metabolism | ACMSD, DDC, MAOA, GCDH, ASMT, MAOB |
| Tryptophan metabolism | ACMSD, DDC, MAOA, GCDH, ASMT |
