## Supporting Table 2 for "AI-based mining of biomedical literature: Applications for drug repurposing for the treatment of dementia"

*Supporting Table 2. Dementia non-specific genes, highly ranked in other disease groups.*

| **Gene** | **Disease/Probability** | **Associated diseases (Genecards)** |
| --- | --- | --- |
| SORCS1 | Diabetes/0.98156 | Ad6, Alzheimer Disease 6, Late-Onset, Alzheimer's Disease 6, Alzheimer Disease 6, Late Onset, Narcolepsy, Alacrima, Achalasia, And Impaired Intellectual Development Syndrome, Parkinson Disease, Late-Onset |
| VPS13C | Diabetes/0.95229 | Parkinson Disease 23, Autosomal Recessive Early-Onset, Young-Onset Parkinson Disease, Parkinson Disease, Late-Onset, Hypothyroidism, Congenital, Nongoitrous 2, Choreoacanthocytosis, Neuroacanthocytosis, Cohen Syndrome, Dystonia 26 Myoclonic, Parkinson Disease 20, Early-Onset |
| ADAM30 | Diabetes/0.583111 | Inflammatory Bowel Disease 13 |
| IDE | Diabetes/0.278512 | Glucose Intolerance, Type 2 Diabetes Mellitus, Alzheimer Disease 19, Alzheimer Disease 6, Diabetes Mellitus |
| NPHS2 | Diabetes/0.185989 | Nephrotic Syndrome Type 2, Genetic Steroid-Resistant Nephrotic Syndrome, Nephrotic Syndrome, Focal Segmental Glomerulosclerosis, Idiopathic Nephrotic Syndrome, Nail-Patella Syndrome, Frasier Syndrome, Lipoid Nephrosis, Membranous Nephropathy, Denys-Drash Syndrome |
| VPS26A | Diabetes/0.0864211 | Ritscher-Schinzel Syndrome, Parkinson Disease Late-Onset, Hereditary Spastic Paraplegia |
| CDR2L | Arthritis/0.497297, Cancer/0.0651226 | Cerebellar Degeneration, Paraneoplastic Cerebellar Degeneration, Spinocerebellar Ataxia, Autosomal Recessive 13, Fallopian Tube Adenocarcinoma, Small Intestine Neuroendocrine Neoplasm |
| CMAS | Arthritis/0.312393 | Coffin-Siris Syndrome 2, Coffin-Siris Syndrome 4, Palmoplantar Keratoderma and Congenital Alopecia 2, Coffin-Siris Syndrome 3, Coffin-Siris Syndrome 1 |
| WRAP53 | Arthritis/0.284242, Cancer/0.154045 | Dyskeratosis Congenita, Autosomal Recessive 3, Li-Fraumeni Syndrome, Dyskeratosis Congenita, Hepatocellular Carcinoma, Revesz Syndrome, Aplastic Anemia, Dyskeratosis Congenita, Autosomal Dominant 1, Retinal Telangiectasia, Coats Disease |
| USP24 | Arthritis/0.234751, Cancer/0.104463 | Non-Syndromic X-Linked Intellectual Disability 99, Parkinson Disease, Late-Onset, Fanconi Anemia, Complementation Group A |
| CLN5 | Arthritis/0.215399 | Ceroid Lipofuscinosis, Neuronal 5, Neuronal Ceroid Lipofuscinosis, Pontocerebellar Hypoplasia Type 2d, Neuronal Ceroid-Lipofuscinoses Ceroid Lipofuscinosis, Neuronal 3, Ceroid Lipofuscinosis, Neuronal 6a, Ceroid Lipofuscinosis, Neuronal, 8, Northern Epilepsy Variant Toxic Encephalopathy |
| PSIP1 | Arthritis/0.198092 | Interstitial Cystitis, Laryngeal Small Cell Carcinoma, Dermatitis Atopic, Immune Deficiency Disease, Cataract |
| SLC24A2 | Hypertension/0.297357, Heart/0.0698319 | Epilepsy, Familial Temporal Lobe, 5 |
| VEGFB | Heart/0.0387546 | Macular Degeneration Age-Related 1, Kuhnt-Junius Degeneration, Macular Retinal Edema, Retinal Vascular Occlusion, Diabetic Macular Edema |
| NPHP1 | Heart/0.00259942 | Senior-Loken Syndrome 1, Nephronophthisis 1, Joubert Syndrome 4, Joubert Syndrome With Renal Defect , Juvenile Nephronophthisis, Apraxia, Cystic Kidney Disease, Nephronophthisis 2, Ocular Motor Apraxia, Oculomotor Apraxia |
| ATP1A3 | Hypertension/0.0473686 | Dystonia 12, Cerebellar Ataxia, Areflexia, Pes Cavus, Optic Atrophy, And Sensorineural Hearing Loss, Alternating Hemiplegia Of Childhood 2, Developmental And Epileptic Encephalopathy 99, Hemiplegia, Parkinsonism, Familial Hemiplegic Migraine, Quadriplegia, Movement Disease, Hemidystonia |
| SCN2B | Hypertension/0.0436521 | Atrial Fibrillation, Familial 14, Brugada Syndrome, Familial Atrial Fibrillation, Atrial Fibrillation, Right Bundle Branch Block, Sudden Infant Death Syndrome, Generalized Epilepsy With Febrile Seizures Plus, Paroxysmal Extreme Pain Disorder |
| ECE2 | Hypertension/0.0258374 | Currarino Syndrome Periventricular Nodular Heterotopia |
| ZFPM2-AS1 | Cancer/0.254333 | 46 Xy Sex Reversal 9, Diaphragmatic Hernia 3, Double Outlet Right Ventricle, Tetralogy Of Fallot |
| HPCA | Cancer/0.102375 | Dystonia 2 Torsion Autosomal Recessive, Torsion Dystonia 2, Dystonia, Dystonia 25, Torsion Dystonia 4, Dystonia 6 Torsion, Dystonia 27 |
| CASP4 | Cancer/0.102103 | Neuroblastoma, Cowpox, Alzheimer Disease, Familial 1 |
| DRD3 | SUD/0.930068 | Tremor Hereditary Essential 1, Schizophrenia, Essential Tremor, Tremor, Schizoaffective Disorder, Bipolar Disorder, Cocaine Dependence, Opiate Dependence |
| GRM4 | SUD/0.999065 | Epilepsy Myoclonic Juvenile, Epilepsy, Schizophrenia, Childhood Absence , Epilepsy, Fragile X Syndrome |
| GRM8 | SUD/0.997041 | Alcohol Dependence, Epilepsy, Schizophrenia, Autism, Attention Deficit-Hyperactivity Disorder |
| COMT | SUD/0.994641 | Schizophrenia, Panic Disorder 1 , Bardet-Biedl Syndrome , Chromosome 22q11.2 Deletion Syndrome Distal, Schizotypal Personality Disorder, Paranoid Schizophrenia, Psychotic Disorder, Obsessive-Compulsive Disorder |
| ZDHHC15 | SUD/0.99609 | Intellectual Developmental Disorder X-Linked 91, Spastic Diplegia, Non-Syndromic X-Linked Intellectual Disability 91, Non-Syndromic X-Linked Intellectual Disability 98, Non-Syndromic X-Linked Intellectual Disability 58, Cerebral Palsy Ataxic Autosomal Recessive, Tonne-Kalscheuer Syndrome |
| GRM6 | SUD/0.992459 | Night Blindness, Congenital Stationary Type 1b, Congenital Stationary Night Blindness, Leber Plus Disease, Fundus Dystrophy , Night Blindness, Night Blindness Congenital Stationary Type 1a, Retinoschisis 1 X-Linked Juvenile, Abnormal Threshold Of Rods, Cone-Rod Dystrophy X-Linked 3 |
| SLC6A9 | SUD/0.997132 | Glycine Encephalopathy With Normal Serum Glycine, Atypical Glycine Encephalopathy, Glycine Encephalopathy 1, Hyperekplexia 3, Glycine Encephalopathy, Iminoglycinuria, Hyperekplexia, Schizophrenia |
| GABRB2 | SUD/0.998377 | Developmental And Epileptic Encephalopathy 92, Non-Specific Early-Onset Epileptic Encephalopathy, Schizophrenia, Developmental And Epileptic , Encephalopathy 74, Bipolar Disorder, Joubert Syndrome 31, Brugada Syndrome 9, Alcohol Dependence |
| GRM5 | SUD/0.99316 | Fragile X Syndrome, Central Nervous System Disease, Temporal Lobe Epilepsy, Status Epilepticus, Obsessive-Compulsive Disorder |
| GRIA4 | SUD/0.998297 | Neurodevelopmental Disorder With Or Without Seizures And Gait Abnormalities, Non-Specific Syndromic Intellectual Disability, Body Mass Index Quantitative Trait Locus 11, Hyperekplexia 4, Barbiturate Dependence, Schizophrenia, Epilepsy, Microcephaly And Chorioretinopathy 2 |
| GRIN2A | SUD/0.993704 | Epilepsy Focal With Speech Disorder And With Or Without Impaired Intellectual Development, Landau-Kleffner Syndrome, Benign Epilepsy With Centrotemporal Spikes, Continuous Spikes And Waves During Sleep, Rolandic Epilepsy-Speech Dyspraxia Syndrome, Speech Disorder, Focal Epilepsy, High Pressure Neurological Syndrome, Aphasia, Huntington Disease |
| SLC32A1 | SUD/0.995051 | Early Infantile Epileptic Encephalopathy, Generalized Epilepsy With Febrile Seizures Plus, Arthrogryposis Distal Type 2a, Retinitis Pigmentosa 84, Rabies, Hyperekplexia, Epilepsy |
| NAT8L | SUD/0.997742 | N-Acetylaspartate Deficiency, Canavan Disease, Developmental And Epileptic Encephalopathy 39 With Leukodystrophy |
| GRIN3B | SUD/0.986516 | Depersonalization Disorder, Schizophrenia, Dissociative Disorder, West Syndrome |
| MAOA | SUD/0.975405 | Brunner Syndrome, Antisocial Personality Disorder, Conduct Disorder, Social Phobia, Personality Disorder, Borderline Personality Disorder |
| ANKK1 | SUD/0.976147 | Heroin Dependence, Antisocial Personality Disorder, Drug Dependence, Pathological Gambling, Neuroleptic Malignant Syndrome |
| SLC6A5 | SUD/0.95194 | Brittle Cornea Syndrome 2, Hyperekplexia 2, Hyperekplexia 1, Hypertonia, Brown-Vialetto-Van Laere Syndrome 1 |
| RASD2 | SUD/0.953032 | Prostate Calculus, Huntington Disease |
| GABRG3 | SUD/0.96172 | Angelman Syndrome, Asperger Syndrome, Alcohol Dependence, Autism Spectrum Disorder, Autism, Pervasive Developmental Disorder |
| GRIK3 | SUD/0.964928 | Schizophrenia, Depersonalization Disorder, Obsessive-Compulsive Disorder, Neuronitis, Alcohol Dependence |
| ALDH5A1 | SUD/0.931324 | Succinic Semialdehyde Dehydrogenase Deficiency, Fetal Akinesia Deformation Sequence 1, Distal Arthrogryposis, Gamma-Amino Butyric Acid Metabolism Disorder, Canavan Disease, Gaba-Transaminase Deficiency, Alcohol-Related Neurodevelopmental Disorder, Epilepsy |
| ANKS1B | SUD/0.946928 | Osteogenesis Imperfecta Type Ix, Supratentorial Meningioma, Avoidant Personality Disorder |
