## Supporting Table 3 for "AI-based mining of biomedical literature: Applications for drug repurposing for the treatment of dementia"

*Supporting Table 3. Disease specific genes, highly ranked in dementia.*

| **Gene (Diabetes)** | **Probability in Dementia** | **Diseases associated with gene (Genecards)** |
| --- | --- | --- |
| SORCS1 | 0.99876 | Alzheimer Disease 6, Narcolepsy, Alacrima, Achalasia, And Impaired Intellectual Development Syndrome, Parkinson Disease Late-Onset |
| VPS13C | 0.999925 | Parkinson Disease 23 Autosomal Recessive Early-Onset, Young-Onset Parkinson Disease, Parkinson Disease Late-Onset, Hypothyroidism Congenital Nongoitrous 2, Choreoacanthocytosis, Neuroacanthocytosis, Cohen Syndrome, Dystonia 26 Myoclonic, Parkinson Disease 20 Early-Onset |
| ADAM30 | 0.999941 | Inflammatory Bowel Disease 13 |
| WFS1 | 0.0981371 | Wolfram-Like Syndrome, Autosomal Dominant, Cataract 41, Wolfram Syndrome 1, Deafness Autosomal Dominant 6, Wolfram Syndrome, Diabetes Insipidus, Wolfram Syndrome 2, Insulinoma 3-Methylglutaconic Aciduria Type Iii, Deafness Autosomal Dominant 1 With Or Without, Thrombocytopenia |
| MTM1 | 0.0675127 | Myopathy, Centronuclear X-Linked, Respiratory System Disease, Centronuclear Myopathy, Polyhydramnios, Myopathy, Charcot-Marie-Tooth Disease Type 4b1, Charcot-Marie-Tooth Disease Type 4b2, Peliosis Hepatis, Myopathy, Centronuclear 1, Inflammatory Bowel Disease 21 |
| **Gene (Arthritis)** | **Probability in Dementia** | **Diseases associated with gene (Genecards)** |
| CHID1 | 0.875140386 | Not mapped |
| CD33 | 0.626172901 | Childhood T-Cell Acute Lymphoblastic Leukemia, Acute Promyelocytic Leukemia, Myeloid Leukemia |
| SEZ6L2 | 0.615366588 | Schizophrenia 3, Spondylocostal Dysostosis 5, Lung Cancer, Autism, Chromosome 16p11.2 Deletion Syndrome, 220-Kb |
| RABGEF1 | 0.597635 | Smith-Mccort Dysplasia 1, Dyggve-Melchior-Clausen Disease |
| GAB2 | 0.38069732 | Breast Tubular Carcinoma, Noonan Syndrome-Like Disorder With Loose Anagen Hair 1, Adult Infiltrating Astrocytic Neoplasm, Common Peroneal Nerve Lesion, Leukemia, Chronic Myeloid |
| ATG16L2 | 0.110349197 | Not mapped |
| LRCH1 | 0.0786002 | Osteoarthritis, Cardiomyopathy, Familial Hypertrophic 2 |
| NKAP | 0.0723239 | Intellectual Developmental Disorder, X-Linked, Syndromic, Hackmann-Di Donato Type |
| CT83 | 0.0543713 | Lung Cancer |
| MTM1 | 0.067512734 | Myopathy, Centronuclear, X-Linked, Respiratory System Disease, Centronuclear Myopathy, Polyhydramnios, Myopathy |
| **Gene (Heart)** | **Probability in Dementia** | **Diseases associated with gene (Genecards)** |
| SLC24A2 | 0.953244 | Epilepsy, Familial Temporal Lobe 5 |
| ADAMTS19 | 0.0292964 | Discrete Subaortic Stenosis, Winchester Syndrome, Nanophthalmos, Weill-Marchesani Syndrome |
| NEBL | 0.019503 | Dilated Cardiomyopathy |
| SLC7A9 | 0.0193229 | Cystinuria, Ureteral Disease, Urolithiasis, Aminoaciduria, Nephrolithiasis Uric Acid, Hypotonia-Cystinuria Syndrome |
| RNF213 | 0.0113761 | Moyamoya Disease 2, Moyamoya Disease 1, Moyamoya Angiopathy, Hemangioma, Atypical Coarctation Of Aorta, Cerebral Arterial Disease, Lymphoma, Median Arcuate Ligament Syndrome, Hemiplegia |
| **Gene (Hypertension)** | **Probability in Dementia** | **Diseases associated with gene (Genecards)** |
| SLC24A2 | 0.953244 | Epilepsy, Familial Temporal Lobe 5 |
| NEBL | 0.019503 | Dilated Cardiomyopathy |
| GJB6 | 0.0129127 | Clouston Syndrome, Deafness, Autosomal Recessive 1b, Deafness Autosomal Dominant 3b, Deafness Autosomal Recessive 1a, Hidrotic Ectodermal Dysplasia 2 |
| MPV17 | 0.0104575 | Mitochondrial DNA Depletion Syndrome 6, Charcot-Marie-Tooth Disease Axonal Type 2ee, Mitochondrial DNA Depletion Syndrome, Mpv17-Related Mitochondrial DNA Maintenance Defect, Mitochondrial DNA Depletion Syndrome 3, Nephrotic Syndrome, Mitochondrial Metabolism Disease, Axonal Neuropathy, Metabolic Acidosis, Kearns-Sayre Syndrome |
| CCM2 | 0.00923541 | Cerebral Cavernous Malformations 2, Cerebral Cavernous Malformations, Cavernous Hemangioma, Cerebral Cavernous Malformation, Familial, Cerebrocostomandibular Syndrome, Brain Angioma, Cerebral Angioma, Klippel-Trenaunay-Weber Syndrome, Intracranial Cavernous Angioma |
| KLHL3 | 0.00872037 | Pseudohypoaldosteronism Type Iid, Pseudohypoaldosteronism Type Iia, Pseudohypoaldosteronism Type Iie, Cerebral Palsy, Pseudohypoaldosteronism, Metabolic Acidosis, Arthrogryposis Distal Type 3, Renal Tubular Transport Disease, Hypomagnesemia 3 Renal |
| EDNRB | 0.001206827 | Waardenburg Syndrome, Type 4a, Abcd Syndrome, Hirschsprung Disease 2, Waardenburg Syndrome, Type 2e, Rare Genetic Deafness |
| **Gene (Cancer)** | **Probability in Dementia** | **Diseases associated with gene (Genecards)** |
| ZFPM2-AS1 | 0.830551 | 46 Xy Sex Reversal 9, Diaphragmatic Hernia 3, Double Outlet Right Ventricle, Tetralogy Of Fallot |
| ACPT | 0.614927 | Amelogenesis Imperfecta Type Ij, Amelogenesis Imperfecta Type Ie, Testicular Cancer, Amelogenesis Imperfecta, Congenital Bile Acid Synthesis Defect, Xeroderma Pigmentosum Complementation Group C |
| CT83 | 0.0543713 | Lung Cancer |
| MAGEC1 | 0.0507373 | Melanoma, Malignant Anus Melanoma, Uterine Adnexa Cancer, Spermatocytoma |
| **Gene (SUD)** | **Probability in Dementia** | **Diseases associated with gene (Genecards)** |
| SLC17A6 | 0.999996 | Arthrogryposis Distal Type 2a, Rabies, Deafness, Autosomal Dominant 25, Gnathodiaphyseal Dysplasia, Arthrogryposis Distal Type 1a |
| DLG3 | 0.999226 | Intellectual Developmental Disorder X-Linked 90, Non-Syndromic X-Linked Intellectual Disability, Non-Syndromic X-Linked Intellectual Disability 90, Syndromic X-Linked Intellectual Disability, Chromosome 3q29 Deletion Syndrome, Acquired Color Blindness, Bipolar Disorder |
| GLRB | 0.998132 | Hyperekplexia 2, Hyperekplexia, Agoraphobia, Hyperekplexia 1, Hyperekplexia 3, Microcephaly and Chorioretinopathy 2, Panic Disorder |
| SLC6A11 | 0.997841 | Giant Axonal Neuropathy 1, Autosomal Recessive, Epilepsy, Childhood Absence Epilepsy, Iminoglycinuria |
| RASD2 | 0.998037 | Prostate Calculus, Huntington Disease |
| MAOB | 0.996708 | Norrie Disease, Antisocial Personality Disorder, Parkinsonism, Psychotic Disorder, Personality Disorder |
| SLC6A3 | 0.997176 | Parkinsonism-Dystonia 1 Infantile-Onset, Tobacco Addiction, Immunodeficiency 67, Cocaine Abuse, Oppositional Defiant Disorder, Parkinsonism, Cocaine Dependence, Rem Sleep Behavior Disorder |
| GRIN3B | 0.990706 | Depersonalization Disorder, Schizophrenia, Dissociative Disorder, West Syndrome |
| GRM3 | 0.992492 | Bipolar Disorder, Schizophrenia, Status Epilepticus, Epilepsy, Schizophreniform Disorder |
| MAP6 | 0.99104 | Cardia Cancer, Schizophrenia, Schizophrenia 1 |
| EXOSC2 | 0.985321 | Short Stature Hearing Loss Retinitis Pigmentosa And Distinctive Facies, Pontocerebellar Hypoplasia Type 1b, Pontocerebellar Hypoplasia Type 1c, Trichohepatoenteric Syndrome 1, Pontocerebellar Hypoplasia Type 1d, Trichohepatoenteric Syndrome 2 |
| GRIN2A | 0.949816359 | Epilepsy, Focal, With Speech Disorder And With Or Without Impaired Intellectual Development, Landau-Kleffner Syndrome, Benign Epilepsy With Centrotemporal Spikes, Continuous Spikes And Waves During Sleep, Rolandic Epilepsy-Speech Dyspraxia Syndrome |
| COMT | 0.928388671 | Schizophrenia, Panic Disorder 1, Bardet-Biedl Syndrome, Chromosome 22q11.2 Deletion Syndrome, Distal |
| BDNF | 0.910142339 | Bulimia Nervosa, Obsessive-Compulsive Disorder, Wagro Syndrome, Wilms Tumor, Aniridia, Genitourinary Anomalies, And Impaired Intellectual Development Syndrome, Congenital Central Hypoventilation Syndrome |
| HTR6 | 0.795129531 | Schizophrenia, Major Depressive Disorder, Mood Disorder, Meckel Syndrome, Type 1, Alzheimer'S Disease |
| GABRA5 | 0.777580876 | Developmental And Epileptic Encephalopathy 79, Non-Specific Early-Onset Epileptic Encephalopathy |
| CHRNA4 | 0.481328444 | Epilepsy, Nocturnal Frontal Lobe, 1, Autosomal Dominant Nocturnal Frontal Lobe Epilepsy, Tobacco Addiction, Frontotemporal Dementia |
| SIGMAR1 | 0.184678928 | Neuronopathy, Distal Hereditary Motor, Autosomal Recessive 2, Amyotrophic Lateral Sclerosis 16, Juvenile, Juvenile Amyotrophic Lateral Sclerosis |
| FOSB | 0.278620203 | Osteoblastoma, Proliferative Fasciitis, Histiocytoid Hemangioma, Hemangioendothelioma, Malignant Epithelioid Hemangioendothelioma |
| GABRA6 | 0.163389171 | Anxiety, Intraventricular Meningioma, Alcohol Dependence, Panic Disorder, Asperger Syndrome |
| GABBR1 | 0.138156799 | Neurodevelopmental Disorder With Language Delay And Variable Cognitive Abnormalities, Autosomal Dominant Non-Syndromic Intellectual Disability, Neurofibromatosis, Type I |
| GAD1 | 0.011471317 | Developmental And Epileptic Encephalopathy 89, Neurodevelopmental Disorder With Progressive Spasticity And Brain White Matter Abnormalities, Spastic Quadriplegic Cerebral Palsy, Developmental And Epileptic Encephalopathy |
