## Supporting Table 4 for "AI-based mining of biomedical literature: Applications for drug repurposing for the treatment of dementia"

*Supporting Table 4. Drugs for repurposing for dementia treatment*

| Gene name | Class predicted by model | FDA-approved drug name | #interactions  In CT database | Gene/drug connection based on literature | Drug bank, Clinical Trials phase completed |
| --- | --- | --- | --- | --- | --- |
| GABBR1 | SUD | Baclofen | 4 | Baclofen is a GABA-B receptor agonist. General use: Baclofen is prescribed to alleviate muscle stiffness and spasms in the body, providing relief from tightness and cramping associated with various medical conditions such as multiple sclerosis or specific spinal injuries (1). | 4 Alcohol Dependency 4 Sequelae of Stroke 4 Cerebral Palsy (CP) / Excessive crying / Pain 4 Severe Spasticity |
| EDNRB | Hypertension | Bosentan | 21 | Bosentan is a EDNRB blocker. Literature data demonstrate that both Bosentan and selective EDNRA blockers effectively prevent the rise in pulmonary artery pressure induced by hypoxia, aiding in the restoration of oxygen saturation (2). | 4 Pulmonary Arterial Hypertension (PAH) 4 Eisenmenger's Syndrome 4 Type 2 Diabetes Mellitus |
| SLC6A3 | SUD | Bupropion | 84 | Bupropion primarily acts by inhibiting the reuptake of norepinephrine and dopamine. Bupropion, originally an antidepressant, is now also utilized in smoking cessation efforts. It is indicated for adult depression, seasonal affective disorder, and aiding in quitting smoking (3). | 4 Smoking, Cessation 4 Depression 4 Nicotine Dependence 4 Type 2 Diabetes Mellitus 4 Friedreich's Ataxia 4 Alcohol Dependency / Smoking 4 Schizoaffective Disorders / Schizophrenia |
| GABRA5 | SUD | Carbamazepine | 2 | Carbamazepine inhibits voltage-gated sodium channels, reducing neuronal excitability, affecting GABAergic neurotransmission. Carbamazepine is prescribed for the management and treatment of epilepsy, trigeminal neuralgia, as well as acute manic and mixed episodes in individuals with bipolar I disorder (4). | 4 Alcohol Dependency 4 Bipolar 1 Disorder 4 Bronchial Asthma 4 Cocaine Related Disorders / Substance Related Disorders 4 Depression, Bipolar 4 Epilepsy |
| GAD1 | SUD | Clozapine | 7 | Clozapine modulates GABAergic neurotransmission, leading to increased GABAergic activity in certain brain regions, affected by GAD1 function. Clozapine, categorized as an atypical or second-generation antipsychotic medication, is employed in the management of treatment-resistant schizophrenia and for reducing the risk of suicide among individuals diagnosed with schizophrenia (5). | 4 Schizophrenia 4 Cannabis Abuse |
| HTR6 | SUD | Clozapine | 4 | Clozapine has an affinity for various serotonin receptors, including the 5-HT6 receptor(6) | 4 Schizophrenia 4 Cannabis Abuse 4 Panic Disorder |
| SIGMAR1 | SUD | Dextromethorphan | 7 | Dextromethorphan agonist of SIGMAR1.  Literature shows the results of preclinical and clinical impacts of Dextromethorphan. The emphasis is placed on its effects in conditions such as depression, stroke, traumatic brain injury, seizures, pain, methotrexate neurotoxicity, Parkinson's disease, and autism (7). | 4 Type 2 Diabetes Mellitus 4 Atherosclerosis / Inflammation / Smoking 4 Major Depressive Disorder (MDD) 4 Atherosclerosis / Inflammation / Smoking 4 Dementia / Pseudobulbar Affect (PBA) / Stroke / Traumatic Brain Injury (TBI) 3 Huntington's Disease (HD) / Irritability |
| FOSB | SUD | Dronabinol | 13 | Dronabinol plays role in brain's reward and addiction pathways (activation of CB1 receptors). Treating anorexia induced by HIV/AIDS and nausea and vomiting resulting from chemotherapy in patients who have not responded to traditional antiemetic therapies. | 4 Post Traumatic Stress Disorder (PTSD) 4 Chest Pain 4 Multiple Sclerosis / Spasticity 4 Opioids Use / Total Knee Arthroplasty (TKA) 3 Cerebral Palsy (CP) |
| GRIN2A | SUD | Dronabinol | 11 | GRIN2A gene encodes the GluN2A subunit of the NMDA receptor that interacts with THC (active ingredient of Drobadiol) (8) |  |
| SEZ6L2 | Arthritis | Dronabinol | 2 | SEZ6L2 gene involved in neuronal development and function of CNS.  THC, the active ingredient in dronabinol, binds to and activates cannabinoid receptors, which leads to various effects, including altered neurotransmitter release, modulation of neuronal excitability, and changes in perception, mood, and cognition. |  |
| COMT | SUD | entacapone | 18 | Entacapone acts in dopamine metabolic pathway, it is a peripheral catechol-O-methyltransferase inhibitor. Entacapone, a selective and reversible inhibitor of COMT, is prescribed as an adjunctive therapy for Parkinson’s disease in combination with LD and carbidopa (9). | 4 Parkinson's Disease (PD) 2,3 Dependence, Cocaine |
| GABRA6 | SUD | Flumazenil | 6 | Flumazenil is a antagonist at the benzodiazepine-binding site on the GABAA receptor. It is acts in GABAergic neurotransmission pathway (10). | 4 General Anesthesia 2 Substance Related Disorders 2,3 Alcohol Dependency 1 Anxiety 1,2 Parkinson's Disease (PD) |
| BDNF | SUD | Fluoxetine | 28 | Treatment with fluoxetine has the potential to enhance BDNF expression and signaling in specific brain regions, notably the hippocampus, which plays a key role in regulating mood and memory. Fluoxetine(FLX), a Selective Serotonin Reuptake Inhibitor (SSRI) prescribed for major depression, can stimulate BDNF expression in the brain. It is increasingly recognized as a valuable tool for modulating neuronal plasticity across various brain regions, resembling the effects of environmental enrichment (EE). FLX promotes neurogenesis and neuronal turnover in the hippocampus, restores youthful plasticity in the adult rat visual cortex, and enhances recovery from spinal cord injuries (11). | 4 Bipolar Disorder (BD) 4 Major Depressive Disorder (MDD) 4 Alcohol Dependency / Depression 4 Anxiety Disorders / Moods Disorders / Schizophrenia 4 Fibromyalgia 4 Alcohol Dependency / Depression / Suicidal Behaviour |
| COMT | SUD | Levodopa | 18 | COMT inhibition is associated with prolonged effects of levodopa in PD Levodopa, a precursor to dopamine, is employed in the treatment of Parkinson's disease, frequently alongside carbidopa, and in other conditions(targed DRD1 - DRD4) characterized by parkinsonism (12). | 4 Post Traumatic Stress Disorder (PTSD) 4 Advanced Idiopathic Parkinson's Disease 4 Akinesia / Delayed Levodopa Onset / Mobility decreased / Motor Symptoms / Parkinson's Disease (PD) 4 Anhedonia / Depression 4 Aphasia / Stroke 4 Depression 4 Dyskinesia / Parkinson's Disease (PD) 4 Stroke 4 Parkinson's Disease (PD) |
| CHRNA4 | SUD | Mecamylamine | 14 | Mecamylamine, a nicotine antagonist, is utilized for the treatment of moderate to severe essential hypertension and uncomplicated malignant hypertension. The relationship between CHRNA4 and mecamylamine stems from their interaction within the cholinergic neurotransmission pathway mediated by nAChRs. Regarding the functional properties of α4β2-nAChR, distinctive characteristics include heightened sensitivities to antagonism by mecamylamine or DHβE, as well as heightened sensitivities to agonist action of EBDN or nicotine (13). | 3 Alcohol Dependency / Depression 2 Age - Related Macular Degeneration (AMD) 2 Depression / Major Depressive Disorder (MDD) 2 Diabetic Macular Edema (DME) 2 Smoking  1 Autism Disorder / Pervasive Development Disorder |
| NQO2 | SUD | Melatonin | 7 | The antioxidant capacity of melatonin is due to its capacity to inhibit a quinone reductase (NQO2) at high concentration (14). | 4 Postural Instability 4 Post-operative Pain Management 4 Febrile Convulsions 4 Anxiety 4 Bipolar Affective Disorders / Schizoaffective Disorders / Schizophrenia 4 Dementia / Sleep disorders and disturbances 4 Diabetes Mellitus / Matrix Metalloproteinases / Melatonin / Periodontitis 4 Epilepsy, Generalized |
| SLC6A3 | SUD | Methylphenidate | 21 | Methylphenidate (MPH) is a catecholamine reuptake inhibitor; it improves cognitive performance of impaired patients by indirectly increasing extracellular dopamine (DA) levels in both striatal and extrastriatal regions through the blockade of dopamine transporters (DAT), while also blocking noradrenaline transporters (15). | 4 Attention Deficit Hyperactivity Disorder (ADHD) 4 Neurofibromatosis, type 1 (von Recklinghausen's disease) 4 Anxiety Disorders 4 Cocaine Related Disorders / Substance Related Disorders 4 Dementia of the Alzheimer's Type 4 Alzheimer's Disease (AD) / Dementia 4 Depression |
| HTR2C | SUD | Olanzapine | 6 | HTR2C is a subtype of serotonin (5-HT) receptor and Olanzapine is its antagonist. This antipsychotic medication is prescribed for managing schizophrenia and bipolar disorder. Its mechanism involves antagonism at various receptors in the brain, such as dopamine D2 receptors, serotonin 5-HT2A receptors, and histamine H1 receptors (16). | 4 Schizoaffective Disorders / Schizophrenia 4 Diabetes 4 Major Depressive Disorder (MDD) 4 Bipolar Disorder (BD) 4 Alzheimer's Disease (AD) / Dementia / Dementia of the Alzheimer's Type / Senile Dementia, Alzheimer Type 4 Headache 4 Psychosis |
| SIGMAR1 | SUD | Pentazocine | 6 | In vitro studies demonstrate that the sigma receptor ligand (+)-pentazocine inhibits apoptotic cell death of retinal ganglion cells induced by homocysteine and glutamate (17). | 4 Pain 2 Bipolar Disorder (BD) / Mania / Manic Disorder / Manic syndromes / Schizoaffective Disorders |
| GABRB2 | SUD | Propofol | 8 | GABRB2 rs3816596 and GABRA1 rs4263535 polymorphisms are associated with susceptibility to the sedation effect of propofol (18). | 4 Disorders of Gallbladder, Biliary Tract and Pancrease 4 Inflammation 4 Neuromuscular Dystrophy / Scoliosis Idiopathic / Spine Deformity |
| MAOB | SUD | rasagiline | 7 | Rasagiline – a novel MAO B inhibitor in Parkinson’s disease therapy. Rasagiline, an irreversible inhibitor of monoamine oxidase, is prescribed for the symptomatic management of idiopathic Parkinson's disease, both as initial monotherapy and as adjunct therapy to levodopa (19). | 4 Parkinson's Disease (PD) 4 Schizophrenia 2 Amyotrophic Lateral Sclerosis (ALS) 2 Multiple System Atrophy (MSA) |
| MAOB | SUD | Selegiline | 32 | Selegiline (L-deprenyl) is a selective, irreversible inhibitor of monoamine oxidase B (MAO-B), used in the treatment of Parkinson's disease (20). | 4 Major Depressive Disorder (MDD) 4 Parkinson's Disease (PD) 3 Borderline Personality Disorder (BPD) 3 Cocaine Related Disorders 3 Major Depressive Disorder (MDD) 2 Cognitive Dysfunctions / Human Immunodeficiency Virus (HIV) Infections 2 Marijuana Abuse 2 Nicotine Dependence |
| SLC18A2 | SUD | Tetrabenazine | 15 | SLC18A2 encodes VMAT2. Tetrabenazine acts as a reversible inhibitor of VMAT2, inhibitor used for the management of chorea associated with Huntington's Disease (21). | 4 Cerebral Palsy (CP) / Excessive crying / Pain 4 Huntington's Disease (HD) 3 Tardive Dyskinesia (TD) |
| COMT | SUD | Tolcapone | 46 | Tolcapone is a medication that inhibits the enzyme catechol-O-methyl transferase (COMT). It is prescribed as an adjunct to levodopa/carbidopa therapy in the treatment of Parkinson's disease (22). | 4 Parkinson's Disease (PD) 2 Alcohol Abuse / Impulsive Behaviors 2 Frontotemporal Lobar Degeneration (FTLD) 2 Nicotine Dependence 2 Pathological Gambling Disorder |
| CHRM4 | SUD | Tropicamide | 14 | Tropicamide effectively reduced tremulous jaw movements caused by the muscarinic agonist pilocarpine and the dopamine antagonist pimozide. Oral tropicamide has been explored as a potential treatment to alleviate sialorrhea in individuals with Parkinson's Disease (23). | 4 Mydriasis 4 Cataracts 3 Type 1 Diabetes Mellitus / Type 2 Diabetes Mellitus |
| CHRNA4 | SUD | Varenicline | 12 | CHRNA4 rs1044396 is associated with smoking cessation in varenicline therapy. Varenicline primarily acts on α4β2 nAChRs, it indirectly influences the expression and function of CHRNA4. By regulating the activity of α4β2 nAChRs, varenicline modifies the subsequent signaling pathways and gene expression patterns linked to these receptors (24). | 4 Recurrences / Smoking, Cessation / Substance Related Disorders 4 Coronary Heart Disease (CHD) 4 Nicotine Dependence 4 Alcohol Dependency / Nicotine Dependence / Schizoaffective Disorders / Schizophrenia 4 Asthma 4 Bipolar Disorder (BD) / Nicotine Dependence / Schizoaffective Disorders / Schizophrenia 4 Cannabis Dependence / Tobacco Dependence 4 Schizoaffective Disorders / Schizophrenia / Schizophreniform Disorders 3 Human Immunodeficiency Virus (HIV) Infections / Smoking |
| SEZ6L2 | Arthritis | Calcitriol | 2 | SEZ6L2 is a gene linked to neural development. Treatment with calcitriol reduced the severity of experimentally induced autoimmune encephalitis by mitigating inflammation and demyelination in the spinal cord (25). | 4 Friedreich's Ataxia 4 Cancer 4 Chronic Kidney Disease 4 Psoriasis |
| CHID1 | Arthritis | Dexamethasone | 6 | Dexamethasone leads to elevated levels of CHID1 mRNA expression (26). | 4 Arthritis 4 Acute Pain 4 Cancer 4 Ocular Inflammation 4 Opioid Use |
| ATG16L2 | Arthritis | cisplatin | 1 | Cisplatin upregulates multiple autophagy-related genes including ATG16L2 (27). | 4 Esophageal Cancer 3 Gastric Cancer 3 Melanoma |
| CD33 | Arthritis | doxorubicin | 7 | CD33 and doxorubicin both can influence immune responses and inflammatory processes through their respective mechanisms of action. Additionally, doxorubicin is known to induce oxidative stress and DNA damage in cells, leading to apoptosis. | 4 Breast Cancer / Obesity 4 Acute Lymphoblastic Leukemia (ALL) 4 Ovarian Neoplasms |
| CD33 | Arthritis | daunorubicin | 18 | The CD33 gene has relevance in the treatment of acute myeloid leukemia (AML) due to its association with the response to daunorubicin, a chemotherapy medication commonly used in AML therapy. | 4 Acute Lymphobkastic Leukemia 4 Leukemias |
| CD33 | Arthritis | cytarabine | 19 | CD33 gene is expressed on the surface of leukemic cells. Cytarabine is a chemotherapy medication commonly used in the treatment of acute myeloid leukemia. It works by interfering with the DNA synthesis process, ultimately leading to the death of rapidly dividing cells, including leukemic cells. | 4 Acute Myeloid Leukemia 4 Lymphoblastic Lymphoma 4 Meningeal Neoplasms |
| CD33 | Arthritis | Tretinoin | 2 | Tretinoin promptly elicited an inflammatory tumor microenvironment dominated by interferon, marked by heightened infiltration of CD8+ T cells (28) | 4 Acne 4 Childhood Acute Promyelocytic Leukemia |
| GAB2 | Arthritis | Tretinoin | 5 | In the nervous system, Gab2 serves as a crucial intermediary connecting bFGF to the PI3K–AKT pathway (29) |  |
| GAB2 | Arthritis | imatinib | 2 | Imatinib induces unique alterations in the phosphorylation state and interactome of Gab2 (30). | 4 Breast Cancer 4 Chronic Myeloid Leukemia (CML) 4 Gastrointestinal Stromal Tumor (GIST) |
| RABGEF1 | Arthritis | Coumarin (not FDA-approved) | 1 | Coumarin can affect the expression and activity of p53and elevate levels of expression of CDK1 (31). | 4 Chronic Venous Insufficiency (CVI) 1 4 Deep Vein Thrombosis / Pulmonary Embolism |
| RABGEF1 | Arthritis | Hydralazine | 1 | RABGEF1, guanine nucleotide exchange factor (GEF), it activates Rab GTPases by promoting the exchange of GDP for GTP on these small GTP-binding proteins playing an important role in vesicle-mediated transport. While hydralazine primarily acts on blood vessels to lower blood pressure, it may indirectly influence cellular processes like vesicle-mediated transport through its effects on cell signaling and metabolism (32). | 4 Hypertension / Type 2 Diabetes Mellitus 4 Congestive Heart Failure (CHF) 3 Atherosclerosis |
| WFS1 | Diabetes | Fulvestrant | 1 | It was illustrated that some mutations in the WFS1 gene can be associated with systemic inflammation. Fulvestrant inhibits estrogen signaling and it may have anti-inflammatory properties by modulation of immune responses and inflammatory processes (33). | 4 Breast Neoplasms / Metastatic Cancer |
| WFS1 | Diabetes | Dexamethasone | 3 | Dexamethasone is an anti-inflammatory agent, additionally it can affect the regulation of cellular calcium (Ca2+) homeostasis and affect the function of WFS1 gene, involved in the same biological processes (34). | 4 Acute Pain / Analgesia / Upper Extremity Injuries 4 Insulin Secretion 4 Ovarian Cancer 4 Systemic Inflammatory Response Syndrome (SIRS) 4 Hypotension |
| WFS1 | Diabetes | pioglitazone | 2 | Pioglitazone suppresses apoptosis and prevents diabetes. WFS1 gene variations are linked to an increased risk of type 2 diabetes, our results could shed light on the gradual decline of beta cells observed in type 2 diabetes (35). | 4 Diabetes 4 Diabetes Mellitus 4 Ataxia-Telangiectasia (A-T) 4 Coronary Artery Stenosis / Diabetes Mellitus |
| MTM1 | Arthritis | prednisolone | 3 | Prednisolone is effective in controlling inflammation and alleviating symptoms (36). | 4 Rheumatoid Arthritis 4 Crohn's Disease (CD) 4 Diabetes Mellitus 4 Lymphoma 4 Metastatic Prostate Cancer |

8.	O'Donnell B, Meissner H, Gupta V. Dronabinol. StatPearls. Treasure Island (FL)2024.

21.	LiverTox: Clinical and Research Information on Drug-Induced Liver Injury. Bethesda (MD)2012.
