## Supporting Table 5 for "AI-based mining of biomedical literature: Applications for drug repurposing for the treatment of dementia"

*Supporting Table 5. Common genes and their pathways*

| Pathway | Gene intersections |
| --- | --- |
| protein binding | TIMM8A, FBXO22, CHD1, SSRP1, CIZ1, CCDC8, ALG10B, LUC7L2, UTP20, RPGR, EIF4E3, NPHP3, EFCAB6, SPEF2, WDR37, MROH1, NSFL1C, UBE2G1, KATNB1, STARD9, F8A1, LARP1B, CDADC1, TMEM234, SLAIN1, ACBD4, EXO1, UCK1, ATF1, CFAP36, MCM7, PIGH, NACAD, DNAJC30, KLHL18, CHEK1, GRSF1, RPS27A, ZNHIT2, LRRC42, CATSPER2, ASB7, FANCD2, SRGAP2B, SNRNP48, KBTBD4, TUBB2B, KIFC2, PRPF19, YY1AP1, UVSSA, USP47, COPB1, FBXO38, RNF151, SERTAD1, BEX4, OTUD6B, WDR47, FTSJ3, BBS4, CRYBB1, WDR36, CHDH, ASXL2, SDHAF2, SP140L, PYCR3, KPNA3, CIB3, HIC2, PPP2R5D, PSMB2, AGO2, GPR148, ANXA10, NUDT21, BCL2L15, SRSF9, RSPH14, SERINC3, RASSF8, PDZD11, PHLDA1, ENDOD1, AACS, ARL2BP, ZFAND4, ATXN3, NPAS4, DPP9, ARMC8, S100A11, NKD1, MIA2, IGSF10, SPATA6, YAF2, CUEDC1, SRP54, MTFP1, MYADM, SOX3, SVBP, SNX4, CCR7, S100A1, RANGRF, NEU2, NPBWR1, GKN2, CYSLTR2, ASPN, KCNMB1, DIAPH3, CTSZ, ORMDL3, PPP1R3G, FCHO2, TRPM5, GIPC1, TNFSF9, SH3GLB1, IL8RB, SFSWAP, CD59, KCNAB3, XCL2, KCNG4, SLK, RAB2A, DLG3, CSF1R, SLC7A9, TSPAN4, PITPNM2, COL4A2, TARM1, NLGN3, CPNE6, EFEMP2, TSLP, PKLR, ARHGDIA, CCL17, MARCKSL1, PCYOX1, TNFAIP8L2, LIMD2, ACBD7, KIF21A, SEMA4D, RAB27A, SLC9A3R2, AP2B1, SLC9A3, MUC16, CNR2, SERF1A, S100A9, COL13A1, PDPN, RGS13, WASHC5, CD93, CLC, CCL20, DRG2, MYO9A, OTOS, VASN, SEC16B, SOCS3, WFDC12, CD1B, LGI2, SLC4A1AP, CLIC6, S100A5, OXER1, LCN1, MYH9, SLC10A2, MUC13, GJB3, CNST, SEMA6B, NPC1L1, SNAP23, CAPZA1, PAK3, ARHGEF5, LCP1, CXCL16, CNR1, DOCK6, GIMAP5, PREX2, KANK2, PDHA2, ZBTB24, ESPNL, ANKRD36C, SVOPL, COL1A2, ZNF343, RDH5, GYPB, SERPINB9, ESRRA, CHST13, CCDC92, HTR3A, LGALS2, FCGR2A, GNAI2, MITF, ZNF140, AHR, BHLHA9, GRK5, OR2L13, FCER2, ZNF75A, IL18RAP, LRRC2, ARHGAP23, NFKBIB, TAS2R19, DMAC2, IFNA13, TSHB, ERICH2, ZNF649, RFTN2, NPY, IREB2, GALNT15, INTS5, KRT16, MXD3, HEMGN, IL16, POLE, DNAJC22, SELENOF, HELLS, NR1D2, SLC43A3, AIDA, HTRA2, RFC2, PRTN3, SECISBP2, TRIO, MT-CO1, NPTX2, EFS, GLO1, WARS, NES, TGM2, BRD1, CLPP, GADD45B, HEBP2, WDR83OS, FBXL20, HP, SERPING1, SLC6A1, UPRT, PIN1, MOXD1, ZNF580, ALDH3A1, CARS, FAM89B, JRK, APOBEC2, AZI2, APC, TMED5, KRT83, RHBDD3, CD177, NUSAP1, AFMID, PFKM, POTED, TMBIM6, LATS1, GLUL, PTMS, NEIL2, RASSF4, UBFD1, RGPD3, XAGE3, DCP1B, KBTBD11, SETD9, POLDIP3, DRAP1, METTL21C, CDC42SE1, MLLT3, PRPF18, XRCC3, BAZ1B, SKIV2L, BCDIN3D, SRP68, SURF6, THAP7, DZIP1, ZNF593, CDC27, ANKRD16, TRNAU1AP, COX14, UBE2E1, E2F8, CAPRIN1, ETFBKMT, RPL27A, GAMT, TFB1M, MARF1, ING4, RASSF7, RNF186, ATP6V1G2, DYSF, FBXO11, NDUFA12, PPP2R5B, HAGH, MRPS18B, BAZ2A, POC5, NME7, DCAF11, RANBP3, DCAF4, POLR2C, MBD6, ACTR8, TASOR, HEATR1, AGO1, NSMCE3, EIF4ENIF1, UBTFL1, IQCF1, DDB1, BEX5, POMP, VPS50, PIGT, KLHL40, HAUS7, EMC2, USB1, RP2, METTL2A, SUPT3H, CAPZB, ST7, RANBP1, CCNG1, PIH1D2, TPST2, CEP85, POLR3K, C1D, HBS1L, PFDN2, CEP192, AGO4, C2CD3, FAM111B, NDUFA11, DDX25, KIF27, MIEF2, MLLT6, CLPSL2, MORN3, LCORL, ARFGEF2, ERCC4, POLD2, TMX3, ARMCX3, MTG1, RECQL, PSMC1, SPOPL, IPO13, EPB41L4A, PPWD1, SRSF5, EIF5A2, JADE2, KSR2, CSNK2B, CCDC68, RBM3, MCTS1, PAK6, ABI1, PRMT7, HMGB4, DDA1, MAP3K8, DENND2A, UBE3A, ELOVL1, PSMD1, SH3BGRL2, FXR1, MCF2, SMURF2, LYPLA2, BAG1, PDCD6, GOLT1B, SEC61G, PPM1H, PPP1R13L, TSSK1B, ATG16L2, CIAO2B, MOSPD3, ZNF185, MOV10, KHSRP, DAP3, SUGT1, TRIM8, SMYD1, SESN3, SETSIP, DNAJB1, DACT1, NDRG1, BIRC7, ESYT1, TRAP1, SSR2, BOK, SAMM50, ARHGEF10L, RBM46, BCS1L, LHX1, ITPKB, CCDC170, SDK1, HPGDS, DAAM1, PEX5L, EPN3, JPH4, CAV3, LTBP2, ARAP2, GCC2, FERMT2, AKAP6, PTPRT, INKA2, AAGAB, GREM2, SBF2, PCP4, ANGPTL5, FGF16, PROCR, YKT6, CXCL9, ANO3, UNC5B, OMD, RAP1GAP, ECM1, INSYN1, ACKR3, TRIL, GUCY2C, MYOM3, OSBPL9, ANGPTL3, FN1, PAG1, MTMR4, SEC24D, MACO1, OSMR, PLA2G1B, CPLX2, OLFML2A, APLN, RAB8B, ASPM, EPHA1, RSPO3, SLC15A3, AZIN2, FSCN2, TRAF3, MCEMP1, DNM2, MMRN2, CRMP1, TLR3, FLRT3, SLC9A6, EXOC4, SERPINE1, RAB3IP, TRPV2, EPHA5, RTN4R, SSH1, SMAGP, AP1G1, ATP2B2, MFAP4, RYR3, CARD16, INSL5, SPRY2, ANKRD22, CHRNB4, EFNB2, BGN, THBS3, WNT16, FDCSP, IL17RE, RIC8A, TEKT5, IGF1R, CLDN12, WFIKKN1, RAPGEF4, NOSIP, C1QTNF7, HOOK1, SCARF2, CEACAM7, FCN3, SPINK5, PATJ, TMEM230, ITGA1, JAM3, KIF5A, LYG1, RAB33B, ABAT, CACNA1I, CEACAM1, LOXL2, FLRT1, MYO9B, NUS1, DEFB104A, HCN2, IFRD2, CRTAC1, KCNAB2, FADS2, FOXL2, CGB5, CD79B, CGB8, TPT1, KANSL1L, ZNF19, HTR7, SCML1, HOMEZ, SNX32, THOP1, UROD, FBXO33, TRAF2, MOS, CSRNP1, UGT1A9, TNNT2, FBXW12, FMO3, HDC, ADRA2A, DTX2, PRSS53, IKZF4, MT1B, CDKN2C, GABRB3, HBB, MANSC1, C6ORF47, UGT1A1, ZNF607, YJEFN3, GDNF, FKBP1B, PRRC1, C14ORF93, TTC12, PLEKHA6, SULT1B1, ZNF230, DGKD, DUOXA2, TIMP2, AP1S3, NBEAL1, SLC43A1, CASP8, PLCD1, KRT33B, GFI1B, LONRF2, CCNQ, CXXC1, CRTC1, MT-ATP8, PAM, OAT, RETREG2, SLC25A3, CREB3L3, ITPA, PHEX, ECHDC1, ACP2, DNPH1, MAGEB6, PRMT1, NT5C1A, KRT77, ARR3, STT3A, SCYL3, OPRM1, DDO, PHB2, IKBKG, FATE1, PAGE2, CTNNA2, SERPINB6, PRPS1, PRMT5, RNF31, HDAC1, PARP9, ZNF148, STRN4, LMOD1, RPIA, MAGEA1, TOB1, ETF1, ITM2B, TPI1, CUTC, RABGGTB, GPX1, CDIPT, HSP90AA1, GCH1, SPIC, AMBP, KRT81, IRF1, CTF1, ZNF628, BIK, PADI4, MREG, TSTD1, DDX18, LGALS1, HPRT1, C1ORF56, MYRF, OCM, STMN1, GSTA2, KRT34, ENDOU, ADH5, DHX9, AOC2, SCO2, ZNF707, RANBP3L, CDX2, LRRC46, SNRNP25, KLF12, DLEC1, CCND2, ZNF652, KRTAP2-3, CAB39L, GLI4, FAM133A, DEFB121, GRWD1, DHRSX, AVEN, TIMM23, GTF2A1, TTC30A, PHF21A, DOLK, CYB5R1, ABT1, TAF7, ZNF277, RAD50, MRNIP, SFPQ, ZDHHC4, SRPX, PPP4R2, TMEM65, ACTR10, OTUB1, CLN3, COA5, CCNE2, ZMYND15, TSSK4, NOL9, UBE2D4, RPL6, MRTO4, RPS6KC1, RASSF6, MAL2, PSD4, INTS11, ENTPD5, SRSF3, TRIM14, TBC1D16, HDGF, ARID4B, TFAP2E, WWP1, NCOA1, C4ORF46, YWHAE, ALG3, KLHL7, DMAP1, ZNF467, TP53INP2, NCOA5, MAPK15, CHCHD4, SSX4, YWHAG, NEK7, GOLGA2, GNG2, ADAMTS12, VPS26A, TMTC1, UBQLN1, KCNE4, FGFRL1, PRKCH, ABCC9, ASB10, BTN3A3, LMTK2, PLXDC2, ADAMTSL3, GFRAL, LANCL1, CXCL14, GRB7, CDH26, RTP2, RAB37, FXYD6, SLC2A9, SIGLEC8, F2RL3, KCNQ4, SPATA5L1, IRAK4, KIDINS220, FLOT1, NEFH, CHODL, ATP8B1, PXDNL, COL3A1, VWA2, GFRA3, SLCO1B1, LGI4, IRAK1, EFNA2, FGF11, CD47, TXLNA, SEMA3A, ENPP1, LRIG2, SOST, PRKCA, CSN3, ITGA11, SHC3, PRICKLE1, RAB41, BEST3, GPM6A, SLC8A2, TMTC2, RBP4, GNG10, ETS1, ZNF526, PLA2G12A, LIF, HCK, UBE2U, PNMT, TEX28, TMEM154, CORT, DPYS, PLEKHG7, UTY, TBX6, PHYKPL, EPHX1, PUDP, PLA2G4C, RPUSD1, ZNF69, LRRC74A, PYY, EMILIN1, MAGEB4, SULT1C2, NR0B1, SLC6A2, POU1F1, ESRRB, PRH1, LAP3, SCT, SERPINB4, RBM5, ELL, MCHR1, EPGN, PRNP, MAFF, OGG1, TFG, EIF3K, HEG1, KRT23, SUMF1, NDST1, PBX3, TAB2, KLF3, FGF4, DPT, OAS2, SHC2, NRL, ATOH1, HAUS3, KRT5, ALDH5A1, HDAC6, PPP4C, TSPAN33, PIAS1, HIST1H2BE, ANKRD33, ABTB2, ZKSCAN1, RIBC2, CEP44, POLR3F, NAP1L4, PUM1, METTL18, L3MBTL2, ABTB1, TOM1, NAT10, SMARCD2, CEP152, PIP5K1B, CCDC110, HPS4, FASTKD3, RBM15B, PCYT1A, PIMREG, WDR73, ANKRD17, B4GALT7, SNW1, RBMX2, FBXL3, SERTAD2, MRPS12, HHATL, TCERG1, PEF1, CSNK2A3, PPP2R2A, USP25, ZC3H10, PELI1, PRMT8, TFEB, PTPN9, GPR15, CPNE1, NEURL1B, TGFB1I1, AMIGO2, USO1, LRATD1, ARFRP1, DTNB, KCNRG, SYT5, PRDX6, RGS11, PLD1, NCR3, CXCL6, KAZN, MYO1B, PRKCQ, PALLD, SLCO5A1, NOXA1, SV2A, PILRA, LMCD1, PLPPR2, SPRY1, FGF19, CYS1, SLC6A17, DLGAP1, LGALS14, EMP2, ANKS4B, ERBB4, ERLEC1, SLC26A1, METRNL, EDA2R, RASGEF1B, SLC19A1, FAM120B, PRKAR2B, LCN9, C4BPB, CCDC148, ARHGAP27, CEBPA, AGA, STON1, MUC6, FES, C8G, TBX15, C17ORF58, CD34, CYP2S1, ST6GALNAC3, FKBPL, IDH1, CPN1, NFE2L3, ALKBH3, CDK2, TALDO1, SAMD8, ASPG, NR2F6, G6PD, RPS16, ZEB2, NANOG, CHIC2, SST, ACR, PALM, HMCES, OLIG2, PIGA, ACY1, TNNC1, TMEM220, NCBP2AS2, POU2F1, LRRC10B, KLHDC4 |
| catalytic activity | FBXO22, CHD1, ALG10B, RPGR, UBE2G1, CDADC1, EXO1, UCK1, MCM7, PIGH, CHEK1, TOP1MT, TUBB2B, KIFC2, PRPF19, GOT1L1, USP47, RNF151, OTUD6B, FTSJ3, CHDH, PYCR3, SDR42E1, PPP2R5D, AGO2, SDR9C7, FKBP9, ENDOD1, AACS, ATXN3, DPP9, HSDL2, SRP54, AK7, NEU2, CTSZ, KCNAB3, PLA2G4A, SLK, RAB2A, CSF1R, ST3GAL4, PKLR, PKDCC, PCYOX1, KIF21A, RAB27A, NXN, RGS13, CLC, CAPN8, DRG2, SMPD4, BST1, PAK3, PDHA2, AMY1A, FGGY, CYP3A43, RAB40A, RDH5, CHST13, CYP27C1, GNAI2, GRK5, IL18RAP, PLCD3, ECHDC3, IREB2, GALNT15, POLE, SELENOF, HELLS, HTRA2, RFC2, NOCT, PRTN3, TRIO, MT-CO1, GLO1, WARS, TGM2, BRD1, MTHFS, CLPP, METTL7B, HP, UPRT, ABCC10, PIN1, MOXD1, ALDH3A1, AKR1E2, CARS, APOBEC2, PGK2, RHBDD3, AFMID, PFKM, LATS1, GLUL, NDUFB3, NEIL2, ZDHHC18, DCP1B, SETD9, METTL21C, PRPF18, XRCC3, BAZ1B, SKIV2L, BCDIN3D, TUBB1, UBE2E1, ETFBKMT, GAMT, TFB1M, MARF1, ING4, RNF186, ATP6V1G2, FBXO11, NDUFA12, PNPLA7, RPEL1, CYB5R4, HAGH, NME7, POLR2C, HFM1, RMRP, MARS2, USB1, METTL2A, TPST2, POLR3K, HBS1L, CPO, AGO4, FAM111B, DDX25, KIF27, ERCC4, TMX3, MTG1, RECQL, PSMC1, PPWD1, NMNAT2, JADE2, KSR2, CSNK2B, PAK6, PRMT7, MAP3K8, UBE3A, ELOVL1, SMURF2, LYPLA2, PPM1H, TSSK1B, MOV10, TRIM8, SMYD1, SESN3, BIRC7, TRAP1, BCS1L, ITPKB, HPGDS, MBOAT4, PTPRT, YKT6, PDE8B, RAP1GAP, CAPN11, GUCY2C, MTMR4, PLA2G1B, RAB8B, EPHA1, AZIN2, TRAF3, DNM2, PDGFRL, CRMP1, PIP5KL1, EPHA5, SSH1, ATP2B2, CARD16, PLPPR4, NDST2, IGF1R, NOSIP, PGAP1, DPEP3, KIF5A, LYG1, RAB33B, ABAT, LOXL2, MYO9B, NUS1, ZDHHC8, KCNAB2, FADS2, GBP6, WSCD1, LDHC, FMO2, ATP13A5, THOP1, UROD, TRAF2, MOS, GSTT2B, ABCA10, UGT1A9, FMO3, HDC, DTX2, PRSS53, AADACL3, CDKN2C, HBB, UGT1A1, GSTT4, YJEFN3, FKBP1B, UGT2B4, SULT1B1, CASD1, GADL1, DGKD, CASP8, PLCD1, MBOAT1, LONRF2, HSD17B2, MT-ATP8, PAM, OAT, SI, CES2, ITPA, PHEX, ECHDC1, ACP2, DNPH1, PRMT1, NT5C1A, STT3A, SCYL3, DDO, IKBKG, PM20D1, PRPS1, PRMT5, BDH1, RNF31, HDAC1, PARP9, GPX5, RPIA, ETF1, TPI1, RABGGTB, GPX1, CDIPT, PAPOLB, HSP90AA1, GCH1, AMBP, NAGS, PADI4, TSTD1, DDX18, GOT1, HPRT1, MYRF, GSTA2, ENDOU, ADH5, DHX9, AOC2, SCO2, DDX3Y, HECTD4, DHRSX, DOLK, CYB5R1, RAD50, WARS2, ALG11, ZDHHC4, TECRL, OTUB1, TSSK4, PTPN20, NOL9, NEIL3, UBE2D4, RPS6KC1, INTS11, TRMT9B, ENTPD5, TRIM14, ZNFX1, WWP1, NCOA1, GLYATL1, ALG3, RNASET2, B3GALT1, MAP3K6, STK31, MAPK15, CHCHD4, NEK7, NMNAT3, SLC27A6, ADAMTS12, TMTC1, FGFRL1, PRKCH, ABCC9, LMTK2, LANCL1, RAB37, SPATA5L1, IRAK4, ATP8B1, PXDNL, IRAK1, ENPP1, PRKCA, RAB41, TMTC2, DDAH1, GNG10, MOGAT1, PLA2G12A, CYP21A2, HCK, UBE2U, PNMT, RRP22, DPYS, UTY, PHYKPL, EPHX1, PUDP, PLA2G4C, PCYT1B, CYP7A1, RPUSD1, SULT1C2, LAP3, AS3MT, OXSM, OGG1, DPP7, SUMF1, NDST1, PFAS, OAS2, ALDH5A1, HDAC6, PPP4C, PIAS1, POLR3F, METTL18, NAT10, NUDT17, PIP5K1B, PCYT1A, B4GALT7, RBMX2, FBXL3, HHATL, CSNK2A3, DTX4, USP25, PELI1, CARNS1, PRMT8, PTPN9, CPNE1, NEURL1B, ARFRP1, RFNG, PRDX6, RGS11, PLPPR3, PLD1, SBK2, PRKCQ, PLPPR2, A4GNT, ADAMTSL1, ERBB4, CERS5, PRKAR2B, AGA, FES, CYP2S1, ST6GALNAC3, IDH1, CPN1, ALKBH3, CDK2, TALDO1, SAMD8, ASPG, G6PD, FUOM, ACR, HMCES, PIGA, ACY1 |
| biological regulation | FBXO22, SGCD, CHD1, SSRP1, CIZ1, PSMA8, CCDC8, ALG10B, TEX15, UTP20, EIF4E3, NPHP3, NSFL1C, KATNB1, F8A1, LARP1B, SLAIN1, RGL4, ATF1, MCM7, DNAJC30, KLHL18, PCDHGC4, CHEK1, GRSF1, CATSPER2, ASB7, FANCD2, TUBB2B, PRPF19, HOXC12, YY1AP1, SLC25A14, USP47, FBXO38, SERTAD1, BEX4, OTUD6B, WDR47, BBS4, WDR36, ASXL2, SDHAF2, SP140L, HIC2, PPP2R5D, AGO2, GPR148, SDR9C7, NUDT21, BCL2L15, SRSF9, SERINC3, RASSF8, PDZD11, PHLDA1, TMIGD3, AACS, ARL2BP, ATXN3, NPAS4, DPP9, S100A11, NKD1, IGSF10, YAF2, PLET1, MYADM, SOX3, FER1L5, SVBP, SNX4, CCR7, S100A1, RANGRF, NPBWR1, GKN2, CYSLTR2, ASPN, KCNMB1, CTSZ, ORMDL3, PPP1R3G, OR51E2, TRPM5, GIPC1, CYSLTR1, KCNMB3, ZNF311, TNFSF9, SH3GLB1, IL8RB, SFSWAP, CD59, KCNAB3, XCL2, KCNG4, PLA2G4A, SLK, OR2W3, HCRTR1, DLG3, CSF1R, ST3GAL4, PITPNM2, COL4A2, TARM1, NLGN3, CPNE6, EFEMP2, TSLP, ARHGDIA, CCL17, TMEM215, PKDCC, MYMK, MARCKSL1, TNFAIP8L2, SEMA4D, RAB27A, AP2B1, SLC9A3, CNR2, NXN, SERF1A, HRH4, S100A9, PDPN, RGS13, WASHC5, CLC, CCL20, DRG2, MYO9A, POU3F4, VASN, SEC16B, SLC25A27, SOCS3, WFDC12, CD1B, CLIC6, OXER1, ZPBP2, MYH9, GJB3, CNST, SEMA6B, SNAP23, CAPZA1, BST1, PAK3, ARHGEF5, LCP1, CXCL16, CNR1, DOCK6, PREX2, KANK2, ZBTB24, LMBRD2, COL1A2, ZNF343, RAB40A, RDH5, SERPINB9, ESRRA, CYP27C1, CCDC92, OR52H1, HTR3A, LGALS2, FCGR2A, GNAI2, MITF, ZNF140, AHR, BHLHA9, GRK5, OR2L13, FCER2, ZNF75A, IL18RAP, PLCD3, ARHGAP23, NFKBIB, TAS2R19, ST18, EVC, ECHDC3, IFNA13, TSHB, ZNF649, NPY, TRBV12-5, IREB2, INTS5, KRT16, MXD3, HEMGN, IL16, DYTN, HELLS, NR1D2, AIDA, HTRA2, RFC2, NOCT, PRTN3, SECISBP2, TRIO, NPTX2, EFS, GLO1, WARS, NES, TGM2, BRD1, GADD45B, CASZ1, FBXL20, HP, SERPING1, SLC6A1, UPRT, PIN1, ZNF580, FAM89B, JRK, AZI2, APC, RHBDD3, CD177, NUSAP1, SP5, PFKM, TMBIM6, LATS1, GLUL, PTMS, ZDHHC18, RASSF4, MIR520D, MIR488, MIR519B, MIR517A, MIR665, HOXA10-AS, DCP1B, MIR532, ZNF471, MIR99A, MIR320E, SETD9, POLDIP3, DRAP1, METTL21C, CDC42SE1, MLLT3, XRCC3, BAZ1B, SKIV2L, BCDIN3D, INTS1, THAP7, DZIP1, TUBB1, ZNF593, CDC27, TRNAU1AP, UBE2E1, E2F8, CAPRIN1, ETFBKMT, GAMT, MARF1, ING4, RASSF7, RNF186, ATP6V1G2, DYSF, FBXO11, CREB3L2, CYB5R4, PPP2R5B, BAZ2A, ACTR8, TASOR, HEATR1, AGO1, NSMCE3, EIF4ENIF1, UBTFL1, IQCF1, DDB1, BEX5, KLHL40, RP2, SUPT3H, ELP6, CAPZB, ST7, RANBP1, CCNG1, PIH1D2, CEP85, C1D, HBS1L, PFDN2, AGO4, C2CD3, DDX25, MIEF2, MLLT6, HSF5, LCORL, ARFGEF2, ERCC4, ARMCX3, MTG1, PSMC1, SPOPL, NMNAT2, EIF5A2, JADE2, KSR2, CSNK2B, CCDC68, RBM3, MCTS1, PAK6, ABI1, PRMT7, HMGB4, DDA1, IRX1, MAP3K8, UBE3A, PSMD1, FXR1, MCF2, SMURF2, ZNF91, BAG1, PDCD6, GOLT1B, PPM1H, PPP1R13L, TSSK1B, ZIC2, MOV10, KHSRP, DAP3, SUGT1, TRIM8, SMYD1, SESN3, SETSIP, DNAJB1, DACT1, NDRG1, BIRC7, TRAP1, BOK, MIR300, ARHGEF10L, RBM46, HOXA4, LHX1, ITPKB, SDK1, HPGDS, DAAM1, PEX5L, JPH4, CAV3, LTBP2, ARAP2, GCC2, MBOAT4, FERMT2, AKAP6, PTPRT, ASIC3, KCNK9, GREM2, PCP4, FGF16, PROCR, PDE8B, CXCL9, GPR6, ANO3, UNC5B, RAP1GAP, ECM1, INSYN1, ACKR3, TRIL, GUCY2C, ANGPTL3, FN1, PAG1, MTMR4, MACO1, OSMR, PLA2G1B, CPLX2, OLFML2A, APLN, RAB8B, ASPM, EPHA1, RSPO3, SLC15A3, AZIN2, TRAF3, DNM2, MMRN2, PDGFRL, CRMP1, TLR3, FLRT3, SLC9A6, EXOC4, SERPINE1, RAB3IP, TRPV2, PIP5KL1, EPHA5, RTN4R, SSH1, AP1G1, ATP2B2, MFAP4, RYR3, CARD16, INSL5, SPRY2, GPR34, CHRNB4, PLPPR4, GPR171, EFNB2, SLC9A2, WNT16, IL17RE, RIC8A, NDST2, IGF1R, WFIKKN1, RAPGEF4, NOSIP, CEACAM7, PCDH1, OLFML3, FCN3, SPINK5, PATJ, PGAP1, ITGA1, JAM3, KIF5A, RAB33B, ABAT, CACNA1I, CEACAM1, LOXL2, MYO9B, NUS1, ZDHHC8, HCN2, IFRD2, KCNAB2, FOXL2, CGB5, CD79B, OR56A5, CGB8, TPT1, FMO2, ZNF19, HTR7, UCP1, HOMEZ, OR6B3, SNX32, FBXO33, TRAF2, MOS, CSRNP1, UGT1A9, TNNT2, OR2L8, OR11L1, ADRA2A, DTX2, IKZF4, MT1B, CDKN2C, GABRB3, HBB, C2CD4A, UGT1A1, CHRND, ZNF607, YJEFN3, GDNF, FKBP1B, PRRC1, TAS2R31, ZNF215, C14ORF93, ZNF184, UGT2B4, SULT1B1, ZNF230, DGKD, DUOXA2, TIMP2, SLC43A1, CASP8, PLCD1, MBOAT1, GFI1B, HSD17B2, XKR4, SLC17A4, CCNQ, CXXC1, CRTC1, CREB3L3, PHEX, ZNF799, DNPH1, MAGEB6, PRMT1, ARR3, OPRM1, DDO, PHB2, IKBKG, FATE1, CTNNA2, SERPINB6, PRMT5, RNF31, HDAC1, PARP9, ZNF148, LMOD1, BSX, MAGEA1, TOB1, ETF1, ITM2B, GPX1, HSP90AA1, GCH1, SPIC, AMBP, IRF1, CTF1, ZNF628, BIK, TSTD1, LGALS1, GOT1, HPRT1, C1ORF56, MYRF, STMN1, ENDOU, ADH5, DHX9, ARID4A, ZNF707, RANBP3L, CDX2, KLF12, DLEC1, CCND2, ZNF652, MIR577, CAB39L, GLI4, MIR135A1, MIR492, ZNF732, MIR410, DHRSX, AVEN, GTF2A1, PHF21A, YBX2, ABT1, TAF7, ZNF277, RAD50, MRNIP, SFPQ, WARS2, SRPX, PPP4R2, TEX2, TMEM65, OTUB1, CLN3, CCNE2, ZMYND15, TSSK4, MEIOC, RPL6, MRTO4, RPS6KC1, RASSF6, PSD4, INTS11, SRSF3, TRIM14, TBC1D16, ZNFX1, FOXB2, HDGF, ARID4B, TFAP2E, WWP1, NCOA1, YWHAE, DMAP1, ZNF467, TP53INP2, MAP3K6, NCOA5, MAPK15, SSX4, YWHAG, NEK7, GOLGA2, GNG2, ADAMTS12, VPS26A, UBQLN1, KCNE4, FGFRL1, PRKCH, GPR142, ABCC9, ASB10, GPR87, BTN3A3, AHSG, GFRAL, LANCL1, CXCL14, GRB7, FXYD6, SIGLEC8, F2RL3, KCNQ4, IRAK4, KIDINS220, FLOT1, NEFH, CHODL, ATP8B1, COL3A1, CLEC16A, VWA2, GFRA3, SLCO1B1, LGI4, IRAK1, EFNA2, FGF11, CD47, CAPS, SEMA3A, ENPP1, LRIG2, SOST, PRKCA, CSN3, ITGA11, SHC3, PRICKLE1, BEST3, GPM6A, SLC12A1, SLC8A2, RBP4, DDAH1, FFAR1, GNG10, ETS1, LIF, OR2AP1, CYP21A2, MAFA, HCK, RRP22, CORT, PLEKHG7, UTY, TBX6, ZNF536, PLA2G4C, CYP7A1, ZNF69, PYY, EMILIN1, TAS2R60, MAGEB4, NR0B1, SLC6A2, POU1F1, ZNF30, ESRRB, SCT, SERPINB4, RBM5, ELL, PANO1, MCHR1, EPGN, PRNP, MAFF, OGG1, TFG, EIF3K, HEG1, NDST1, PBX3, TAB2, KLF3, FGF4, DPT, OAS2, SHC2, NRL, ATOH1, CAAP1, KRT5, HDAC6, PPP4C, PIAS1, ZKSCAN1, MIR1283-2, POLR3F, PUM1, METTL18, L3MBTL2, TOM1, NAT10, SMARCD2, HPS4, FASTKD3, RBM15B, WDR73, ANKRD17, B4GALT7, SNW1, FBXL3, SERTAD2, HHATL, TCERG1, PEF1, CSNK2A3, DTX4, USP25, ZC3H10, MIR195, PELI1, PRMT8, TFEB, PTPN9, GPR15, CPNE1, NEURL1B, TGFB1I1, AMIGO2, USO1, ARFRP1, DTNB, RFNG, KCNRG, SYT5, PRDX6, RGS11, PLPPR3, PLD1, KCNK12, SBK2, NCR3, CXCL6, HOXB8, PRKCQ, NOXA1, SV2A, PILRA, LMCD1, PLPPR2, SPRY1, FGF19, MUC12, DLGAP1, A4GNT, EMP2, ERBB4, ERLEC1, POPDC2, METRNL, EDA2R, RASGEF1B, TMEM204, SLC22A13, FAM120B, PRKAR2B, C4BPB, ARHGAP27, CEBPA, STON1, FES, C8G, TBX15, FOLR2, CD34, CYP2S1, FKBPL, IDH1, NFE2L3, CDK2, SAMD8, PTGFR, NR2F6, G6PD, ZEB2, NANOG, SST, ACR, PALM, HMCES, OLIG2, TNNC1, POU2F1, MIR671, MIR885 |
| response to stimulus | FBXO22, SGCD, SSRP1, TEX15, RPGR, NPHP3, RGL4, EXO1, ATF1, MCM7, CHEK1, ASB7, FANCD2, TUBB2B, PRPF19, SLC25A14, UVSSA, USP47, FBXO38, WDR47, BBS4, ASXL2, SDHAF2, PPP2R5D, PSMB2, GPR148, SRSF9, SERINC3, RASSF8, TMIGD3, ENDOD1, ARL2BP, ATXN3, NPAS4, S100A11, NKD1, SRP54, MTFP1, PLET1, MYADM, SNX4, CCR7, S100A1, RANGRF, NPBWR1, GKN2, CYSLTR2, ASPN, KCNMB1, OR51E2, TRPM5, GIPC1, CYSLTR1, KCNMB3, TNFSF9, SH3GLB1, IL8RB, CD59, XCL2, PLA2G4A, OR2W3, HCRTR1, CSF1R, ST3GAL4, PITPNM2, COL4A2, TARM1, NLGN3, CPNE6, TSLP, PKLR, ARHGDIA, CCL17, MYMK, TNFAIP8L2, SEMA4D, RAB27A, CNR2, NXN, HRH4, S100A9, PDPN, RGS13, CLC, CCL20, DRG2, MYO9A, VASN, SLC25A27, SOCS3, WFDC12, CD1B, OXER1, LCN1, MYH9, SLC10A2, GJB3, SEMA6B, NPC1L1, SNAP23, SMPD4, BST1, PAK3, ARHGEF5, LCP1, CXCL16, CNR1, DOCK6, PREX2, KANK2, LMBRD2, COL1A2, RAB40A, RDH5, SERPINB9, ESRRA, CCDC92, OR52H1, HTR3A, LGALS2, FCGR2A, GNAI2, MITF, AHR, IGHV3-23, GRK5, OR2L13, IGLC2, FCER2, IL18RAP, PLCD3, ARHGAP23, NFKBIB, TAS2R19, ST18, EVC, ECHDC3, IFNA13, TSHB, RFTN2, NPY, TRBV12-5, KRT16, IL16, POLE, SELENOF, HELLS, NR1D2, AIDA, HTRA2, RFC2, NOCT, PRTN3, TRIO, MT-CO1, NPTX2, EFS, TGM2, BRD1, GADD45B, FBXL20, HP, SERPING1, SLC6A1, UPRT, PIN1, ZNF580, ALDH3A1, FAM89B, JRK, AZI2, APC, RHBDD3, CD177, SP5, TMBIM6, LATS1, GLUL, NEIL2, ZDHHC18, RASSF4, MIR320E, SETD9, METTL21C, CDC42SE1, MLLT3, XRCC3, BAZ1B, SRP68, DZIP1, TUBB1, MARF1, ING4, RASSF7, RNF186, DYSF, NDUFA12, CREB3L2, CYB5R4, PPP2R5B, ACTR8, NSMCE3, DDB1, BEX5, SUPT3H, RANBP1, CCNG1, POLR3K, HBS1L, CEP192, C2CD3, CLPSL2, ARFGEF2, ERCC4, POLD2, SWI5, RECQL, JADE2, KSR2, CSNK2B, CCDC68, MCTS1, PAK6, ABI1, MAP3K8, UBE3A, FXR1, MCF2, SMURF2, BAG1, PDCD6, GOLT1B, TSSK1B, MOV10, KHSRP, DAP3, TRIM8, SESN3, DNAJB1, DACT1, NDRG1, BIRC7, TRAP1, BOK, ARHGEF10L, LHX1, ITPKB, SDK1, HPGDS, DAAM1, PEX5L, CAV3, LTBP2, ARAP2, FERMT2, AKAP6, PTPRT, ASIC3, GREM2, PCP4, FGF16, PROCR, PDE8B, CXCL9, GPR6, ANO3, UNC5B, RAP1GAP, ECM1, INSYN1, ACKR3, TRIL, GUCY2C, ANGPTL3, FN1, PAG1, MTMR4, MACO1, OSMR, PLA2G1B, CPLX2, OLFML2A, APLN, ASPM, EPHA1, RSPO3, SLC15A3, TRAF3, DNM2, MMRN2, PDGFRL, TLR3, FLRT3, SLC9A6, SERPINE1, TRPV2, EPHA5, RTN4R, SSH1, AP1G1, MFAP4, RYR3, CARD16, INSL5, SPRY2, GPR34, CHRNB4, PLPPR4, GPR171, EFNB2, WNT16, IL17RE, RIC8A, IGF1R, WFIKKN1, RAPGEF4, CEACAM7, OLFML3, FCN3, SPINK5, PATJ, ITGA1, JAM3, LYG1, RAB33B, ABAT, CACNA1I, CEACAM1, LOXL2, MYO9B, NUS1, DEFB104A, HCN2, CGB5, CD79B, GBP6, OR56A5, CGB8, IGHV6-1, TPT1, FMO2, HTR7, UCP1, OR6B3, TRAF2, MOS, CSRNP1, UGT1A9, TNNT2, OR2L8, OR11L1, ADRA2A, DTX2, MT1B, GABRB3, HBB, C2CD4A, UGT1A1, IGHV3-48, CHRND, YJEFN3, TRBJ2-3, GDNF, FKBP1B, TRDV2, TAS2R31, TRGV1, SULT1B1, DGKD, DUOXA2, TIMP2, CASP8, PLCD1, HSD17B2, CRTC1, PAM, CREB3L3, CES2, PHEX, PRMT1, ARR3, SCYL3, OPRM1, PHB2, IKBKG, CTNNA2, SERPINB6, PRMT5, RNF31, HDAC1, PARP9, ZNF148, GPX5, MAGEA1, TOB1, GPX1, HSP90AA1, GCH1, AMBP, IRF1, CTF1, BIK, PADI4, LGALS1, GOT1, HPRT1, STMN1, GSTA2, ENDOU, ADH5, DHX9, ARID4A, AOC2, SCO2, IGKV1-27, DLEC1, CCND2, CAB39L, DEFB121, TAF7, ZNF277, RAD50, MRNIP, SFPQ, SRPX, STRC, PPP4R2, TEX2, OTUB1, CLN3, TSSK4, MEIOC, NEIL3, RPS6KC1, RASSF6, PSD4, SRSF3, TRIM14, ZNFX1, HDGF, ARID4B, WWP1, NCOA1, YWHAE, DMAP1, RNASET2, MAP3K6, NCOA5, MAPK15, YWHAG, NEK7, NMNAT3, GNG2, ADAMTS12, UBQLN1, FGFRL1, PRKCH, GPR142, ABCC9, ASB10, GPR87, BTN3A3, AHSG, GFRAL, LANCL1, CXCL14, GRB7, RTP2, SIGLEC8, F2RL3, IRAK4, KIDINS220, FLOT1, NEFH, PXDNL, COL3A1, CLEC16A, VWA2, GFRA3, SLCO1B1, IRAK1, EFNA2, FGF11, CD47, CAPS, SEMA3A, ENPP1, LRIG2, SOST, PRKCA, ITGA11, SHC3, PRICKLE1, GPM6A, SLC8A2, RBP4, DDAH1, FFAR1, GNG10, ETS1, LIF, OR2AP1, MAFA, HCK, IGKV3-20, IGLV3-9, RRP22, CORT, IGLV3-32, PLEKHG7, TBX6, EPHX1, ZNF536, PLA2G4C, CYP7A1, PYY, EMILIN1, TAS2R60, NR0B1, SLC6A2, ESRRB, AS3MT, SCT, SERPINB4, MCHR1, EPGN, PRNP, OGG1, TFG, HEG1, NDST1, PFAS, TAB2, KLF3, FGF4, IGKV1-8, OAS2, SHC2, NRL, ATOH1, CAAP1, KRT5, HDAC6, PPP4C, PIAS1, HIST1H2BE, ABTB2, POLR3F, PUM1, TOM1, SMARCD2, HPS4, ANKRD17, SNW1, FBXL3, PEF1, DTX4, PPP2R2A, USP25, PELI1, TFEB, GPR15, CPNE1, NEURL1B, TGFB1I1, USO1, XPR1, ARFRP1, RFNG, SYT5, PRDX6, RGS11, PLPPR3, PLD1, SBK2, NCR3, CXCL6, PRKCQ, PILRA, LMCD1, PLPPR2, SPRY1, FGF19, DLGAP1, EMP2, ANKS4B, ERBB4, ERLEC1, METRNL, EDA2R, RASGEF1B, TMEM204, SLC22A13, FAM120B, PRKAR2B, IGKV1-37, C4BPB, ARHGAP27, CEBPA, STON1, FES, C8G, FOLR2, CD34, CYP2S1, FKBPL, IDH1, CPN1, NFE2L3, ALKBH3, CDK2, PTGFR, NR2F6, G6PD, RPS16, ZEB2, SST, ACR, PALM, HMCES, PIGA, TNNC1 |
| multicellular organismal process | TIMM8A, SGCD, PSMA8, TEX15, RPGR, NPHP3, SPEF2, WDR37, KATNB1, EXO1, ATF1, DNAJC30, PCDHGC4, CHEK1, GRSF1, CATSPER2, FANCD2, SRGAP2B, TUBB2B, FBXO38, RNF151, WDR47, BBS4, CRYBB1, WDR36, SDHAF2, HIC2, PPP2R5D, AGO2, GPR148, TMIGD3, ATXN3, NPAS4, NKD1, IGSF10, SPATA6, SRP54, MYADM, SOX3, SVBP, SNX4, CCR7, S100A1, RANGRF, ASPN, KCNMB1, CTSZ, ORMDL3, OR51E2, GIPC1, CYSLTR1, KCNMB3, TNFSF9, IL8RB, CD59, XCL2, PLA2G4A, OR2W3, HCRTR1, CSF1R, ST3GAL4, COL4A2, TARM1, NLGN3, CPNE6, EFEMP2, MFAP5, TSLP, TMEM215, PKDCC, MYMK, MARCKSL1, TNFAIP8L2, SEMA4D, RAB27A, AP2B1, CNR2, NXN, SERF1A, S100A9, COL13A1, PDPN, WASHC5, CD93, CLC, MYO9A, POU3F4, OTOS, VASN, SOCS3, LGI2, ZPBP2, LCN1, MYH9, MUC13, GJB3, SEMA6B, NPC1L1, SNAP23, SMPD4, BST1, PAK3, ARHGEF5, LCP1, CNR1, PREX2, KANK2, ESPNL, COL1A2, RDH5, OR52H1, HTR3A, LGALS2, GNAI2, MITF, AHR, OR2L13, FCER2, IL18RAP, PLCD3, TAS2R19, GREB1L, EVC, IFNA13, NPY, IREB2, KRT16, HEMGN, IL16, POLE, SELENOF, HELLS, NR1D2, AIDA, HTRA2, NOCT, SECISBP2, TRIO, MT-CO1, NPTX2, WARS, NES, TGM2, BRD1, CASZ1, FBXL20, SERPING1, SLC6A1, UPRT, PIN1, ZNF580, AZI2, APC, KRT83, RHBDD3, CD177, SP5, LATS1, GLUL, MLLT3, INTS1, DZIP1, TUBB1, E2F8, CAPRIN1, GAMT, MARF1, DYSF, FBXO11, NDUFA12, CREB3L2, PPP2R5B, ACTR8, TASOR, AGO1, EIF4ENIF1, UBTFL1, IQCF1, PIGT, RP2, AGO4, C2CD3, DDX25, KIF27, MLLT6, CLPSL2, ARFGEF2, MTG1, NMNAT2, EIF5A2, JADE2, KSR2, CSNK2B, PAK6, ABI1, IRX1, MAP3K8, UBE3A, FXR1, MCF2, SMURF2, LYPLA2, PDCD6, PPP1R13L, TSSK1B, ZIC2, MOSPD3, MOV10, KHSRP, TRIM8, SMYD1, DNAJB1, DACT1, NDRG1, BIRC7, BOK, RBM46, HOXA4, LHX1, ITPKB, SDK1, HPGDS, JPH4, CAV3, FERMT2, AKAP6, ASIC3, GREM2, SBF2, PCP4, FGF16, PROCR, PDE8B, UNC5B, RAP1GAP, ECM1, INSYN1, ACKR3, TRIL, ANGPTL3, FN1, PAG1, MTMR4, SEC24D, MACO1, PLA2G1B, CPLX2, APLN, RAB8B, ASPM, EPHA1, RSPO3, AZIN2, FSCN2, TRAF3, DNM2, MMRN2, CRMP1, TLR3, FLRT3, SLC9A6, EXOC4, SERPINE1, TRPV2, LUZP1, EPHA5, RTN4R, SSH1, AP1G1, ATP2B2, RYR3, CARD16, INSL5, SPRY2, CHRNB4, PLPPR4, GPR171, EFNB2, BGN, THBS3, WNT16, RIC8A, NDST2, IGF1R, CLDN12, WFIKKN1, HOOK1, PCDH1, SLC26A2, SPINK5, PGAP1, ITGA1, JAM3, KIF5A, RAB33B, ABAT, CACNA1I, CEACAM1, LOXL2, NUS1, ZDHHC8, HCN2, CRTAC1, FOXL2, CGB5, PCDHA8, CD79B, OR56A5, CGB8, TPT1, FMO2, HTR7, UCP1, OR6B3, TRAF2, CSRNP1, UGT1A9, TNNT2, OR2L8, OR11L1, ADRA2A, CDKN2C, GABRB3, HBB, C2CD4A, UGT1A1, CHRND, YJEFN3, GDNF, FKBP1B, TAS2R31, TTC12, DGKD, CASP8, MBOAT1, KRT33B, GFI1B, SPRR2G, HSD17B2, CRTC1, OAT, CFAP97, SI, PHEX, PRMT1, KRT77, ARR3, SCYL3, OPRM1, DDO, PHB2, CTNNA2, SERPINB6, PM20D1, PRPS1, PRMT5, HDAC1, ZNF148, LMOD1, BSX, TOB1, ITM2B, RABGGTB, GPX1, HSP90AA1, GCH1, SPIC, AMBP, KRT81, IRF1, CTF1, ZNF628, BIK, PADI4, LGALS1, HPRT1, MYRF, STMN1, ENDOU, ADH5, DHX9, ARID4A, AOC2, SCO2, DDX3Y, RANBP3L, CDX2, DLEC1, CCND2, YBX2, ABT1, NXNL2, WARS2, STRC, TMEM65, CLN3, ZMYND15, TSSK4, MEIOC, MAL2, ARID4B, WWP1, NCOA1, YWHAE, TP53INP2, YWHAG, ADAMTS12, KCNE4, FGFRL1, PRKCH, ABCC9, BTN3A3, AHSG, GFRAL, CDH26, RTP2, F2RL3, KCNQ4, KIDINS220, FLOT1, NEFH, CHODL, ATP8B1, COL3A1, GFRA3, LGI4, IRAK1, EFNA2, FGF11, CD47, TXLNA, SEMA3A, ENPP1, LRIG2, SOST, PRKCA, ITGA11, SHC3, PRICKLE1, GPM6A, SLC8A2, RBP4, DDAH1, FFAR1, ETS1, LIF, OR2AP1, IGKV3-20, TBX6, ZNF536, PLA2G4C, PCYT1B, PYY, EMILIN1, TAS2R60, NR0B1, SLC6A2, POU1F1, ESRRB, PRH1, SCT, ELL, MCHR1, EPGN, PRNP, MAFF, HEG1, NDST1, PBX3, TAB2, FGF4, OAS2, NRL, ATOH1, KRT5, ALDH5A1, HDAC6, PIAS1, POLR3F, PUM1, SMARCD2, HPS4, ANKRD17, SNW1, PEF1, DTX4, PELI1, TFEB, PTPN9, GPR15, CPNE1, TGFB1I1, AMIGO2, USO1, ARFRP1, DTNB, RFNG, NCR3, CXCL6, KAZN, HOXB8, PRKCQ, PALLD, LMCD1, SPRY1, PCDH12, FGF19, SLC6A17, DLGAP1, EMP2, ADAMTSL1, ERBB4, POPDC2, CERS5, METRNL, SLC19A1, TMEM204, PRKAR2B, C4BPB, CEBPA, MUC6, FES, TBX15, FOLR2, CD34, FKBPL, IDH1, PTGFR, NR2F6, G6PD, ZEB2, NANOG, SST, ACR, PALM, HMCES, OLIG2, TNNC1 |
| regulation of biological process | FBXO22, SGCD, CHD1, SSRP1, CIZ1, PSMA8, CCDC8, ALG10B, TEX15, UTP20, EIF4E3, NPHP3, NSFL1C, KATNB1, F8A1, LARP1B, SLAIN1, RGL4, ATF1, MCM7, DNAJC30, KLHL18, PCDHGC4, CHEK1, GRSF1, CATSPER2, ASB7, FANCD2, TUBB2B, PRPF19, HOXC12, YY1AP1, SLC25A14, USP47, FBXO38, SERTAD1, BEX4, OTUD6B, WDR47, BBS4, WDR36, ASXL2, SDHAF2, SP140L, HIC2, PPP2R5D, AGO2, GPR148, NUDT21, BCL2L15, SRSF9, SERINC3, RASSF8, PDZD11, PHLDA1, TMIGD3, AACS, ARL2BP, ATXN3, NPAS4, DPP9, S100A11, NKD1, IGSF10, YAF2, PLET1, MYADM, SOX3, FER1L5, SVBP, SNX4, CCR7, S100A1, RANGRF, NPBWR1, GKN2, CYSLTR2, ASPN, KCNMB1, CTSZ, ORMDL3, PPP1R3G, OR51E2, TRPM5, GIPC1, CYSLTR1, ZNF311, TNFSF9, SH3GLB1, IL8RB, SFSWAP, CD59, KCNAB3, XCL2, KCNG4, PLA2G4A, SLK, OR2W3, HCRTR1, DLG3, CSF1R, ST3GAL4, PITPNM2, COL4A2, TARM1, NLGN3, CPNE6, EFEMP2, TSLP, ARHGDIA, CCL17, TMEM215, PKDCC, MYMK, MARCKSL1, TNFAIP8L2, SEMA4D, RAB27A, CNR2, NXN, HRH4, S100A9, PDPN, RGS13, WASHC5, CLC, CCL20, DRG2, MYO9A, POU3F4, VASN, SEC16B, SLC25A27, SOCS3, WFDC12, CD1B, CLIC6, OXER1, ZPBP2, MYH9, GJB3, CNST, SEMA6B, SNAP23, CAPZA1, BST1, PAK3, ARHGEF5, LCP1, CXCL16, CNR1, DOCK6, PREX2, KANK2, ZBTB24, LMBRD2, COL1A2, ZNF343, RAB40A, SERPINB9, ESRRA, CCDC92, OR52H1, HTR3A, LGALS2, FCGR2A, GNAI2, MITF, ZNF140, AHR, BHLHA9, GRK5, OR2L13, FCER2, ZNF75A, IL18RAP, PLCD3, ARHGAP23, NFKBIB, TAS2R19, ST18, EVC, ECHDC3, IFNA13, TSHB, ZNF649, NPY, TRBV12-5, IREB2, INTS5, KRT16, MXD3, HEMGN, IL16, DYTN, HELLS, NR1D2, AIDA, HTRA2, RFC2, NOCT, PRTN3, SECISBP2, TRIO, NPTX2, EFS, GLO1, WARS, NES, TGM2, BRD1, GADD45B, CASZ1, FBXL20, HP, SERPING1, SLC6A1, PIN1, ZNF580, FAM89B, JRK, AZI2, APC, RHBDD3, CD177, NUSAP1, SP5, PFKM, TMBIM6, LATS1, GLUL, PTMS, ZDHHC18, RASSF4, MIR520D, MIR488, MIR519B, MIR517A, MIR665, HOXA10-AS, DCP1B, MIR532, ZNF471, MIR99A, MIR320E, SETD9, POLDIP3, DRAP1, METTL21C, CDC42SE1, MLLT3, XRCC3, BAZ1B, SKIV2L, BCDIN3D, INTS1, THAP7, DZIP1, ZNF593, CDC27, TRNAU1AP, UBE2E1, E2F8, CAPRIN1, ETFBKMT, GAMT, MARF1, ING4, RASSF7, RNF186, ATP6V1G2, DYSF, FBXO11, CREB3L2, CYB5R4, PPP2R5B, BAZ2A, ACTR8, TASOR, HEATR1, AGO1, NSMCE3, EIF4ENIF1, UBTFL1, IQCF1, DDB1, BEX5, KLHL40, SUPT3H, ELP6, CAPZB, ST7, RANBP1, CCNG1, CEP85, C1D, HBS1L, PFDN2, AGO4, C2CD3, DDX25, MIEF2, MLLT6, HSF5, LCORL, ARFGEF2, ERCC4, ARMCX3, MTG1, PSMC1, SPOPL, NMNAT2, EIF5A2, JADE2, KSR2, CSNK2B, CCDC68, RBM3, MCTS1, PAK6, ABI1, PRMT7, HMGB4, DDA1, IRX1, MAP3K8, UBE3A, PSMD1, FXR1, MCF2, SMURF2, ZNF91, BAG1, PDCD6, GOLT1B, PPP1R13L, TSSK1B, ZIC2, MOV10, KHSRP, DAP3, TRIM8, SMYD1, SESN3, SETSIP, DNAJB1, DACT1, NDRG1, BIRC7, TRAP1, BOK, MIR300, ARHGEF10L, RBM46, HOXA4, LHX1, ITPKB, SDK1, HPGDS, DAAM1, PEX5L, JPH4, CAV3, LTBP2, ARAP2, GCC2, FERMT2, AKAP6, PTPRT, ASIC3, GREM2, PCP4, FGF16, PROCR, PDE8B, CXCL9, GPR6, UNC5B, RAP1GAP, ECM1, INSYN1, ACKR3, TRIL, GUCY2C, ANGPTL3, FN1, PAG1, MTMR4, MACO1, OSMR, PLA2G1B, CPLX2, OLFML2A, APLN, RAB8B, ASPM, EPHA1, RSPO3, SLC15A3, AZIN2, TRAF3, DNM2, MMRN2, PDGFRL, CRMP1, TLR3, FLRT3, SLC9A6, EXOC4, SERPINE1, RAB3IP, TRPV2, PIP5KL1, EPHA5, RTN4R, SSH1, AP1G1, ATP2B2, MFAP4, RYR3, CARD16, INSL5, SPRY2, GPR34, CHRNB4, PLPPR4, GPR171, EFNB2, WNT16, IL17RE, RIC8A, IGF1R, WFIKKN1, RAPGEF4, CEACAM7, PCDH1, OLFML3, FCN3, SPINK5, PATJ, PGAP1, ITGA1, JAM3, KIF5A, RAB33B, ABAT, CACNA1I, CEACAM1, LOXL2, MYO9B, NUS1, ZDHHC8, HCN2, IFRD2, KCNAB2, FOXL2, CGB5, CD79B, OR56A5, CGB8, TPT1, FMO2, ZNF19, HTR7, UCP1, HOMEZ, OR6B3, SNX32, FBXO33, TRAF2, MOS, CSRNP1, TNNT2, OR2L8, OR11L1, ADRA2A, DTX2, IKZF4, MT1B, CDKN2C, GABRB3, HBB, C2CD4A, UGT1A1, CHRND, ZNF607, YJEFN3, GDNF, FKBP1B, PRRC1, TAS2R31, ZNF215, C14ORF93, ZNF184, ZNF230, DGKD, DUOXA2, TIMP2, SLC43A1, CASP8, PLCD1, MBOAT1, GFI1B, CCNQ, CXXC1, CRTC1, CREB3L3, PHEX, ZNF799, DNPH1, MAGEB6, PRMT1, ARR3, OPRM1, PHB2, IKBKG, FATE1, CTNNA2, SERPINB6, PRMT5, RNF31, HDAC1, PARP9, ZNF148, LMOD1, BSX, MAGEA1, TOB1, ETF1, ITM2B, GPX1, HSP90AA1, GCH1, SPIC, AMBP, IRF1, CTF1, ZNF628, BIK, TSTD1, LGALS1, GOT1, HPRT1, C1ORF56, MYRF, STMN1, ENDOU, DHX9, ARID4A, ZNF707, RANBP3L, CDX2, KLF12, DLEC1, CCND2, ZNF652, MIR577, CAB39L, GLI4, MIR135A1, MIR492, ZNF732, MIR410, DHRSX, AVEN, GTF2A1, PHF21A, YBX2, ABT1, TAF7, ZNF277, RAD50, MRNIP, SFPQ, WARS2, SRPX, PPP4R2, TEX2, TMEM65, OTUB1, CLN3, CCNE2, ZMYND15, TSSK4, MEIOC, RPL6, MRTO4, RPS6KC1, RASSF6, PSD4, INTS11, SRSF3, TRIM14, TBC1D16, ZNFX1, FOXB2, HDGF, ARID4B, TFAP2E, WWP1, NCOA1, YWHAE, DMAP1, ZNF467, TP53INP2, MAP3K6, NCOA5, MAPK15, SSX4, YWHAG, NEK7, GOLGA2, GNG2, ADAMTS12, VPS26A, UBQLN1, KCNE4, FGFRL1, PRKCH, GPR142, ABCC9, ASB10, GPR87, BTN3A3, AHSG, GFRAL, LANCL1, CXCL14, GRB7, FXYD6, SIGLEC8, F2RL3, KCNQ4, IRAK4, KIDINS220, FLOT1, NEFH, CHODL, ATP8B1, COL3A1, CLEC16A, VWA2, GFRA3, LGI4, IRAK1, EFNA2, FGF11, CD47, CAPS, SEMA3A, ENPP1, LRIG2, SOST, PRKCA, ITGA11, SHC3, PRICKLE1, BEST3, GPM6A, SLC8A2, RBP4, DDAH1, FFAR1, GNG10, ETS1, LIF, OR2AP1, MAFA, HCK, RRP22, CORT, PLEKHG7, UTY, TBX6, ZNF536, PLA2G4C, CYP7A1, ZNF69, PYY, EMILIN1, TAS2R60, MAGEB4, NR0B1, SLC6A2, POU1F1, ZNF30, ESRRB, SCT, SERPINB4, RBM5, ELL, PANO1, MCHR1, EPGN, PRNP, MAFF, OGG1, TFG, EIF3K, HEG1, NDST1, PBX3, TAB2, KLF3, FGF4, DPT, OAS2, SHC2, NRL, ATOH1, CAAP1, KRT5, HDAC6, PPP4C, PIAS1, ZKSCAN1, MIR1283-2, POLR3F, PUM1, METTL18, L3MBTL2, TOM1, NAT10, SMARCD2, HPS4, FASTKD3, RBM15B, WDR73, ANKRD17, B4GALT7, SNW1, FBXL3, SERTAD2, HHATL, TCERG1, PEF1, CSNK2A3, DTX4, USP25, ZC3H10, MIR195, PELI1, TFEB, PTPN9, GPR15, CPNE1, NEURL1B, TGFB1I1, AMIGO2, USO1, ARFRP1, DTNB, RFNG, KCNRG, SYT5, PRDX6, RGS11, PLPPR3, PLD1, KCNK12, SBK2, NCR3, CXCL6, HOXB8, PRKCQ, NOXA1, SV2A, PILRA, LMCD1, PLPPR2, SPRY1, FGF19, MUC12, DLGAP1, A4GNT, EMP2, ERBB4, ERLEC1, POPDC2, METRNL, EDA2R, RASGEF1B, TMEM204, SLC22A13, FAM120B, PRKAR2B, C4BPB, ARHGAP27, CEBPA, STON1, FES, C8G, TBX15, FOLR2, CD34, FKBPL, IDH1, NFE2L3, CDK2, SAMD8, PTGFR, NR2F6, G6PD, ZEB2, NANOG, SST, PALM, HMCES, OLIG2, TNNC1, POU2F1, MIR671, MIR885 |
| regulation of cellular process | FBXO22, SGCD, CHD1, SSRP1, CIZ1, PSMA8, CCDC8, ALG10B, TEX15, UTP20, EIF4E3, NPHP3, NSFL1C, KATNB1, LARP1B, SLAIN1, RGL4, ATF1, MCM7, DNAJC30, KLHL18, PCDHGC4, CHEK1, GRSF1, CATSPER2, ASB7, FANCD2, TUBB2B, PRPF19, HOXC12, YY1AP1, SLC25A14, USP47, FBXO38, SERTAD1, BEX4, OTUD6B, WDR47, BBS4, WDR36, ASXL2, SDHAF2, SP140L, HIC2, PPP2R5D, AGO2, GPR148, NUDT21, BCL2L15, SRSF9, SERINC3, RASSF8, PHLDA1, TMIGD3, AACS, ARL2BP, ATXN3, NPAS4, DPP9, S100A11, NKD1, IGSF10, YAF2, PLET1, MYADM, SOX3, FER1L5, SVBP, SNX4, CCR7, S100A1, RANGRF, NPBWR1, GKN2, CYSLTR2, ASPN, KCNMB1, CTSZ, ORMDL3, PPP1R3G, OR51E2, TRPM5, GIPC1, CYSLTR1, ZNF311, TNFSF9, SH3GLB1, IL8RB, SFSWAP, CD59, KCNAB3, XCL2, KCNG4, PLA2G4A, SLK, OR2W3, HCRTR1, DLG3, CSF1R, ST3GAL4, PITPNM2, COL4A2, TARM1, NLGN3, CPNE6, EFEMP2, TSLP, ARHGDIA, CCL17, TMEM215, PKDCC, MARCKSL1, TNFAIP8L2, SEMA4D, RAB27A, CNR2, NXN, HRH4, S100A9, PDPN, RGS13, WASHC5, CLC, CCL20, DRG2, MYO9A, POU3F4, VASN, SEC16B, SLC25A27, SOCS3, CD1B, CLIC6, OXER1, ZPBP2, MYH9, CNST, SEMA6B, CAPZA1, BST1, PAK3, ARHGEF5, LCP1, CXCL16, CNR1, DOCK6, PREX2, KANK2, ZBTB24, LMBRD2, COL1A2, ZNF343, RAB40A, SERPINB9, ESRRA, OR52H1, HTR3A, LGALS2, FCGR2A, GNAI2, MITF, ZNF140, AHR, BHLHA9, GRK5, OR2L13, FCER2, ZNF75A, IL18RAP, PLCD3, ARHGAP23, NFKBIB, TAS2R19, ST18, EVC, ECHDC3, IFNA13, TSHB, ZNF649, NPY, TRBV12-5, IREB2, INTS5, KRT16, MXD3, HEMGN, IL16, HELLS, NR1D2, AIDA, HTRA2, RFC2, NOCT, PRTN3, SECISBP2, TRIO, NPTX2, EFS, GLO1, WARS, NES, TGM2, BRD1, GADD45B, CASZ1, FBXL20, HP, SLC6A1, PIN1, ZNF580, FAM89B, JRK, AZI2, APC, RHBDD3, CD177, NUSAP1, SP5, PFKM, TMBIM6, LATS1, GLUL, PTMS, RASSF4, MIR520D, MIR488, MIR519B, MIR517A, MIR665, HOXA10-AS, DCP1B, MIR532, ZNF471, MIR99A, MIR320E, SETD9, POLDIP3, DRAP1, METTL21C, CDC42SE1, MLLT3, XRCC3, BAZ1B, SKIV2L, BCDIN3D, INTS1, THAP7, DZIP1, ZNF593, CDC27, TRNAU1AP, UBE2E1, E2F8, CAPRIN1, ETFBKMT, MARF1, ING4, RASSF7, RNF186, ATP6V1G2, DYSF, FBXO11, CREB3L2, PPP2R5B, BAZ2A, ACTR8, TASOR, HEATR1, AGO1, NSMCE3, EIF4ENIF1, UBTFL1, IQCF1, DDB1, BEX5, SUPT3H, ELP6, CAPZB, ST7, RANBP1, CCNG1, CEP85, C1D, HBS1L, PFDN2, AGO4, C2CD3, DDX25, MIEF2, MLLT6, HSF5, LCORL, ARFGEF2, ERCC4, ARMCX3, MTG1, NMNAT2, EIF5A2, JADE2, KSR2, CSNK2B, CCDC68, RBM3, MCTS1, PAK6, ABI1, PRMT7, HMGB4, IRX1, MAP3K8, UBE3A, FXR1, MCF2, SMURF2, ZNF91, BAG1, PDCD6, GOLT1B, PPP1R13L, TSSK1B, ZIC2, MOV10, KHSRP, DAP3, TRIM8, SMYD1, SESN3, SETSIP, DNAJB1, DACT1, NDRG1, BIRC7, TRAP1, BOK, MIR300, ARHGEF10L, RBM46, HOXA4, LHX1, ITPKB, HPGDS, DAAM1, PEX5L, JPH4, CAV3, LTBP2, ARAP2, GCC2, FERMT2, AKAP6, PTPRT, ASIC3, GREM2, PCP4, FGF16, PDE8B, CXCL9, GPR6, UNC5B, RAP1GAP, ECM1, INSYN1, ACKR3, TRIL, GUCY2C, ANGPTL3, FN1, PAG1, MTMR4, MACO1, OSMR, PLA2G1B, CPLX2, OLFML2A, APLN, RAB8B, ASPM, EPHA1, RSPO3, SLC15A3, AZIN2, TRAF3, DNM2, MMRN2, PDGFRL, CRMP1, TLR3, FLRT3, SLC9A6, EXOC4, SERPINE1, RAB3IP, TRPV2, PIP5KL1, EPHA5, RTN4R, SSH1, AP1G1, RYR3, CARD16, INSL5, SPRY2, GPR34, CHRNB4, PLPPR4, GPR171, EFNB2, WNT16, IL17RE, RIC8A, IGF1R, WFIKKN1, RAPGEF4, CEACAM7, OLFML3, FCN3, SPINK5, PATJ, PGAP1, ITGA1, JAM3, RAB33B, ABAT, CACNA1I, CEACAM1, LOXL2, MYO9B, NUS1, ZDHHC8, HCN2, IFRD2, KCNAB2, FOXL2, CGB5, CD79B, OR56A5, CGB8, TPT1, FMO2, ZNF19, HTR7, UCP1, HOMEZ, OR6B3, SNX32, TRAF2, MOS, CSRNP1, OR2L8, OR11L1, ADRA2A, DTX2, IKZF4, CDKN2C, GABRB3, HBB, C2CD4A, UGT1A1, CHRND, ZNF607, YJEFN3, GDNF, FKBP1B, PRRC1, TAS2R31, ZNF215, C14ORF93, ZNF184, ZNF230, DGKD, DUOXA2, TIMP2, SLC43A1, CASP8, PLCD1, MBOAT1, GFI1B, CCNQ, CXXC1, CRTC1, CREB3L3, ZNF799, DNPH1, MAGEB6, PRMT1, ARR3, OPRM1, PHB2, IKBKG, FATE1, CTNNA2, PRMT5, RNF31, HDAC1, PARP9, ZNF148, LMOD1, BSX, MAGEA1, TOB1, ETF1, ITM2B, GPX1, HSP90AA1, GCH1, SPIC, AMBP, IRF1, CTF1, ZNF628, BIK, TSTD1, LGALS1, GOT1, HPRT1, C1ORF56, MYRF, STMN1, ENDOU, DHX9, ARID4A, ZNF707, RANBP3L, CDX2, KLF12, DLEC1, CCND2, ZNF652, MIR577, CAB39L, GLI4, MIR135A1, MIR492, ZNF732, MIR410, DHRSX, AVEN, GTF2A1, PHF21A, YBX2, ABT1, TAF7, ZNF277, RAD50, MRNIP, SFPQ, SRPX, PPP4R2, TEX2, OTUB1, CLN3, CCNE2, ZMYND15, TSSK4, MEIOC, RPL6, MRTO4, RPS6KC1, RASSF6, PSD4, INTS11, SRSF3, TRIM14, ZNFX1, FOXB2, HDGF, ARID4B, TFAP2E, WWP1, NCOA1, YWHAE, DMAP1, ZNF467, TP53INP2, MAP3K6, NCOA5, MAPK15, SSX4, YWHAG, NEK7, GOLGA2, GNG2, ADAMTS12, VPS26A, UBQLN1, KCNE4, FGFRL1, PRKCH, GPR142, ABCC9, ASB10, GPR87, BTN3A3, AHSG, GFRAL, LANCL1, CXCL14, GRB7, FXYD6, SIGLEC8, F2RL3, KCNQ4, IRAK4, KIDINS220, FLOT1, NEFH, CHODL, ATP8B1, COL3A1, CLEC16A, VWA2, GFRA3, LGI4, IRAK1, EFNA2, FGF11, CD47, CAPS, SEMA3A, ENPP1, LRIG2, SOST, PRKCA, ITGA11, SHC3, PRICKLE1, GPM6A, SLC8A2, RBP4, DDAH1, FFAR1, GNG10, ETS1, LIF, OR2AP1, MAFA, HCK, RRP22, CORT, PLEKHG7, UTY, TBX6, ZNF536, PLA2G4C, CYP7A1, ZNF69, PYY, EMILIN1, TAS2R60, MAGEB4, NR0B1, POU1F1, ZNF30, ESRRB, SCT, SERPINB4, RBM5, ELL, PANO1, MCHR1, EPGN, PRNP, MAFF, OGG1, TFG, EIF3K, HEG1, NDST1, PBX3, TAB2, KLF3, FGF4, DPT, OAS2, SHC2, NRL, ATOH1, CAAP1, KRT5, HDAC6, PPP4C, PIAS1, ZKSCAN1, MIR1283-2, POLR3F, PUM1, METTL18, L3MBTL2, TOM1, NAT10, SMARCD2, HPS4, FASTKD3, RBM15B, WDR73, ANKRD17, B4GALT7, SNW1, SERTAD2, HHATL, TCERG1, CSNK2A3, DTX4, USP25, ZC3H10, MIR195, PELI1, TFEB, PTPN9, GPR15, CPNE1, NEURL1B, TGFB1I1, AMIGO2, USO1, ARFRP1, RFNG, KCNRG, SYT5, PRDX6, RGS11, PLPPR3, PLD1, KCNK12, SBK2, NCR3, CXCL6, HOXB8, PRKCQ, NOXA1, PILRA, LMCD1, PLPPR2, SPRY1, FGF19, MUC12, DLGAP1, A4GNT, EMP2, ERBB4, ERLEC1, METRNL, EDA2R, RASGEF1B, TMEM204, SLC22A13, FAM120B, PRKAR2B, C4BPB, ARHGAP27, CEBPA, STON1, FES, TBX15, FOLR2, CD34, IDH1, NFE2L3, CDK2, SAMD8, PTGFR, NR2F6, G6PD, ZEB2, NANOG, SST, PALM, HMCES, OLIG2, TNNC1, POU2F1, MIR671, MIR885 |
| organonitrogen compound metabolic process | FBXO22, PSMA8, ALG10B, RPGR, EIF4E3, NSFL1C, UBE2G1, F8A1, LARP1B, CDADC1, UCK1, PIGH, DNAJC30, KLHL18, CHEK1, RPS27A, ASB7, PRPF19, GOT1L1, UVSSA, USP47, FBXO38, RNF151, SERTAD1, BEX4, OTUD6B, CHDH, SDHAF2, PYCR3, PPP2R5D, PSMB2, AGO2, FKBP9, SERINC3, PHLDA1, ARL2BP, ATXN3, DPP9, ARMC8, NKD1, MYADM, SVBP, CCR7, NEU2, CTSZ, ORMDL3, GIPC1, PLA2G4A, SLK, DLG3, CSF1R, ST3GAL4, TSLP, PKDCC, PCYOX1, SEMA4D, SLC9A3R2, NXN, SERF1A, S100A9, CAPN8, DRG2, SOCS3, WFDC12, ZPBP2, LCN1, MYH9, NPC1L1, SMPD4, BST1, PAK3, ARHGEF5, PDHA2, RAB40A, SERPINB9, CHST13, GNAI2, GRK5, ST18, DMAC2, IFNA13, IREB2, GALNT15, SELENOF, AIDA, HTRA2, NOCT, PRTN3, SECISBP2, TRIO, WARS, TGM2, MTHFS, CLPP, GADD45B, WDR83OS, FBXL20, HP, SERPING1, UPRT, PIN1, MOXD1, CARS, APOBEC2, APC, RHBDD3, AFMID, LATS1, GLUL, NDUFB3, NEIL2, ZDHHC18, DCP1B, POLDIP3, METTL21C, BAZ1B, SKIV2L, INTS1, THAP7, CDC27, TRNAU1AP, UBE2E1, CAPRIN1, ETFBKMT, MRPL22, RPL27A, GAMT, ING4, RNF186, FBXO11, NDUFA12, PNPLA7, RPEL1, PPP2R5B, HAGH, MRPS18B, BAZ2A, NME7, DCAF11, MRPL39, DCAF4, AGO1, NSMCE3, EIF4ENIF1, DDB1, MARS2, PIGT, KLHL40, RP2, SUPT3H, ELP6, CCNG1, TPST2, CEP85, HBS1L, CPO, PFDN2, AGO4, C2CD3, FAM111B, NDUFA11, DDX25, TMX3, MTG1, PSMC1, SPOPL, PPWD1, NMNAT2, EIF5A2, JADE2, KSR2, CSNK2B, RBM3, MCTS1, PAK6, ABI1, PRMT7, DDA1, MAP3K8, UBE3A, ELOVL1, PSMD1, FXR1, SMURF2, LYPLA2, BAG1, PDCD6, PPM1H, TSSK1B, ATG16L2, CIAO2B, MOV10, KHSRP, DAP3, TRIM8, SMYD1, DNAJB1, DACT1, BIRC7, TRAP1, BOK, ITPKB, CAV3, MBOAT4, FERMT2, PTPRT, FGF16, PDE8B, ECM1, CAPN11, GUCY2C, FN1, MTMR4, PLA2G1B, EPHA1, AZIN2, TRAF3, TLR3, SERPINE1, PIP5KL1, EPHA5, SSH1, CARD16, SPRY2, BGN, NDST2, IGF1R, WFIKKN1, NOSIP, FCN3, SPINK5, PGAP1, DPEP3, ITGA1, LYG1, ABAT, CEACAM1, LOXL2, NUS1, ZDHHC8, IFRD2, KCNAB2, FOXL2, LDHC, FMO2, THOP1, UROD, FBXO33, TRAF2, MOS, FMO3, HDC, ADRA2A, DTX2, PRSS53, CDKN2C, UGT1A1, FKBP1B, PRRC1, SULT1B1, DUOXA2, TIMP2, CASP8, MBOAT1, CXXC1, MT-ATP8, PAM, OAT, ITPA, PHEX, DNPH1, PRMT1, NT5C1A, ARR3, STT3A, SCYL3, DDO, PHB2, SERPINB6, PM20D1, PRPS1, PRMT5, RNF31, HDAC1, PARP9, RPIA, TOB1, ETF1, ITM2B, RABGGTB, GPX1, HSP90AA1, GCH1, AMBP, NAGS, CTF1, PADI4, GOT1, HPRT1, MYRF, ENDOU, ADH5, DHX9, AOC2, CCND2, CAB39L, DOLK, YBX2, TAF7, RAD50, MRNIP, WARS2, ALG11, ZDHHC4, PPP4R2, TEX2, OTUB1, CLN3, CCNE2, TSSK4, PTPN20, UBE2D4, RPL6, RPS6KC1, ENTPD5, TRIM14, WWP1, GLYATL1, YWHAE, ALG3, KLHL7, DMAP1, B3GALT1, TP53INP2, MAP3K6, STK31, MAPK15, CHCHD4, YWHAG, NEK7, NMNAT3, GOLGA2, ADAMTS12, TMTC1, UBQLN1, PRKCH, ASB10, AHSG, LMTK2, LANCL1, GRB7, SLC2A9, IRAK4, FLOT1, COL3A1, SLCO1B1, IRAK1, ENPP1, LRIG2, PRKCA, PRICKLE1, SLC8A2, TMTC2, DDAH1, LIF, HCK, UBE2U, PNMT, DPYS, UTY, PLA2G4C, PCYT1B, MRPS30, SULT1C2, LAP3, SERPINB4, PANO1, EPGN, OXSM, PRNP, OGG1, EIF3K, HEG1, DPP7, SUMF1, NDST1, PFAS, TAB2, FGF4, CAAP1, ALDH5A1, HDAC6, TSPAN33, PIAS1, PUM1, METTL18, ABTB1, NAT10, NUDT17, FASTKD3, PCYT1A, B4GALT7, FBXL3, MRPS12, HHATL, PEF1, CSNK2A3, DTX4, PPP2R2A, USP25, PELI1, CARNS1, PRMT8, PTPN9, CPNE1, NEURL1B, PLD1, SBK2, PRKCQ, SPRY1, FGF19, SLC6A17, A4GNT, EMP2, ERBB4, ERLEC1, CERS5, SLC19A1, MT-TL1, SLC22A13, PRKAR2B, C4BPB, CEBPA, AGA, FES, ST6GALNAC3, IDH1, CPN1, CDK2, TALDO1, SAMD8, ASPG, G6PD, RPS16, ACR, HMCES, PIGA, ACY1 |
| developmental process | TIMM8A, FBXO22, SGCD, PSMA8, TEX15, NPHP3, SPEF2, WDR37, KATNB1, EXO1, ATF1, DNAJC30, PCDHGC4, CHEK1, GRSF1, CATSPER2, FANCD2, SRGAP2B, TUBB2B, YY1AP1, FBXO38, RNF151, WDR47, BBS4, CRYBB1, WDR36, ASXL2, SDHAF2, PPP2R5D, AGO2, NUDT21, ATXN3, NPAS4, NKD1, IGSF10, SPATA6, SRP54, PLET1, MYADM, SOX3, SVBP, CCR7, S100A1, ASPN, CTSZ, ORMDL3, OR51E2, CYSLTR1, TNFSF9, IL8RB, DLG3, CSF1R, COL4A2, NLGN3, CPNE6, EFEMP2, MFAP5, TMEM215, PKDCC, MYMK, MARCKSL1, SEMA4D, RAB27A, AP2B1, NXN, SERF1A, S100A9, COL13A1, PDPN, WASHC5, CLC, MYO9A, POU3F4, VASN, SLC25A27, SOCS3, LGI2, ZPBP2, MYH9, GJB3, SEMA6B, SMPD4, PAK3, LCP1, CNR1, PREX2, KANK2, ZBTB24, COL1A2, MITF, AHR, BHLHA9, GRK5, PLCD3, GREB1L, EVC, IFNA13, TSHB, NPY, IREB2, KRT16, HEMGN, POLE, SELENOF, HELLS, NR1D2, AIDA, HTRA2, NOCT, PRTN3, SECISBP2, TRIO, MT-CO1, GLO1, WARS, NES, TGM2, BRD1, GADD45B, CASZ1, UPRT, PIN1, AZI2, APC, KRT83, RHBDD3, SP5, LATS1, GLUL, METTL21C, CDC42SE1, MLLT3, INTS1, DZIP1, TUBB1, E2F8, CAPRIN1, GAMT, MARF1, ATP6V1G2, CREB3L2, CYB5R4, PPP2R5B, ACTR8, TASOR, AGO1, EIF4ENIF1, UBTFL1, IQCF1, DDB1, PIGT, KLHL40, CAPZB, ST7, CCNG1, AGO4, C2CD3, DDX25, KIF27, MIEF2, NMNAT2, EIF5A2, JADE2, CSNK2B, PAK6, ABI1, PRMT7, IRX1, UBE3A, ELOVL1, FXR1, MCF2, SMURF2, LYPLA2, PDCD6, PPP1R13L, TSSK1B, ZIC2, MOSPD3, MOV10, TRIM8, SMYD1, SETSIP, DNAJB1, DACT1, NDRG1, BIRC7, BOK, RBM46, HOXA4, LHX1, ITPKB, SDK1, DAAM1, CAV3, FERMT2, AKAP6, GREM2, SBF2, PCP4, FGF16, CXCL9, UNC5B, RAP1GAP, ECM1, ACKR3, ANGPTL3, FN1, SEC24D, MACO1, CPLX2, APLN, ASPM, EPHA1, RSPO3, AZIN2, FSCN2, DNM2, MMRN2, CRMP1, TLR3, FLRT3, SLC9A6, EXOC4, SERPINE1, TRPV2, LUZP1, EPHA5, RTN4R, SSH1, AP1G1, ATP2B2, SPRY2, PLPPR4, GPR171, EFNB2, BGN, SLC9A2, THBS3, WNT16, RIC8A, IGF1R, WFIKKN1, HOOK1, PCDH1, SLC26A2, SPINK5, PGAP1, ITGA1, JAM3, KIF5A, RAB33B, ABAT, CEACAM1, LOXL2, MYO9B, NUS1, CRTAC1, FOXL2, PCDHA8, CD79B, TPT1, UCP1, CSRNP1, UGT1A9, TNNT2, CDKN2C, GABRB3, UGT1A1, CHRND, YJEFN3, GDNF, FKBP1B, C14ORF93, TTC12, SULT1B1, CASP8, MBOAT1, KRT33B, GFI1B, SPRR2G, HSD17B2, XKR4, CFAP97, PHEX, DNPH1, PRMT1, KRT77, SCYL3, OPRM1, PHB2, CTNNA2, PRPS1, PRMT5, HDAC1, ZNF148, LMOD1, BSX, TOB1, ITM2B, GPX1, HSP90AA1, SPIC, KRT81, IRF1, CTF1, ZNF628, BIK, PADI4, MREG, LGALS1, HPRT1, MYRF, STMN1, GSTA2, KRT34, DHX9, ARID4A, SCO2, DDX3Y, RANBP3L, CDX2, DLEC1, YBX2, ABT1, WARS2, STRC, TMEM65, ZMYND15, TSSK4, MEIOC, MAL2, FOXB2, ARID4B, TFAP2E, WWP1, NCOA1, YWHAE, TP53INP2, YWHAG, ADAMTS12, FGFRL1, PRKCH, AHSG, GFRAL, CXCL14, CDH26, KCNQ4, KIDINS220, FLOT1, NEFH, CHODL, ATP8B1, COL3A1, GFRA3, LGI4, EFNA2, FGF11, CD47, SEMA3A, ENPP1, LRIG2, PRKCA, CSN3, ITGA11, SHC3, PRICKLE1, GPM6A, RBP4, DDAH1, ETS1, LIF, HCK, TBX6, ZNF536, PCYT1B, PYY, EMILIN1, NR0B1, POU1F1, ESRRB, SCT, ELL, EPGN, MAFF, HEG1, KRT23, NDST1, PFAS, PBX3, TAB2, KLF3, FGF4, OAS2, NRL, ATOH1, KRT5, ALDH5A1, HDAC6, PIAS1, ANKRD33, PUM1, SMARCD2, CEP152, HPS4, ANKRD17, SNW1, PEF1, TFEB, PTPN9, GPR15, CPNE1, TGFB1I1, AMIGO2, LRATD1, ARFRP1, DTNB, RFNG, KAZN, HOXB8, PRKCQ, PALLD, SPRY1, PCDH12, FGF19, SLC6A17, EMP2, ADAMTSL1, ANKS4B, ERBB4, POPDC2, CERS5, METRNL, EDA2R, TMEM204, FAM120B, CEBPA, FES, TBX15, CD34, FKBPL, IDH1, MIAT, NR2F6, G6PD, ZEB2, NANOG, PALM, HMCES, OLIG2, TNNC1 |
| anatomical structure development | TIMM8A, FBXO22, SGCD, NPHP3, SPEF2, WDR37, KATNB1, EXO1, ATF1, DNAJC30, PCDHGC4, CHEK1, GRSF1, CATSPER2, FANCD2, SRGAP2B, TUBB2B, FBXO38, WDR47, BBS4, CRYBB1, WDR36, ASXL2, SDHAF2, PPP2R5D, AGO2, NUDT21, ATXN3, NPAS4, NKD1, IGSF10, SRP54, PLET1, MYADM, SOX3, SVBP, CCR7, S100A1, ASPN, CTSZ, ORMDL3, CYSLTR1, TNFSF9, IL8RB, DLG3, CSF1R, COL4A2, NLGN3, CPNE6, EFEMP2, MFAP5, TMEM215, PKDCC, MYMK, MARCKSL1, SEMA4D, AP2B1, NXN, SERF1A, S100A9, COL13A1, PDPN, WASHC5, CLC, MYO9A, POU3F4, VASN, SLC25A27, SOCS3, LGI2, ZPBP2, MYH9, GJB3, SEMA6B, SMPD4, PAK3, LCP1, CNR1, PREX2, KANK2, ZBTB24, COL1A2, MITF, AHR, PLCD3, GREB1L, EVC, IFNA13, TSHB, NPY, IREB2, KRT16, POLE, SELENOF, HELLS, NR1D2, AIDA, HTRA2, NOCT, PRTN3, SECISBP2, TRIO, MT-CO1, GLO1, WARS, NES, TGM2, BRD1, CASZ1, UPRT, PIN1, AZI2, APC, KRT83, RHBDD3, SP5, LATS1, GLUL, METTL21C, CDC42SE1, MLLT3, INTS1, DZIP1, TUBB1, E2F8, CAPRIN1, GAMT, MARF1, CREB3L2, CYB5R4, PPP2R5B, ACTR8, TASOR, AGO1, EIF4ENIF1, UBTFL1, IQCF1, DDB1, PIGT, KLHL40, CAPZB, CCNG1, AGO4, C2CD3, DDX25, KIF27, MIEF2, NMNAT2, JADE2, CSNK2B, PAK6, ABI1, IRX1, UBE3A, ELOVL1, FXR1, MCF2, SMURF2, LYPLA2, PDCD6, PPP1R13L, TSSK1B, ZIC2, MOSPD3, MOV10, SMYD1, SETSIP, DNAJB1, DACT1, NDRG1, BIRC7, BOK, RBM46, HOXA4, LHX1, ITPKB, SDK1, DAAM1, CAV3, FERMT2, AKAP6, GREM2, SBF2, PCP4, FGF16, CXCL9, UNC5B, RAP1GAP, ECM1, ACKR3, ANGPTL3, FN1, SEC24D, MACO1, CPLX2, APLN, ASPM, EPHA1, RSPO3, FSCN2, DNM2, MMRN2, CRMP1, TLR3, FLRT3, SLC9A6, EXOC4, SERPINE1, TRPV2, LUZP1, EPHA5, RTN4R, SSH1, ATP2B2, SPRY2, PLPPR4, GPR171, EFNB2, BGN, SLC9A2, THBS3, WNT16, RIC8A, IGF1R, WFIKKN1, HOOK1, PCDH1, SLC26A2, SPINK5, PGAP1, ITGA1, JAM3, KIF5A, RAB33B, ABAT, CEACAM1, LOXL2, MYO9B, NUS1, CRTAC1, FOXL2, PCDHA8, CD79B, TPT1, CSRNP1, UGT1A9, TNNT2, CDKN2C, GABRB3, UGT1A1, CHRND, YJEFN3, GDNF, FKBP1B, TTC12, SULT1B1, CASP8, MBOAT1, KRT33B, GFI1B, SPRR2G, HSD17B2, XKR4, PHEX, DNPH1, PRMT1, KRT77, SCYL3, OPRM1, PHB2, CTNNA2, PRPS1, PRMT5, HDAC1, ZNF148, LMOD1, BSX, ITM2B, GPX1, HSP90AA1, SPIC, KRT81, IRF1, CTF1, BIK, LGALS1, HPRT1, MYRF, STMN1, GSTA2, KRT34, ARID4A, SCO2, RANBP3L, CDX2, YBX2, ABT1, WARS2, STRC, TMEM65, ZMYND15, TSSK4, MEIOC, MAL2, FOXB2, ARID4B, TFAP2E, WWP1, NCOA1, YWHAE, YWHAG, ADAMTS12, FGFRL1, PRKCH, AHSG, GFRAL, CXCL14, CDH26, KCNQ4, KIDINS220, FLOT1, NEFH, CHODL, ATP8B1, COL3A1, GFRA3, LGI4, EFNA2, FGF11, CD47, SEMA3A, ENPP1, LRIG2, PRKCA, CSN3, ITGA11, SHC3, PRICKLE1, GPM6A, RBP4, DDAH1, ETS1, LIF, HCK, TBX6, ZNF536, PCYT1B, PYY, EMILIN1, NR0B1, POU1F1, ESRRB, SCT, ELL, EPGN, MAFF, HEG1, KRT23, NDST1, PFAS, PBX3, TAB2, KLF3, FGF4, OAS2, NRL, ATOH1, KRT5, ALDH5A1, HDAC6, PIAS1, ANKRD33, SMARCD2, CEP152, ANKRD17, SNW1, PEF1, TFEB, PTPN9, GPR15, CPNE1, TGFB1I1, AMIGO2, LRATD1, ARFRP1, DTNB, RFNG, KAZN, HOXB8, PRKCQ, PALLD, SPRY1, PCDH12, FGF19, SLC6A17, EMP2, ADAMTSL1, ERBB4, POPDC2, CERS5, EDA2R, TMEM204, CEBPA, FES, TBX15, CD34, FKBPL, IDH1, NR2F6, G6PD, ZEB2, NANOG, PALM, HMCES, OLIG2, TNNC1 |
| cell communication | FBXO22, SGCD, NPHP3, RGL4, ATF1, CHEK1, ASB7, FANCD2, TUBB2B, PRPF19, USP47, BBS4, ASXL2, SDHAF2, PPP2R5D, GPR148, SERINC3, RASSF8, PDZD11, TMIGD3, AACS, ARL2BP, ATXN3, NPAS4, S100A11, NKD1, MYADM, FER1L5, SNX4, CCR7, S100A1, RANGRF, NPBWR1, CYSLTR2, ASPN, KCNMB1, OR51E2, TRPM5, GIPC1, CYSLTR1, KCNMB3, TNFSF9, SH3GLB1, IL8RB, CD59, XCL2, PLA2G4A, OR2W3, HCRTR1, DLG3, CSF1R, PITPNM2, COL4A2, NLGN3, TSLP, ARHGDIA, CCL17, SEMA4D, CNR2, NXN, HRH4, S100A9, PDPN, RGS13, CCL20, DRG2, MYO9A, VASN, SOCS3, OXER1, MYH9, GJB3, SEMA6B, SNAP23, BST1, PAK3, ARHGEF5, LCP1, CXCL16, CNR1, DOCK6, PREX2, KANK2, LMBRD2, COL1A2, RAB40A, ESRRA, OR52H1, HTR3A, FCGR2A, GNAI2, MITF, AHR, GRK5, OR2L13, FCER2, IL18RAP, PLCD3, ARHGAP23, NFKBIB, TAS2R19, ST18, EVC, IFNA13, TSHB, NPY, TRBV12-5, IL16, DYTN, HELLS, NR1D2, AIDA, HTRA2, TRIO, NPTX2, EFS, TGM2, GADD45B, FBXL20, SLC6A1, PIN1, FAM89B, JRK, AZI2, APC, RHBDD3, CD177, PFKM, TMBIM6, LATS1, GLUL, RASSF4, MIR320E, SETD9, METTL21C, CDC42SE1, MLLT3, XRCC3, DZIP1, ING4, RASSF7, RNF186, DYSF, CYB5R4, PPP2R5B, DDB1, BEX5, RANBP1, CCNG1, HBS1L, C2CD3, ARFGEF2, JADE2, KSR2, CSNK2B, CCDC68, PAK6, ABI1, MAP3K8, UBE3A, FXR1, MCF2, SMURF2, BAG1, PDCD6, GOLT1B, TSSK1B, DAP3, TRIM8, SESN3, DACT1, NDRG1, BIRC7, TRAP1, BOK, ARHGEF10L, LHX1, ITPKB, HPGDS, DAAM1, PEX5L, JPH4, CAV3, LTBP2, ARAP2, FERMT2, AKAP6, PTPRT, ASIC3, GREM2, PCP4, FGF16, PDE8B, CXCL9, GPR6, UNC5B, RAP1GAP, ECM1, INSYN1, ACKR3, TRIL, GUCY2C, ANGPTL3, FN1, PAG1, MTMR4, MACO1, OSMR, PLA2G1B, CPLX2, OLFML2A, APLN, RAB8B, ASPM, EPHA1, RSPO3, SLC15A3, TRAF3, DNM2, MMRN2, PDGFRL, TLR3, FLRT3, SLC9A6, EXOC4, SERPINE1, EPHA5, RTN4R, SSH1, CARD16, INSL5, SPRY2, GPR34, CHRNB4, PLPPR4, GPR171, EFNB2, WNT16, IL17RE, RIC8A, IGF1R, WFIKKN1, RAPGEF4, CEACAM7, PCDH1, OLFML3, FCN3, PATJ, ITGA1, JAM3, KIF5A, RAB33B, ABAT, CACNA1I, CEACAM1, MYO9B, NUS1, HCN2, FOXL2, CGB5, CD79B, OR56A5, CGB8, TPT1, HTR7, OR6B3, TRAF2, MOS, CSRNP1, OR2L8, OR11L1, ADRA2A, DTX2, GABRB3, CHRND, YJEFN3, GDNF, FKBP1B, TAS2R31, DGKD, CASP8, PLCD1, PHEX, PRMT1, ARR3, OPRM1, PHB2, IKBKG, PRMT5, RNF31, HDAC1, PARP9, MAGEA1, TOB1, GPX1, HSP90AA1, AMBP, IRF1, CTF1, BIK, LGALS1, GOT1, STMN1, ENDOU, ARID4A, CAB39L, TAF7, RAD50, MRNIP, SFPQ, SRPX, TEX2, CLN3, TSSK4, RPS6KC1, RASSF6, PSD4, HDGF, ARID4B, WWP1, NCOA1, YWHAE, MAP3K6, NCOA5, MAPK15, YWHAG, NEK7, GNG2, ADAMTS12, UBQLN1, FGFRL1, PRKCH, GPR142, ASB10, GPR87, BTN3A3, AHSG, GFRAL, LANCL1, CXCL14, GRB7, SIGLEC8, F2RL3, IRAK4, KIDINS220, FLOT1, NEFH, COL3A1, CLEC16A, VWA2, GFRA3, IRAK1, EFNA2, FGF11, CD47, CAPS, SEMA3A, ENPP1, LRIG2, SOST, PRKCA, ITGA11, SHC3, PRICKLE1, SLC8A2, RBP4, DDAH1, FFAR1, GNG10, LIF, OR2AP1, MAFA, HCK, RRP22, CORT, PLEKHG7, TBX6, ZNF536, PLA2G4C, CYP7A1, PYY, EMILIN1, TAS2R60, NR0B1, SLC6A2, ESRRB, SCT, MCHR1, EPGN, PRNP, TFG, HEG1, NDST1, TAB2, FGF4, OAS2, SHC2, ATOH1, CAAP1, HDAC6, PPP4C, PIAS1, PUM1, TOM1, ANKRD17, SNW1, DTX4, PELI1, TFEB, GPR15, CPNE1, NEURL1B, TGFB1I1, USO1, XPR1, ARFRP1, DTNB, RFNG, SYT5, RGS11, PLPPR3, PLD1, SBK2, NCR3, CXCL6, PRKCQ, SV2A, PILRA, LMCD1, PLPPR2, SPRY1, FGF19, DLGAP1, EMP2, ERBB4, ERLEC1, METRNL, EDA2R, RASGEF1B, TMEM204, FAM120B, PRKAR2B, ARHGAP27, CEBPA, STON1, FES, FOLR2, CD34, CDK2, PTGFR, NR2F6, ZEB2, SST, PALM |
| cellular response to stimulus | FBXO22, SGCD, SSRP1, TEX15, NPHP3, RGL4, EXO1, ATF1, MCM7, CHEK1, ASB7, FANCD2, PRPF19, SLC25A14, UVSSA, USP47, BBS4, ASXL2, SDHAF2, PPP2R5D, GPR148, SERINC3, RASSF8, TMIGD3, ARL2BP, ATXN3, NPAS4, S100A11, NKD1, SRP54, MYADM, CCR7, S100A1, RANGRF, NPBWR1, CYSLTR2, ASPN, KCNMB1, OR51E2, TRPM5, GIPC1, CYSLTR1, TNFSF9, SH3GLB1, IL8RB, CD59, XCL2, PLA2G4A, OR2W3, HCRTR1, CSF1R, PITPNM2, COL4A2, NLGN3, CPNE6, TSLP, PKLR, ARHGDIA, CCL17, SEMA4D, CNR2, NXN, HRH4, S100A9, PDPN, RGS13, CCL20, DRG2, MYO9A, VASN, SOCS3, OXER1, MYH9, GJB3, SEMA6B, NPC1L1, SMPD4, BST1, PAK3, ARHGEF5, LCP1, CXCL16, CNR1, DOCK6, PREX2, KANK2, LMBRD2, COL1A2, RAB40A, SERPINB9, ESRRA, OR52H1, HTR3A, FCGR2A, GNAI2, MITF, AHR, GRK5, OR2L13, FCER2, IL18RAP, PLCD3, ARHGAP23, NFKBIB, TAS2R19, ST18, EVC, ECHDC3, IFNA13, TSHB, NPY, TRBV12-5, IL16, POLE, SELENOF, HELLS, NR1D2, AIDA, HTRA2, RFC2, TRIO, NPTX2, EFS, TGM2, GADD45B, HP, PIN1, ZNF580, ALDH3A1, FAM89B, JRK, AZI2, APC, RHBDD3, CD177, SP5, TMBIM6, LATS1, GLUL, NEIL2, RASSF4, MIR320E, SETD9, METTL21C, CDC42SE1, MLLT3, XRCC3, BAZ1B, DZIP1, MARF1, ING4, RASSF7, RNF186, CREB3L2, PPP2R5B, ACTR8, NSMCE3, DDB1, BEX5, SUPT3H, RANBP1, CCNG1, HBS1L, C2CD3, ARFGEF2, ERCC4, POLD2, SWI5, RECQL, JADE2, KSR2, CSNK2B, CCDC68, MCTS1, PAK6, ABI1, MAP3K8, UBE3A, MCF2, SMURF2, BAG1, PDCD6, GOLT1B, TSSK1B, KHSRP, DAP3, TRIM8, SESN3, DNAJB1, DACT1, NDRG1, BIRC7, TRAP1, BOK, ARHGEF10L, ITPKB, HPGDS, DAAM1, PEX5L, CAV3, LTBP2, ARAP2, FERMT2, AKAP6, PTPRT, ASIC3, GREM2, PCP4, FGF16, PDE8B, CXCL9, GPR6, UNC5B, RAP1GAP, ECM1, INSYN1, ACKR3, TRIL, GUCY2C, ANGPTL3, FN1, PAG1, MTMR4, MACO1, OSMR, PLA2G1B, OLFML2A, APLN, ASPM, EPHA1, RSPO3, SLC15A3, TRAF3, DNM2, MMRN2, PDGFRL, TLR3, FLRT3, SLC9A6, SERPINE1, EPHA5, RTN4R, SSH1, AP1G1, MFAP4, RYR3, CARD16, INSL5, SPRY2, GPR34, CHRNB4, PLPPR4, GPR171, EFNB2, WNT16, IL17RE, RIC8A, IGF1R, WFIKKN1, RAPGEF4, CEACAM7, OLFML3, FCN3, PATJ, ITGA1, JAM3, RAB33B, ABAT, CACNA1I, CEACAM1, MYO9B, NUS1, DEFB104A, HCN2, CGB5, CD79B, GBP6, OR56A5, CGB8, TPT1, FMO2, HTR7, UCP1, OR6B3, TRAF2, MOS, CSRNP1, UGT1A9, OR2L8, OR11L1, ADRA2A, DTX2, MT1B, GABRB3, HBB, UGT1A1, CHRND, YJEFN3, GDNF, FKBP1B, TAS2R31, SULT1B1, DGKD, CASP8, PLCD1, CREB3L3, CES2, PHEX, PRMT1, ARR3, OPRM1, PHB2, IKBKG, SERPINB6, PRMT5, RNF31, HDAC1, PARP9, GPX5, MAGEA1, TOB1, GPX1, HSP90AA1, GCH1, AMBP, IRF1, CTF1, BIK, LGALS1, GOT1, STMN1, GSTA2, ENDOU, ADH5, DHX9, ARID4A, AOC2, CCND2, CAB39L, TAF7, ZNF277, RAD50, MRNIP, SFPQ, SRPX, PPP4R2, TEX2, OTUB1, CLN3, TSSK4, MEIOC, NEIL3, RPS6KC1, RASSF6, PSD4, SRSF3, HDGF, ARID4B, WWP1, NCOA1, YWHAE, DMAP1, MAP3K6, NCOA5, MAPK15, YWHAG, NEK7, GNG2, ADAMTS12, UBQLN1, FGFRL1, PRKCH, GPR142, ASB10, GPR87, BTN3A3, AHSG, GFRAL, LANCL1, CXCL14, GRB7, SIGLEC8, F2RL3, IRAK4, KIDINS220, FLOT1, NEFH, PXDNL, COL3A1, CLEC16A, VWA2, GFRA3, SLCO1B1, IRAK1, EFNA2, FGF11, CD47, CAPS, SEMA3A, ENPP1, LRIG2, SOST, PRKCA, ITGA11, SHC3, PRICKLE1, SLC8A2, DDAH1, FFAR1, GNG10, LIF, OR2AP1, MAFA, HCK, RRP22, CORT, PLEKHG7, TBX6, EPHX1, ZNF536, PLA2G4C, CYP7A1, PYY, EMILIN1, TAS2R60, NR0B1, ESRRB, AS3MT, SCT, MCHR1, EPGN, PRNP, OGG1, TFG, HEG1, NDST1, TAB2, KLF3, FGF4, OAS2, SHC2, ATOH1, CAAP1, HDAC6, PPP4C, PIAS1, ABTB2, PUM1, TOM1, SMARCD2, ANKRD17, SNW1, DTX4, USP25, PELI1, TFEB, GPR15, CPNE1, NEURL1B, TGFB1I1, USO1, XPR1, ARFRP1, RFNG, SYT5, PRDX6, RGS11, PLPPR3, PLD1, SBK2, NCR3, CXCL6, PRKCQ, PILRA, LMCD1, PLPPR2, SPRY1, FGF19, DLGAP1, EMP2, ANKS4B, ERBB4, ERLEC1, METRNL, EDA2R, RASGEF1B, TMEM204, FAM120B, PRKAR2B, ARHGAP27, CEBPA, STON1, FES, FOLR2, CD34, CYP2S1, NFE2L3, ALKBH3, CDK2, PTGFR, NR2F6, G6PD, RPS16, ZEB2, SST, PALM, HMCES, PIGA |
| signaling | SGCD, NPHP3, RGL4, ATF1, CHEK1, ASB7, FANCD2, TUBB2B, PRPF19, USP47, BBS4, ASXL2, SDHAF2, PPP2R5D, GPR148, SERINC3, RASSF8, PDZD11, TMIGD3, AACS, ARL2BP, ATXN3, NPAS4, S100A11, NKD1, MYADM, FER1L5, SNX4, CCR7, S100A1, RANGRF, NPBWR1, CYSLTR2, ASPN, KCNMB1, OR51E2, TRPM5, GIPC1, CYSLTR1, TNFSF9, IL8RB, CD59, XCL2, PLA2G4A, OR2W3, HCRTR1, DLG3, CSF1R, PITPNM2, COL4A2, NLGN3, TSLP, ARHGDIA, CCL17, SEMA4D, CNR2, NXN, HRH4, S100A9, PDPN, RGS13, CCL20, DRG2, MYO9A, VASN, SOCS3, OXER1, MYH9, GJB3, SEMA6B, SNAP23, BST1, PAK3, ARHGEF5, LCP1, CXCL16, CNR1, DOCK6, PREX2, KANK2, LMBRD2, COL1A2, RAB40A, ESRRA, OR52H1, HTR3A, FCGR2A, GNAI2, MITF, AHR, GRK5, OR2L13, FCER2, IL18RAP, PLCD3, ARHGAP23, NFKBIB, TAS2R19, ST18, EVC, IFNA13, TSHB, NPY, TRBV12-5, IL16, DYTN, HELLS, NR1D2, AIDA, HTRA2, TRIO, NPTX2, EFS, TGM2, GADD45B, FBXL20, SLC6A1, PIN1, FAM89B, JRK, AZI2, APC, RHBDD3, CD177, PFKM, TMBIM6, LATS1, RASSF4, MIR320E, SETD9, METTL21C, CDC42SE1, MLLT3, XRCC3, DZIP1, ING4, RASSF7, RNF186, DYSF, CYB5R4, PPP2R5B, DDB1, BEX5, RANBP1, CCNG1, HBS1L, C2CD3, ARFGEF2, JADE2, KSR2, CSNK2B, CCDC68, PAK6, ABI1, MAP3K8, UBE3A, FXR1, MCF2, SMURF2, BAG1, PDCD6, GOLT1B, TSSK1B, DAP3, TRIM8, SESN3, DACT1, NDRG1, BIRC7, TRAP1, BOK, ARHGEF10L, LHX1, ITPKB, HPGDS, DAAM1, PEX5L, JPH4, CAV3, LTBP2, ARAP2, FERMT2, AKAP6, PTPRT, ASIC3, GREM2, PCP4, FGF16, PDE8B, CXCL9, GPR6, UNC5B, RAP1GAP, ECM1, INSYN1, ACKR3, TRIL, GUCY2C, ANGPTL3, FN1, PAG1, MTMR4, MACO1, OSMR, PLA2G1B, CPLX2, OLFML2A, APLN, RAB8B, ASPM, EPHA1, RSPO3, SLC15A3, TRAF3, DNM2, MMRN2, PDGFRL, TLR3, FLRT3, SLC9A6, EXOC4, SERPINE1, EPHA5, RTN4R, SSH1, CARD16, INSL5, SPRY2, GPR34, CHRNB4, PLPPR4, GPR171, EFNB2, WNT16, IL17RE, RIC8A, IGF1R, WFIKKN1, RAPGEF4, CEACAM7, PCDH1, OLFML3, FCN3, PATJ, ITGA1, KIF5A, RAB33B, ABAT, CACNA1I, CEACAM1, MYO9B, NUS1, HCN2, FOXL2, CGB5, CD79B, OR56A5, CGB8, TPT1, HTR7, OR6B3, TRAF2, MOS, CSRNP1, OR2L8, OR11L1, ADRA2A, DTX2, GABRB3, CHRND, YJEFN3, GDNF, FKBP1B, TAS2R31, DGKD, CASP8, PLCD1, PHEX, PRMT1, ARR3, OPRM1, PHB2, IKBKG, PRMT5, RNF31, HDAC1, PARP9, MAGEA1, TOB1, GPX1, HSP90AA1, AMBP, IRF1, CTF1, BIK, LGALS1, GOT1, STMN1, ENDOU, ARID4A, CAB39L, TAF7, RAD50, MRNIP, SFPQ, SRPX, TEX2, CLN3, TSSK4, RPS6KC1, RASSF6, PSD4, TBC1D16, HDGF, ARID4B, WWP1, NCOA1, YWHAE, MAP3K6, NCOA5, MAPK15, YWHAG, NEK7, GNG2, ADAMTS12, UBQLN1, FGFRL1, PRKCH, GPR142, ASB10, GPR87, BTN3A3, AHSG, GFRAL, LANCL1, CXCL14, GRB7, SIGLEC8, F2RL3, IRAK4, KIDINS220, FLOT1, NEFH, COL3A1, CLEC16A, VWA2, GFRA3, IRAK1, EFNA2, FGF11, CD47, CAPS, SEMA3A, ENPP1, LRIG2, SOST, PRKCA, ITGA11, SHC3, PRICKLE1, SLC8A2, RBP4, DDAH1, FFAR1, GNG10, LIF, OR2AP1, MAFA, HCK, RRP22, CORT, PLEKHG7, TBX6, ZNF536, PLA2G4C, CYP7A1, PYY, EMILIN1, TAS2R60, NR0B1, SLC6A2, ESRRB, SCT, MCHR1, EPGN, PRNP, TFG, HEG1, NDST1, TAB2, FGF4, OAS2, SHC2, ATOH1, CAAP1, HDAC6, PPP4C, PIAS1, PUM1, TOM1, ANKRD17, SNW1, DTX4, PELI1, GPR15, CPNE1, NEURL1B, TGFB1I1, USO1, ARFRP1, DTNB, RFNG, SYT5, RGS11, PLPPR3, PLD1, SBK2, NCR3, CXCL6, PRKCQ, SV2A, PILRA, LMCD1, PLPPR2, SPRY1, FGF19, DLGAP1, EMP2, ERBB4, ERLEC1, METRNL, EDA2R, RASGEF1B, TMEM204, FAM120B, PRKAR2B, ARHGAP27, CEBPA, STON1, CD34, CDK2, PTGFR, NR2F6, ZEB2, SST, PALM |
| multicellular organism development | TIMM8A, SGCD, NPHP3, SPEF2, WDR37, KATNB1, EXO1, ATF1, DNAJC30, PCDHGC4, CHEK1, GRSF1, FANCD2, SRGAP2B, TUBB2B, FBXO38, WDR47, BBS4, CRYBB1, WDR36, PPP2R5D, AGO2, ATXN3, NPAS4, NKD1, IGSF10, SRP54, MYADM, SOX3, SVBP, S100A1, ASPN, CTSZ, ORMDL3, CYSLTR1, TNFSF9, IL8RB, CSF1R, COL4A2, NLGN3, CPNE6, EFEMP2, MFAP5, TMEM215, PKDCC, MARCKSL1, SEMA4D, AP2B1, NXN, SERF1A, S100A9, COL13A1, PDPN, WASHC5, CLC, MYO9A, POU3F4, SOCS3, LGI2, MYH9, GJB3, SEMA6B, SMPD4, PAK3, CNR1, PREX2, KANK2, COL1A2, MITF, AHR, PLCD3, GREB1L, EVC, NPY, IREB2, POLE, HELLS, AIDA, HTRA2, NOCT, SECISBP2, TRIO, MT-CO1, WARS, NES, TGM2, BRD1, CASZ1, PIN1, APC, RHBDD3, SP5, LATS1, GLUL, MLLT3, INTS1, DZIP1, TUBB1, E2F8, CAPRIN1, CREB3L2, PPP2R5B, ACTR8, TASOR, AGO1, EIF4ENIF1, UBTFL1, PIGT, AGO4, C2CD3, KIF27, NMNAT2, JADE2, CSNK2B, PAK6, ABI1, IRX1, UBE3A, FXR1, MCF2, SMURF2, LYPLA2, PDCD6, PPP1R13L, ZIC2, MOSPD3, MOV10, SMYD1, DNAJB1, DACT1, NDRG1, BIRC7, BOK, RBM46, HOXA4, LHX1, ITPKB, SDK1, CAV3, FERMT2, AKAP6, GREM2, SBF2, PCP4, UNC5B, RAP1GAP, ECM1, ACKR3, ANGPTL3, FN1, SEC24D, MACO1, CPLX2, APLN, ASPM, EPHA1, RSPO3, FSCN2, DNM2, MMRN2, CRMP1, TLR3, FLRT3, SLC9A6, EXOC4, SERPINE1, TRPV2, LUZP1, EPHA5, RTN4R, ATP2B2, SPRY2, PLPPR4, GPR171, EFNB2, BGN, THBS3, WNT16, RIC8A, IGF1R, WFIKKN1, PCDH1, SLC26A2, SPINK5, PGAP1, ITGA1, JAM3, KIF5A, RAB33B, ABAT, CEACAM1, LOXL2, NUS1, CRTAC1, FOXL2, PCDHA8, CSRNP1, UGT1A9, TNNT2, CDKN2C, GABRB3, UGT1A1, YJEFN3, GDNF, FKBP1B, CASP8, MBOAT1, GFI1B, HSD17B2, PHEX, PRMT1, SCYL3, OPRM1, PHB2, CTNNA2, PRPS1, PRMT5, HDAC1, ZNF148, ITM2B, GPX1, HSP90AA1, SPIC, IRF1, CTF1, BIK, HPRT1, MYRF, STMN1, ARID4A, SCO2, RANBP3L, CDX2, ABT1, WARS2, STRC, TMEM65, MAL2, ARID4B, WWP1, NCOA1, YWHAE, YWHAG, ADAMTS12, FGFRL1, PRKCH, AHSG, GFRAL, KCNQ4, KIDINS220, FLOT1, NEFH, CHODL, ATP8B1, COL3A1, GFRA3, LGI4, EFNA2, FGF11, CD47, SEMA3A, ENPP1, LRIG2, PRKCA, SHC3, PRICKLE1, GPM6A, RBP4, DDAH1, ETS1, LIF, TBX6, ZNF536, PCYT1B, PYY, EMILIN1, NR0B1, POU1F1, SCT, ELL, EPGN, MAFF, HEG1, NDST1, PBX3, TAB2, FGF4, OAS2, NRL, ATOH1, ALDH5A1, HDAC6, SMARCD2, ANKRD17, SNW1, PEF1, TFEB, PTPN9, GPR15, CPNE1, TGFB1I1, AMIGO2, ARFRP1, DTNB, RFNG, HOXB8, PRKCQ, PALLD, SPRY1, PCDH12, FGF19, SLC6A17, EMP2, ADAMTSL1, ERBB4, POPDC2, CERS5, TMEM204, CEBPA, FES, TBX15, CD34, FKBPL, IDH1, NR2F6, G6PD, ZEB2, NANOG, PALM, HMCES, OLIG2, TNNC1 |
| regulation of nitrogen compound metabolic process | FBXO22, CHD1, CIZ1, ALG10B, TEX15, EIF4E3, F8A1, LARP1B, ATF1, MCM7, DNAJC30, CHEK1, GRSF1, FANCD2, PRPF19, HOXC12, YY1AP1, USP47, SERTAD1, BEX4, OTUD6B, ASXL2, SP140L, HIC2, PPP2R5D, AGO2, SRSF9, TMIGD3, ARL2BP, ATXN3, NPAS4, S100A11, NKD1, YAF2, MYADM, SOX3, SVBP, CCR7, CTSZ, ORMDL3, GIPC1, ZNF311, SFSWAP, DLG3, CSF1R, TSLP, SEMA4D, NXN, S100A9, POU3F4, SOCS3, WFDC12, MYH9, BST1, ARHGEF5, KANK2, ZBTB24, ZNF343, SERPINB9, ESRRA, GNAI2, MITF, ZNF140, AHR, BHLHA9, ZNF75A, IL18RAP, NFKBIB, ST18, IFNA13, ZNF649, IREB2, INTS5, MXD3, NR1D2, AIDA, HTRA2, RFC2, NOCT, SECISBP2, GLO1, WARS, BRD1, GADD45B, CASZ1, FBXL20, SERPING1, PIN1, ZNF580, APC, RHBDD3, SP5, PFKM, TMBIM6, LATS1, PTMS, DCP1B, ZNF471, POLDIP3, DRAP1, MLLT3, BAZ1B, BCDIN3D, INTS1, THAP7, ZNF593, TRNAU1AP, UBE2E1, E2F8, CAPRIN1, ING4, CREB3L2, PPP2R5B, BAZ2A, ACTR8, TASOR, HEATR1, AGO1, NSMCE3, EIF4ENIF1, UBTFL1, DDB1, KLHL40, SUPT3H, ELP6, CCNG1, CEP85, C1D, HBS1L, PFDN2, AGO4, C2CD3, DDX25, MLLT6, HSF5, LCORL, ERCC4, ARMCX3, MTG1, PSMC1, SPOPL, NMNAT2, EIF5A2, JADE2, CSNK2B, RBM3, PAK6, ABI1, HMGB4, DDA1, IRX1, UBE3A, PSMD1, FXR1, SMURF2, ZNF91, PDCD6, PPP1R13L, ZIC2, MOV10, KHSRP, TRIM8, SMYD1, SETSIP, DNAJB1, DACT1, BIRC7, TRAP1, BOK, RBM46, HOXA4, LHX1, ITPKB, CAV3, FERMT2, PTPRT, FGF16, ECM1, FN1, PLA2G1B, APLN, EPHA1, AZIN2, TRAF3, DNM2, TLR3, SERPINE1, PIP5KL1, EPHA5, SSH1, CARD16, SPRY2, IGF1R, WFIKKN1, FCN3, SPINK5, ITGA1, ABAT, CEACAM1, LOXL2, IFRD2, FOXL2, ZNF19, UCP1, HOMEZ, FBXO33, TRAF2, CSRNP1, ADRA2A, IKZF4, CDKN2C, HBB, ZNF607, GDNF, FKBP1B, PRRC1, ZNF215, ZNF184, ZNF230, DUOXA2, TIMP2, CASP8, GFI1B, CCNQ, CXXC1, CRTC1, CREB3L3, ZNF799, MAGEB6, PRMT1, ARR3, OPRM1, PHB2, IKBKG, SERPINB6, PRMT5, RNF31, HDAC1, PARP9, ZNF148, BSX, MAGEA1, TOB1, ETF1, ITM2B, GPX1, HSP90AA1, SPIC, AMBP, IRF1, CTF1, ZNF628, TSTD1, HPRT1, MYRF, DHX9, ARID4A, ZNF707, CDX2, KLF12, CCND2, ZNF652, CAB39L, GLI4, ZNF732, GTF2A1, PHF21A, YBX2, ABT1, TAF7, RAD50, MRNIP, SFPQ, PPP4R2, OTUB1, CLN3, CCNE2, ZMYND15, TSSK4, MEIOC, RPL6, INTS11, SRSF3, TRIM14, FOXB2, HDGF, ARID4B, TFAP2E, WWP1, NCOA1, YWHAE, DMAP1, ZNF467, TP53INP2, NCOA5, MAPK15, SSX4, YWHAG, NEK7, GOLGA2, UBQLN1, PRKCH, AHSG, GRB7, FLOT1, ATP8B1, IRAK1, CD47, ENPP1, LRIG2, SOST, PRKCA, PRICKLE1, SLC8A2, DDAH1, ETS1, LIF, MAFA, HCK, TBX6, ZNF536, ZNF69, MAGEB4, NR0B1, POU1F1, ZNF30, ESRRB, SERPINB4, RBM5, ELL, PANO1, EPGN, PRNP, MAFF, OGG1, EIF3K, HEG1, PBX3, TAB2, KLF3, FGF4, OAS2, NRL, ATOH1, CAAP1, HDAC6, PPP4C, PIAS1, ZKSCAN1, POLR3F, PUM1, METTL18, L3MBTL2, NAT10, SMARCD2, FASTKD3, RBM15B, ANKRD17, SNW1, SERTAD2, HHATL, TCERG1, PEF1, CSNK2A3, USP25, ZC3H10, PELI1, TFEB, TGFB1I1, PRDX6, PLD1, HOXB8, PRKCQ, LMCD1, SPRY1, FGF19, EMP2, ERBB4, EDA2R, PRKAR2B, C4BPB, CEBPA, TBX15, NFE2L3, CDK2, SAMD8, NR2F6, G6PD, ZEB2, NANOG, HMCES, OLIG2, POU2F1 |
| cytoplasm | TIMM8A, FBXO22, SGCD, CHD1, PSMA8, CCDC8, ALG10B, TEX15, UTP20, RPGR, EIF4E3, NPHP3, EFCAB6, SPEF2, WDR37, MROH1, NSFL1C, UBE2G1, KATNB1, STARD9, F8A1, LARP1B, CDADC1, SLAIN1, RGL4, ACBD4, UCK1, CFAP36, MCM7, PIGH, NACAD, DNAJC30, CHEK1, GRSF1, RPS27A, ASB7, FANCD2, TOP1MT, SNRNP48, TUBB2B, KIFC2, PRPF19, YY1AP1, SLC25A14, GOT1L1, USP47, COPB1, FBXO38, RNF151, SERTAD1, SSR4, BEX4, OTUD6B, WDR47, BBS4, CHDH, SDHAF2, PYCR3, KPNA3, PPP2R5D, PSMB2, AGO2, ANXA10, SDR9C7, MTMR11, NUDT21, BCL2L15, FKBP9, RSPH14, SERINC3, PDZD11, PHLDA1, ENDOD1, AACS, ARL2BP, ATXN3, NPAS4, DPP9, ARMC8, S100A11, NKD1, MIA2, HSDL2, YAF2, SRP54, MTFP1, MYADM, FER1L5, SVBP, AK7, SNX4, CCR7, S100A1, RANGRF, NEU2, DIAPH3, CTSZ, ORMDL3, PPP1R3G, OR51E2, FCHO2, GIPC1, SH3GLB1, IL8RB, CD59, KCNAB3, PLA2G4A, SLK, RAB2A, DLG3, ST3GAL4, PITPNM2, COL4A2, TARM1, NLGN3, CPNE6, PKLR, ARHGDIA, PKDCC, MYMK, MARCKSL1, PCYOX1, TNFAIP8L2, LIMD2, KIF21A, RAB27A, AP2B1, SLC9A3, MUC16, CNR2, NXN, SERF1A, S100A9, COL13A1, PDPN, RGS13, WASHC5, CD93, CLC, CAPN8, DRG2, MYO9A, VASN, SEC16B, SLC25A27, SOCS3, CD1B, SLC4A1AP, CLIC6, ZPBP2, MYH9, MUC13, GJB3, CNST, NPC1L1, SNAP23, SMPD4, CAPZA1, BST1, PAK3, ARHGEF5, LCP1, CNR1, DOCK6, GIMAP5, PREX2, KANK2, PDHA2, ESPNL, COL1A2, FGGY, CYP3A43, RAB40A, RDH5, SERPINB9, ESRRA, CHST13, CYP27C1, CCDC92, FCGR2A, GNAI2, MITF, AHR, BHLHA9, GRK5, PLCD3, ARHGAP23, NFKBIB, DMAC2, EVC, ECHDC3, TSHB, A1BG, NPY, IREB2, GALNT15, INTS5, KRT16, IL16, SELENOF, NR1D2, AIDA, HTRA2, NOCT, PRTN3, SECISBP2, TRIO, MT-CO1, EFS, GLO1, WARS, NES, TGM2, MTHFS, CLPP, GADD45B, HEBP2, CASZ1, METTL7B, WDR83OS, FBXL20, HP, SERPING1, UPRT, ABCC10, PIN1, MOXD1, ALDH3A1, AKR1E2, CARS, FAM89B, JRK, APOBEC2, AZI2, APC, TMED5, KRT83, PGK2, CD177, NUSAP1, AFMID, PFKM, TMBIM6, LATS1, GLUL, NDUFB3, NEIL2, ZDHHC18, RGPD3, DCP1B, POLDIP3, METTL21C, CDC42SE1, MLLT3, XRCC3, SKIV2L, BCDIN3D, SRP68, DZIP1, TUBB1, ZNF593, CDC27, ANKRD16, TRNAU1AP, COX14, UBE2E1, E2F8, CAPRIN1, ETFBKMT, MRPL22, RPL27A, GAMT, TFB1M, MARF1, ING4, RASSF7, RNF186, ATP6V1G2, DYSF, FBXO11, NDUFA12, CREB3L2, PNPLA7, RPEL1, CYB5R4, PPP2R5B, HAGH, MRPS18B, BAZ2A, POC5, NME7, MRPL39, RANBP3, POLR2C, HEATR1, AGO1, NSMCE3, EIF4ENIF1, UBTFL1, TMEM87A, IQCF1, DDB1, BEX5, MARS2, POMP, VPS50, PIGT, KLHL40, HAUS7, EMC2, RP2, CKAP2L, METTL2A, ELP6, CAPZB, RANBP1, CCNG1, PIH1D2, TPST2, CEP85, POLR3K, C1D, HBS1L, PFDN2, CEP192, AGO4, C2CD3, NDUFA11, DDX25, KIF27, MIEF2, ARFGEF2, TMX3, ARMCX3, MTG1, RECQL, PSMC1, SPOPL, IPO13, EPB41L4A, NMNAT2, SRSF5, EIF5A2, KSR2, CSNK2B, CCDC68, RBM3, MCTS1, PAK6, ABI1, PRMT7, MAP3K8, DENND2A, UBE3A, ELOVL1, PSMD1, SH3BGRL2, FXR1, MCF2, SMURF2, LYPLA2, BAG1, PDCD6, GOLT1B, SEC61G, PPM1H, PPP1R13L, TSSK1B, ATG16L2, CIAO2B, ZIC2, MOSPD3, ZNF185, MOV10, KHSRP, DAP3, SUGT1, TRIM8, HELLPAR, SMYD1, SESN3, SETSIP, DNAJB1, DACT1, NDRG1, BIRC7, ESYT1, TRAP1, SSR2, BOK, SAMM50, SLC36A4, ARHGEF10L, BCS1L, ITPKB, CCDC170, HPGDS, DAAM1, PEX5L, EPN3, JPH4, CAV3, ARAP2, GCC2, MBOAT4, FERMT2, AKAP6, ASIC3, KCNK9, AAGAB, SBF2, PCP4, FGF16, PROCR, YKT6, PDE8B, OMD, RAP1GAP, ECM1, ACKR3, CAPN11, GUCY2C, MYOM3, OSBPL9, ANGPTL3, FN1, MTMR4, SEC24D, MACO1, CPLX2, RAB8B, ASPM, SLC15A3, AZIN2, FSCN2, TRAF3, MCEMP1, DNM2, CRMP1, TLR3, FLRT3, SLC9A6, EXOC4, SERPINE1, RAB3IP, TRPV2, LUZP1, PIP5KL1, EPHA5, RTN4R, SSH1, SMAGP, AP1G1, ATP2B2, RYR3, CARD16, SPRY2, CHRNB4, BGN, THBS3, WNT16, IL17RE, RIC8A, NDST2, RAPGEF4, NOSIP, HOOK1, CEACAM7, SPINK5, PATJ, PGAP1, DPEP3, TMEM230, ITGA1, JAM3, KIF5A, RAB33B, ABAT, CEACAM1, LOXL2, MYO9B, NUS1, ZDHHC8, KCNAB2, FADS2, CGB5, GBP6, CGB8, WSCD1, LDHC, TPT1, FMO2, HTR7, UCP1, ATP13A5, HOMEZ, OR6B3, SNX32, THOP1, UROD, TRAF2, MOS, GSTT2B, UGT1A9, TNNT2, FBXW12, CMC4, FMO3, HDC, ADRA2A, DTX2, MT1B, CDKN2C, GABRB3, HBB, MANSC1, UGT1A1, CHRND, GSTT4, GDNF, FKBP1B, PRRC1, SLC25A52, UGT2B4, TTC12, SULT1B1, CASD1, GADL1, DGKD, DUOXA2, TIMP2, AP1S3, NBEAL1, SLC43A1, CASP8, PLCD1, MBOAT1, KRT33B, LONRF2, SPRR2G, HSD17B2, CXXC1, CRTC1, MT-ATP8, PAM, OAT, RETREG2, SLC25A3, CREB3L3, SI, CES2, ITPA, PHEX, ECHDC1, ACP2, DNPH1, PRMT1, NT5C1A, KRT77, ARR3, STT3A, SCYL3, OPRM1, DDO, PHB2, IKBKG, FATE1, CTNNA2, SERPINB6, PRPS1, PRMT5, BDH1, RNF31, HDAC1, PARP9, ZNF148, STRN4, LMOD1, RPIA, MAGEA1, TOB1, ETF1, ITM2B, TPI1, CUTC, RABGGTB, GPX1, CDIPT, PAPOLB, HSP90AA1, GCH1, AMBP, KRT81, IRF1, NAGS, BIK, PADI4, MREG, TSTD1, LGALS1, GOT1, HPRT1, MYRF, OCM, STMN1, GSTA2, KRT34, ENDOU, ADH5, DHX9, AOC2, SCO2, DDX3Y, RANBP3L, RN7SL494P, SNRNP25, KLF12, DLEC1, CCND2, KRTAP2-3, CAB39L, GRWD1, AVEN, TIMM23, GTF2A1, TTC30A, DOLK, YBX2, CYB5R1, ARMCX6, TAF7, SFPQ, WARS2, ALG11, ZDHHC4, TECRL, SRPX, PPP4R2, TEX2, TMEM65, ACTR10, OTUB1, CLN3, COA5, CCNE2, ZMYND15, TSSK4, PTPN20, MEIOC, RPL6, MRTO4, RPS6KC1, MAL2, INTS11, TRMT9B, ENTPD5, SRSF3, TRIM14, TBC1D16, ZNFX1, HDGF, ARID4B, WWP1, NCOA1, GLYATL1, C4ORF46, YWHAE, ALG3, KLHL7, DMAP1, RNASET2, B3GALT1, TP53INP2, STK31, MAPK15, CHCHD4, YWHAG, NEK7, NMNAT3, GOLGA2, VPS26A, TMTC1, NIPA2, UBQLN1, FGFRL1, PRKCH, GPR142, ABCC9, ASB10, AHSG, LMTK2, LANCL1, CXCL14, GRB7, RAB37, SPATA5L1, IRAK4, KIDINS220, FLOT1, NEFH, CHODL, ATP8B1, PXDNL, COL3A1, CLEC16A, GFRA3, IRAK1, FGF11, CD47, TXLNA, CAPS, ENPP1, LRIG2, SOST, PRKCA, SHC3, PRICKLE1, RAB41, TMTC2, DDAH1, COL22A1, ETS1, MOGAT1, PLA2G12A, LIF, CYP21A2, HCK, PNMT, DPYS, PHYKPL, EPHX1, PUDP, PLA2G4C, PCYT1B, CYP7A1, MRPS30, MAGEB4, SULT1C2, NR0B1, POU1F1, ESRRB, LAP3, AS3MT, SERPINB4, ELL, EPGN, OXSM, PRNP, MAFF, OGG1, TFG, EIF3K, KRT23, DPP7, SUMF1, NDST1, PFAS, TAB2, FGF4, OAS2, SHC2, NRL, HAUS3, KRT5, ALDH5A1, HDAC6, PPP4C, TSPAN33, HIST1H2BE, ANKRD33, RIBC2, CEP44, POLR3F, NAP1L4, PUM1, METTL18, TMEM214, ABTB1, TOM1, KRTAP5-5, CEP152, NUDT17, TUBE1, PIP5K1B, CCDC110, HPS4, FASTKD3, PCYT1A, WDR73, ANKRD17, B4GALT7, RBMX2, FBXL3, SERTAD2, MRPS12, HHATL, PEF1, CSNK2A3, DTX4, PPP2R2A, USP25, PELI1, CARNS1, TFEB, PTPN9, GPR15, CPNE1, NEURL1B, TGFB1I1, USO1, LRATD1, XPR1, ARFRP1, DTNB, RFNG, KCNRG, SYT5, PRDX6, RGS11, PLD1, KAZN, MYO1B, PRKCQ, PALLD, NOXA1, SV2A, LMCD1, SPRY1, FGF19, CYS1, MUC12, SLC6A17, A4GNT, EMP2, ADAMTSL1, ANKS4B, ERBB4, ERLEC1, CERS5, RASGEF1B, SLC22A13, PRKAR2B, ARHGAP27, AGA, STON1, MUC6, FES, CD34, CYP2S1, ST6GALNAC3, FKBPL, IDH1, ALKBH3, CDK2, TALDO1, SAMD8, ASPG, PTGFR, G6PD, RPS16, FUOM, ZEB2, CHIC2, SST, ACR, PALM, OLIG2, PIGA, ACY1, TNNC1, POU2F1, CLEC18B |
| cytosol | FBXO22, PSMA8, CCDC8, EIF4E3, NPHP3, NSFL1C, UBE2G1, KATNB1, LARP1B, UCK1, MCM7, CHEK1, RPS27A, ASB7, FANCD2, SNRNP48, GOT1L1, USP47, COPB1, FBXO38, BEX4, BBS4, SDHAF2, PYCR3, KPNA3, PPP2R5D, PSMB2, AGO2, BCL2L15, PDZD11, PHLDA1, ENDOD1, AACS, ARL2BP, ATXN3, NPAS4, DPP9, ARMC8, YAF2, SRP54, AK7, S100A1, RANGRF, NEU2, DIAPH3, FCHO2, GIPC1, SH3GLB1, PLA2G4A, SLK, RAB2A, DLG3, PITPNM2, PKLR, ARHGDIA, LIMD2, KIF21A, RAB27A, AP2B1, NXN, SERF1A, S100A9, PDPN, RGS13, WASHC5, CLC, DRG2, MYO9A, SEC16B, SOCS3, CD1B, MYH9, MUC13, CAPZA1, PAK3, ARHGEF5, LCP1, DOCK6, PREX2, SERPINB9, GNAI2, AHR, GRK5, ARHGAP23, NFKBIB, IREB2, INTS5, KRT16, IL16, HTRA2, PRTN3, TRIO, GLO1, WARS, TGM2, MTHFS, CASZ1, WDR83OS, FBXL20, PIN1, ALDH3A1, AKR1E2, CARS, APC, KRT83, PGK2, AFMID, PFKM, LATS1, GLUL, DCP1B, POLDIP3, METTL21C, MLLT3, XRCC3, SKIV2L, BCDIN3D, SRP68, CDC27, UBE2E1, E2F8, CAPRIN1, RPL27A, GAMT, ING4, ATP6V1G2, FBXO11, NDUFA12, RPEL1, PPP2R5B, HAGH, BAZ2A, POC5, NME7, POLR2C, AGO1, EIF4ENIF1, TMEM87A, POMP, VPS50, HAUS7, CKAP2L, ELP6, CAPZB, RANBP1, CEP85, POLR3K, HBS1L, PFDN2, CEP192, AGO4, C2CD3, ARFGEF2, ARMCX3, PSMC1, NMNAT2, SRSF5, EIF5A2, KSR2, CSNK2B, MCTS1, PAK6, ABI1, PRMT7, MAP3K8, DENND2A, UBE3A, PSMD1, FXR1, MCF2, SMURF2, LYPLA2, BAG1, PDCD6, GOLT1B, SEC61G, PPP1R13L, ATG16L2, CIAO2B, MOV10, KHSRP, SUGT1, TRIM8, DNAJB1, DACT1, NDRG1, BIRC7, ARHGEF10L, ITPKB, HPGDS, DAAM1, PEX5L, GCC2, FERMT2, AAGAB, SBF2, PCP4, YKT6, PDE8B, RAP1GAP, OSBPL9, MTMR4, SEC24D, CPLX2, AZIN2, TRAF3, DNM2, CRMP1, FLRT3, EXOC4, RAB3IP, PIP5KL1, AP1G1, CARD16, SPRY2, RAPGEF4, NOSIP, HOOK1, CEACAM7, SPINK5, PATJ, KIF5A, MYO9B, ZDHHC8, KCNAB2, LDHC, TPT1, HOMEZ, OR6B3, THOP1, UROD, TRAF2, MOS, GSTT2B, TNNT2, FBXW12, HDC, CDKN2C, HBB, CHRND, GSTT4, FKBP1B, SULT1B1, GADL1, DGKD, DUOXA2, AP1S3, NBEAL1, CASP8, KRT33B, SPRR2G, CXXC1, CRTC1, CREB3L3, ITPA, ECHDC1, DNPH1, PRMT1, NT5C1A, KRT77, DDO, IKBKG, CTNNA2, SERPINB6, PRPS1, PRMT5, RNF31, HDAC1, PARP9, LMOD1, RPIA, ETF1, TPI1, CUTC, RABGGTB, GPX1, HSP90AA1, GCH1, AMBP, KRT81, IRF1, PADI4, TSTD1, GOT1, HPRT1, MYRF, STMN1, GSTA2, KRT34, ADH5, DHX9, DDX3Y, SNRNP25, KLF12, DLEC1, CCND2, KRTAP2-3, CAB39L, GRWD1, AVEN, GTF2A1, CYB5R1, SFPQ, ACTR10, OTUB1, CLN3, CCNE2, RPL6, INTS11, TBC1D16, ARID4B, WWP1, NCOA1, YWHAE, KLHL7, DMAP1, TP53INP2, YWHAG, VPS26A, UBQLN1, PRKCH, GPR142, ASB10, LMTK2, GRB7, IRAK4, KIDINS220, CHODL, ATP8B1, CLEC16A, GFRA3, IRAK1, TXLNA, CAPS, PRKCA, SHC3, PRICKLE1, RAB41, DDAH1, LIF, HCK, PNMT, DPYS, PUDP, PLA2G4C, SULT1C2, POU1F1, ESRRB, AS3MT, SERPINB4, ELL, OXSM, PRNP, OGG1, TFG, EIF3K, KRT23, PFAS, TAB2, OAS2, SHC2, NRL, HAUS3, KRT5, HDAC6, PPP4C, HIST1H2BE, ANKRD33, POLR3F, PUM1, METTL18, TMEM214, ABTB1, TOM1, KRTAP5-5, CEP152, PIP5K1B, CCDC110, HPS4, PCYT1A, WDR73, FBXL3, SERTAD2, CSNK2A3, DTX4, PPP2R2A, USP25, PELI1, CARNS1, TFEB, CPNE1, NEURL1B, TGFB1I1, USO1, ARFRP1, PRDX6, KAZN, PRKCQ, PALLD, NOXA1, SPRY1, CYS1, EMP2, ERBB4, PRKAR2B, ARHGAP27, FES, FKBPL, IDH1, ALKBH3, CDK2, TALDO1, SAMD8, ASPG, G6PD, RPS16, FUOM, ZEB2, ACY1, TNNC1 |
| endomembrane system | SGCD, ALG10B, RPGR, SPEF2, NSFL1C, F8A1, PIGH, RPS27A, COPB1, SSR4, KPNA3, FKBP9, SERINC3, ATXN3, ARMC8, S100A11, MIA2, SRP54, FER1L5, SNX4, S100A1, RANGRF, CTSZ, ORMDL3, OR51E2, FCHO2, GIPC1, SH3GLB1, IL8RB, CD59, PLA2G4A, RAB2A, ST3GAL4, PITPNM2, COL4A2, TARM1, CPNE6, PKDCC, MYMK, RAB27A, SLC9A3R2, AP2B1, SLC9A3, MUC16, CNR2, S100A9, COL13A1, WASHC5, CD93, CAPN8, SEC16B, CD1B, ZPBP2, MYH9, MUC13, CNST, SNAP23, SMPD4, BST1, GIMAP5, COL1A2, CYP3A43, RAB40A, RDH5, SERPINB9, CHST13, FCGR2A, GNAI2, GRK5, A1BG, NPY, GALNT15, INTS5, SELENOF, HTRA2, PRTN3, TGM2, HEBP2, METTL7B, WDR83OS, HP, SERPING1, MOXD1, ALDH3A1, APC, TMED5, CD177, TMBIM6, GLUL, ZDHHC18, RGPD3, SRP68, INTS1, THAP7, RPL27A, MARF1, RNF186, ATP6V1G2, DYSF, CREB3L2, PNPLA7, CYB5R4, RANBP3, TMEM87A, IQCF1, POMP, VPS50, PIGT, EMC2, RP2, RANBP1, TPST2, CEP85, ARFGEF2, TMX3, NMNAT2, EIF5A2, CSNK2B, ABI1, ELOVL1, PSMD1, SH3BGRL2, FXR1, LYPLA2, PDCD6, GOLT1B, SEC61G, TSSK1B, NDRG1, BIRC7, ESYT1, SSR2, BOK, ITPKB, CCDC170, EPN3, JPH4, CAV3, GCC2, MBOAT4, AKAP6, KCNK9, SBF2, YKT6, OMD, RAP1GAP, ECM1, ACKR3, CAPN11, GUCY2C, OSBPL9, ANGPTL3, FN1, MTMR4, SEC24D, MACO1, RAB8B, SLC15A3, AZIN2, TRAF3, MCEMP1, DNM2, TLR3, FLRT3, SLC9A6, SERPINE1, RAB3IP, EPHA5, RTN4R, AP1G1, RYR3, CHRNB4, BGN, NDST2, NOSIP, SPINK5, PGAP1, DPEP3, TMEM230, ITGA1, JAM3, RAB33B, CEACAM1, LOXL2, NUS1, ZDHHC8, KCNAB2, FADS2, WSCD1, TPT1, FMO2, HTR7, ATP13A5, SNX32, TRAF2, UGT1A9, FMO3, DTX2, HBB, MANSC1, UGT1A1, GDNF, FKBP1B, PRRC1, UGT2B4, CASD1, DGKD, DUOXA2, TIMP2, AP1S3, SLC43A1, MBOAT1, HSD17B2, PAM, RETREG2, CREB3L3, SI, CES2, PHEX, STT3A, SCYL3, OPRM1, FATE1, SERPINB6, PRMT5, ZNF148, ITM2B, CDIPT, HSP90AA1, GCH1, AMBP, BIK, MREG, LGALS1, MYRF, RANBP3L, CCND2, AVEN, DOLK, CYB5R1, NUP210L, ALG11, ZDHHC4, TECRL, SRPX, TEX2, ACTR10, CLN3, TSSK4, RPL6, RPS6KC1, MAL2, ENTPD5, TBC1D16, ALG3, RNASET2, B3GALT1, STK31, MAPK15, GOLGA2, VPS26A, TMTC1, NIPA2, UBQLN1, FGFRL1, AHSG, LMTK2, CXCL14, RAB37, IRAK4, KIDINS220, FLOT1, CHODL, ATP8B1, PXDNL, COL3A1, CLEC16A, IRAK1, CD47, SOST, PRKCA, PRICKLE1, RAB41, TMTC2, COL22A1, MOGAT1, CYP21A2, HCK, TEX28, EPHX1, PLA2G4C, PCYT1B, CYP7A1, PRNP, TFG, DPP7, SUMF1, NDST1, TAB2, HDAC6, TSPAN33, TMEM214, TOM1, PIP5K1B, HPS4, RBM15B, PCYT1A, ANKRD17, B4GALT7, RBMX2, HHATL, PEF1, USP25, GPR15, CPNE1, NEURL1B, USO1, XPR1, ARFRP1, RFNG, KCNRG, SYT5, PRDX6, PLD1, MYO1B, SV2A, SPRY1, MUC12, SLC6A17, A4GNT, EMP2, ADAMTSL1, ANKS4B, ERLEC1, CERS5, RASGEF1B, SLC22A13, AGA, STON1, MUC6, FES, CYP2S1, ST6GALNAC3, IDH1, CDK2, SAMD8, CHIC2, SST, ACR, PIGA, POU2F1, CLEC18B |
| membrane | TIMM8A, SGCD, CIZ1, CCDC8, ALG10B, UTP20, KATNB1, TMEM234, RGL4, ACBD4, EXO1, MCM7, PIGH, DNAJC30, PCDHGC4, RPS27A, CATSPER2, PRPF19, SLC25A14, COPB1, SSR4, BBS4, CHDH, SDR42E1, HIC2, PSMB2, AGO2, GPR148, SERINC3, PDZD11, TMIGD3, ENDOD1, ATXN3, NKD1, MIA2, HSDL2, MTFP1, PLET1, MYADM, FER1L5, SNX4, CCR7, RANGRF, NEU2, NPBWR1, CYSLTR2, KCNMB1, CTSZ, ORMDL3, OR51E2, FCHO2, TRPM5, GIPC1, CYSLTR1, KCNMB3, TNFSF9, SH3GLB1, IL8RB, CD59, KCNAB3, KCNG4, PLA2G4A, OR2W3, HCRTR1, RAB2A, DLG3, CSF1R, SLC7A9, TSPAN4, ST3GAL4, PITPNM2, COL4A2, TARM1, NLGN3, CPNE6, ARHGDIA, TMEM215, MYMK, MARCKSL1, LIMD2, KIF21A, SEMA4D, RAB27A, SLC9A3R2, AP2B1, SLC9A3, MUC16, CNR2, SERF1A, HRH4, S100A9, COL13A1, PDPN, RGS13, WASHC5, CD93, DRG2, MYO9A, VASN, SEC16B, SLC25A27, SOCS3, CD1B, SLC4A1AP, CLIC6, OXER1, MYH9, SLC10A2, MUC13, GJB3, CNST, SEMA6B, NPC1L1, SNAP23, SMPD4, BST1, PAK3, ARHGEF5, LCP1, CXCL16, CNR1, GIMAP5, PREX2, SVOPL, LMBRD2, CYP3A43, RAB40A, RDH5, GYPB, SERPINB9, CHST13, CYP27C1, OR52H1, HTR3A, FCGR2A, GNAI2, IGHV3-23, GRK5, OR2L13, IGLC2, FCER2, IL18RAP, PLCD3, TAS2R19, MS4A4E, GREB1L, DMAC2, ITFG1, EVC, A1BG, RFTN2, TRBV12-5, GALNT15, INTS5, KRT16, IL16, DYTN, POLE, DNAJC22, SLC43A3, AIDA, HTRA2, PRTN3, TRIO, MT-CO1, EFS, GLO1, TGM2, METTL7B, WDR83OS, SLC6A1, ABCC10, MOXD1, ALDH3A1, APC, TMED5, RHBDD3, CD177, PFKM, POTED, TMBIM6, GLUL, NDUFB3, ZDHHC18, DCP1B, CDC42SE1, INTS1, THAP7, CDC27, COX14, CAPRIN1, MRPL22, RPL27A, MARF1, RNF186, ATP6V1G2, DYSF, NDUFA12, CREB3L2, PNPLA7, CYB5R4, MRPS18B, MRPL39, HEATR1, EIF4ENIF1, TMEM87A, POMP, VPS50, PIGT, HAUS7, EMC2, RP2, ELP6, CAPZB, ST7, TPST2, HBS1L, CPO, AGO4, NDUFA11, MIEF2, ARFGEF2, TMX3, ARMCX3, MTG1, RECQL, PSMC1, NMNAT2, EIF5A2, KSR2, CSNK2B, ELOVL1, PSMD1, SH3BGRL2, FXR1, MCF2, SMURF2, BAG1, PDCD6, GOLT1B, SEC61G, ATG16L2, MOSPD3, KHSRP, DAP3, NDRG1, ESYT1, TRAP1, SSR2, BOK, SAMM50, SLC36A4, BCS1L, ITPKB, SDK1, DAAM1, PEX5L, EPN3, JPH4, CAV3, GCC2, MBOAT4, FERMT2, AKAP6, PTPRT, ASIC3, KCNK9, SBF2, PROCR, YKT6, CXCL9, GPR6, ANO3, UNC5B, RAP1GAP, ACKR3, TRIL, GUCY2C, OSBPL9, FN1, PAG1, MTMR4, SEC24D, MACO1, OSMR, CPLX2, RAB8B, ASPM, EPHA1, SLC15A3, AZIN2, TRAF3, MCEMP1, DNM2, TLR3, FLRT3, SLC9A6, EXOC4, SERPINE1, TRPV2, LUZP1, PIP5KL1, EPHA5, RTN4R, SSH1, SMAGP, ODR4, AP1G1, ATP2B2, RYR3, SPRY2, GPR34, CHRNB4, PLPPR4, GPR171, EFNB2, BGN, SLC9A2, IL17RE, RIC8A, NDST2, IGF1R, CLDN12, RAPGEF4, NOSIP, HOOK1, SCARF2, CEACAM7, PCDH1, FCN3, SLC26A2, SPINK5, PATJ, PGAP1, DPEP3, TMEM230, ITGA1, JAM3, KIF5A, RAB33B, CACNA1I, CEACAM1, LOXL2, FLRT1, MYO9B, NUS1, ZDHHC8, HCN2, KCNAB2, FADS2, PCDHA8, CD79B, OR56A5, C5ORF15, IGHV6-1, WSCD1, LDHC, FMO2, HTR7, UCP1, ATP13A5, CD163L1, OR6B3, IFITM10, TRAF2, ABCA10, UGT1A9, OR2L8, FMO3, OR11L1, ADRA2A, DTX2, AADACL3, GABRB3, MANSC1, UGT1A1, IGHV3-48, CHRND, TRBJ2-3, FKBP1B, TRDV2, TAS2R31, TMEM94, TRGV1, SLC25A52, UGT2B4, CASD1, DGKD, DUOXA2, AP1S3, NBEAL1, SLC43A1, CASP8, PLCD1, MBOAT1, GFI1B, SPRR2G, HSD17B2, XKR4, VSIG10L, SLC17A4, CRTC1, MT-ATP8, PAM, RETREG2, SLC25A3, CREB3L3, SI, PHEX, ECHDC1, ACP2, KRT77, STT3A, SCYL3, OPRM1, PHB2, FATE1, CTNNA2, SERPINB6, PRPS1, BDH1, RNF31, PARP9, STRN4, LMOD1, MAGEA1, ITM2B, RABGGTB, CDIPT, HSP90AA1, GCH1, AMBP, BIK, MREG, DDX18, MYRF, TMEM200B, STMN1, ENDOU, DHX9, ARID4A, AOC2, SCO2, DDX3Y, IGKV1-27, TMEM265, CCND2, AVEN, TIMM23, DOLK, CYB5R1, ARMCX6, RAD50, NUP210L, WARS2, ALG11, ZDHHC4, TECRL, SRPX, TEX2, TMEM65, CLN3, PTPN20, NOL9, RPL6, RPS6KC1, MAL2, PSD4, ENTPD5, TRIM14, ZNFX1, SMIM30, WWP1, NCOA1, YWHAE, ALG3, KLHL7, B3GALT1, YWHAG, GOLGA2, SLC27A6, GNG2, VPS26A, TMTC1, NIPA2, UBQLN1, KCNE4, BEST4, FGFRL1, PRKCH, GPR142, ABCC9, GPR87, BTN3A3, LMTK2, PLXDC2, GFRAL, LANCL1, GRB7, CDH26, RTP2, RAB37, FXYD6, SLC2A9, SIGLEC8, F2RL3, KCNQ4, IRAK4, KIDINS220, FLOT1, CHODL, ATP8B1, PXDNL, CLEC16A, GFRA3, SLCO1B1, IRAK1, EFNA2, CD47, TXLNA, CAPS, ENPP1, LRIG2, PRKCA, ITGA11, SHC3, PRICKLE1, RAB41, BEST3, GPM6A, SLC12A1, SLC8A2, TMTC2, FFAR1, GNG10, MOGAT1, OR2AP1, CYP21A2, HCK, IGKV3-20, TEX28, IGLV3-9, RRP22, TMEM154, IGLV3-32, EPHX1, SLC22A24, PLA2G4C, PCYT1B, CYP7A1, MRPS30, EMILIN1, TAS2R60, NR0B1, SLC6A2, SERPINB4, MCHR1, EPGN, PRNP, EIF3K, HEG1, NDST1, TAB2, IGKV1-8, OAS2, SHC2, KRT5, HDAC6, PPP4C, TSPAN33, TMEM214, ABTB1, TOM1, NAT10, PIP5K1B, HPS4, PCYT1A, WDR73, ANKRD17, B4GALT7, RBMX2, MRPS12, HHATL, PEF1, PRMT8, TFEB, GPR15, CPNE1, AMIGO2, USO1, XPR1, ARFRP1, DTNB, RFNG, SYT5, PRDX6, RGS11, PLPPR3, PLD1, KCNK12, NCR3, KAZN, MYO1B, PRKCQ, PALLD, SLCO5A1, NOXA1, SV2A, PILRA, PLPPR2, SPRY1, PCDH12, CYS1, MUC12, SLC6A17, DLGAP1, A4GNT, EMP2, ADAMTSL1, ANKS4B, ERBB4, SLC26A1, POPDC2, CERS5, EDA2R, RASGEF1B, SLC19A1, TMEM204, SLC22A13, FAM171A2, SLC45A1, PRKAR2B, IGKV1-37, C4BPB, ARHGAP27, STON1, MUC6, FES, C8G, FOLR2, CD34, CYP2S1, ST6GALNAC3, SAMD8, PTGFR, SERINC4, G6PD, RPS16, CHIC2, PALM, PIGA, TMEM220 |
| extracellular region | PSMA8, NPHP3, SPEF2, UBE2G1, CHEK1, RPS27A, SSR4, PSMB2, AGO2, MTMR11, PDZD11, ENDOD1, ARMC8, S100A11, IGSF10, SPATA6, PLET1, SVBP, S100A1, GKN2, ASPN, CTSZ, GIPC1, TNFSF9, CD59, XCL2, SLK, RAB2A, DLG3, ST3GAL4, COL4A2, CPNE6, EFEMP2, MFAP5, TSLP, PKLR, ARHGDIA, CCL17, PKDCC, MARCKSL1, PCYOX1, SEMA4D, RAB27A, SLC9A3R2, SLC9A3, MUC16, S100A9, COL13A1, CCL20, OTOS, VASN, WFDC12, CD1B, LGI2, CLIC6, ZPBP2, LCN1, MYH9, MUC13, SNAP23, CAPZA1, BST1, LCP1, CXCL16, AMY1A, COL1A2, SCGB1C1, SERPINB9, GNAI2, IGHV3-23, IGLC2, FCER2, SBSPON, ARHGAP23, ITFG1, IFNA13, TSHB, A1BG, ZNF649, NPY, KRT16, IL16, PRTN3, NPTX2, GLO1, WARS, TGM2, HEBP2, HP, SERPING1, MOXD1, ALDH3A1, KRT83, PGK2, CD177, SCGB1D4, GLUL, MLLT3, TUBB1, DYSF, DDB1, VPS50, RP2, CAPZB, HBS1L, CPO, KIF27, CLPSL2, JADE2, CSNK2B, ABI1, PSMD1, LYPLA2, PDCD6, MOV10, DNAJB1, NDRG1, SAMM50, EPN3, LTBP2, GREM2, ANGPTL5, FGF16, PROCR, CXCL9, OMD, ECM1, TRIL, ANGPTL3, FN1, MTMR4, PLA2G1B, OLFML2A, APLN, RAB8B, RSPO3, DNM2, MMRN2, PDGFRL, TLR3, FLRT3, SERPINE1, LUZP1, RTN4R, ATP2B2, MFAP4, INSL5, BGN, THBS3, WNT16, FDCSP, IL17RE, WFIKKN1, C1QTNF7, CEACAM7, OLFML3, FCN3, SLC26A2, SPINK5, PATJ, ITGA1, JAM3, LYG1, CEACAM1, LOXL2, DEFB104A, CRTAC1, CGB5, CD79B, GBP6, CGB8, IGHV6-1, LDHC, TPT1, CD163L1, GSTT2B, OR11L1, PRSS53, HBB, IGHV3-48, GDNF, C14ORF93, TIMP2, PLCD1, KRT33B, PAM, SLC25A3, SI, ACP2, DNPH1, KRT77, SERPINB6, PM20D1, GPX5, ITM2B, TPI1, HSP90AA1, AMBP, KRT81, CTF1, LGALS1, GOT1, HPRT1, C1ORF56, STMN1, GSTA2, KRT34, ENDOU, ADH5, IGKV1-27, DEFB121, DHRSX, CYB5R1, ACTR10, OTUB1, MAL2, INTS11, ENTPD5, HDGF, WWP1, YWHAE, RNASET2, NCOA5, MAPK15, YWHAG, GNG2, ADAMTS12, NXPE4, PRKCH, AHSG, ADAMTSL3, CXCL14, F2RL3, IRAK4, FLOT1, PXDNL, COL3A1, VWA2, LGI4, CD47, TXLNA, SEMA3A, ENPP1, LRIG2, SOST, PRKCA, CSN3, GPM6A, SLC12A1, RBP4, DDAH1, COL22A1, PLA2G12A, LIF, IGKV3-20, IGLV3-9, CORT, IGLV3-32, DPYS, PYY, EMILIN1, PRH1, LAP3, SCT, SERPINB4, EPGN, PRNP, HEG1, DPP7, PFAS, FGF4, DPT, IGKV1-8, KRT5, HIST1H2BE, TOM1, PEF1, CPNE1, RFNG, PRDX6, CXCL6, MYO1B, PILRA, PCDH12, FGF19, ADAMTSL1, ERBB4, METRNL, SLC22A13, PRKAR2B, LCN9, IGKV1-37, C4BPB, AGA, MUC6, C8G, FOLR2, CD34, FKBPL, IDH1, CPN1, TALDO1, PTGFR, G6PD, RPS16, SST, ACR, ACY1, CLEC18B |
| extracellular space | PSMA8, UBE2G1, CHEK1, RPS27A, SSR4, PSMB2, AGO2, MTMR11, ENDOD1, S100A11, GKN2, ASPN, CTSZ, GIPC1, TNFSF9, CD59, XCL2, SLK, RAB2A, DLG3, COL4A2, CPNE6, EFEMP2, TSLP, PKLR, ARHGDIA, CCL17, MARCKSL1, PCYOX1, SEMA4D, RAB27A, SLC9A3R2, SLC9A3, MUC16, S100A9, COL13A1, CCL20, VASN, CD1B, CLIC6, LCN1, MYH9, MUC13, SNAP23, CAPZA1, BST1, LCP1, CXCL16, AMY1A, COL1A2, SERPINB9, GNAI2, IGHV3-23, IGLC2, FCER2, ARHGAP23, ITFG1, IFNA13, TSHB, A1BG, ZNF649, NPY, KRT16, IL16, PRTN3, GLO1, WARS, TGM2, HEBP2, HP, SERPING1, MOXD1, ALDH3A1, KRT83, PGK2, CD177, GLUL, MLLT3, TUBB1, DYSF, DDB1, VPS50, RP2, CAPZB, HBS1L, CPO, JADE2, CSNK2B, ABI1, LYPLA2, PDCD6, MOV10, DNAJB1, NDRG1, SAMM50, EPN3, LTBP2, GREM2, ANGPTL5, FGF16, PROCR, CXCL9, OMD, ECM1, TRIL, ANGPTL3, FN1, MTMR4, PLA2G1B, OLFML2A, APLN, RAB8B, DNM2, MMRN2, TLR3, FLRT3, SERPINE1, LUZP1, RTN4R, ATP2B2, MFAP4, BGN, WNT16, WFIKKN1, OLFML3, FCN3, SLC26A2, PATJ, ITGA1, JAM3, CEACAM1, LOXL2, CRTAC1, CGB5, CD79B, GBP6, CGB8, LDHC, TPT1, GSTT2B, OR11L1, HBB, GDNF, TIMP2, PLCD1, KRT33B, PAM, SLC25A3, SI, ACP2, DNPH1, KRT77, SERPINB6, PM20D1, ITM2B, TPI1, HSP90AA1, AMBP, KRT81, CTF1, LGALS1, GOT1, HPRT1, STMN1, GSTA2, KRT34, ENDOU, ADH5, IGKV1-27, CYB5R1, OTUB1, MAL2, INTS11, ENTPD5, HDGF, WWP1, YWHAE, RNASET2, NCOA5, YWHAG, GNG2, NXPE4, PRKCH, AHSG, CXCL14, IRAK4, FLOT1, PXDNL, COL3A1, VWA2, LGI4, CD47, ENPP1, LRIG2, SOST, PRKCA, CSN3, GPM6A, SLC12A1, RBP4, DDAH1, COL22A1, LIF, IGKV3-20, IGLV3-9, CORT, IGLV3-32, DPYS, EMILIN1, PRH1, LAP3, SCT, SERPINB4, EPGN, PRNP, DPP7, PFAS, FGF4, DPT, IGKV1-8, KRT5, HIST1H2BE, TOM1, PEF1, CPNE1, PRDX6, CXCL6, MYO1B, PILRA, PCDH12, FGF19, METRNL, SLC22A13, PRKAR2B, LCN9, IGKV1-37, C4BPB, AGA, MUC6, C8G, IDH1, CPN1, TALDO1, G6PD, RPS16, SST, ACY1, CLEC18B |
| cell periphery | SGCD, CIZ1, CCDC8, ALG10B, UTP20, KATNB1, RGL4, EXO1, PCDHGC4, RPS27A, CATSPER2, SLC25A14, COPB1, BBS4, HIC2, GPR148, SERINC3, PDZD11, TMIGD3, ATXN3, NKD1, PLET1, MYADM, FER1L5, SNX4, CCR7, RANGRF, NPBWR1, CYSLTR2, ASPN, KCNMB1, CTSZ, ORMDL3, OR51E2, FCHO2, TRPM5, GIPC1, CYSLTR1, KCNMB3, TNFSF9, IL8RB, CD59, KCNAB3, KCNG4, OR2W3, HCRTR1, DLG3, CSF1R, SLC7A9, TSPAN4, COL4A2, TARM1, NLGN3, CPNE6, EFEMP2, MFAP5, ARHGDIA, MYMK, MARCKSL1, LIMD2, KIF21A, SEMA4D, RAB27A, SLC9A3R2, AP2B1, SLC9A3, MUC16, CNR2, HRH4, S100A9, COL13A1, PDPN, RGS13, CD93, CLC, VASN, SOCS3, CD1B, SLC4A1AP, CLIC6, OXER1, MYH9, SLC10A2, MUC13, GJB3, CNST, SEMA6B, NPC1L1, SNAP23, SMPD4, BST1, PAK3, ARHGEF5, LCP1, CXCL16, CNR1, PREX2, LMBRD2, COL1A2, RAB40A, GYPB, SERPINB9, OR52H1, HTR3A, FCGR2A, GNAI2, IGHV3-23, GRK5, OR2L13, IGLC2, FCER2, SBSPON, IL18RAP, PLCD3, TAS2R19, ITFG1, EVC, A1BG, RFTN2, TRBV12-5, KRT16, IL16, DYTN, POLE, SLC43A3, HTRA2, PRTN3, EFS, GLO1, TGM2, SERPING1, SLC6A1, ABCC10, ALDH3A1, APC, CD177, PFKM, POTED, TMBIM6, GLUL, CDC42SE1, DYSF, HAUS7, RP2, CAPZB, CPO, TMX3, KSR2, CSNK2B, SMURF2, NDRG1, ESYT1, TRAP1, SLC36A4, SDK1, DAAM1, EPN3, JPH4, CAV3, LTBP2, FERMT2, AKAP6, PTPRT, ASIC3, KCNK9, ANGPTL5, PROCR, YKT6, CXCL9, GPR6, ANO3, UNC5B, OMD, ECM1, ACKR3, TRIL, GUCY2C, ANGPTL3, FN1, PAG1, OSMR, OLFML2A, RAB8B, ASPM, EPHA1, TRAF3, MCEMP1, DNM2, MMRN2, TLR3, FLRT3, SLC9A6, EXOC4, SERPINE1, TRPV2, EPHA5, RTN4R, SSH1, SMAGP, AP1G1, ATP2B2, MFAP4, RYR3, SPRY2, GPR34, CHRNB4, PLPPR4, GPR171, EFNB2, BGN, SLC9A2, THBS3, IL17RE, RIC8A, IGF1R, CLDN12, RAPGEF4, CEACAM7, PCDH1, FCN3, SLC26A2, SPINK5, PATJ, DPEP3, ITGA1, JAM3, CACNA1I, CEACAM1, LOXL2, MYO9B, HCN2, KCNAB2, FADS2, PCDHA8, CD79B, OR56A5, IGHV6-1, HTR7, ATP13A5, CD163L1, OR6B3, IFITM10, TRAF2, OR2L8, OR11L1, ADRA2A, GABRB3, UGT1A1, IGHV3-48, CHRND, TRBJ2-3, TRDV2, TAS2R31, TRGV1, DGKD, DUOXA2, TIMP2, AP1S3, SLC43A1, CASP8, PLCD1, GFI1B, SPRR2G, XKR4, SLC17A4, CRTC1, SLC25A3, SI, PHEX, KRT77, OPRM1, PHB2, CTNNA2, SERPINB6, RNF31, MAGEA1, ITM2B, RABGGTB, CDIPT, HSP90AA1, AMBP, MREG, LGALS1, ENDOU, ARID4A, AOC2, IGKV1-27, CYB5R1, WARS2, ZDHHC4, SRPX, TMEM65, CLN3, PTPN20, MAL2, PSD4, HDGF, WWP1, NCOA1, YWHAE, KLHL7, SLC27A6, GNG2, ADAMTS12, NIPA2, UBQLN1, KCNE4, BEST4, FGFRL1, PRKCH, GPR142, ABCC9, GPR87, BTN3A3, AHSG, ADAMTSL3, GFRAL, LANCL1, GRB7, CDH26, RTP2, RAB37, FXYD6, SLC2A9, SIGLEC8, F2RL3, KCNQ4, IRAK4, FLOT1, ATP8B1, PXDNL, COL3A1, VWA2, GFRA3, SLCO1B1, IRAK1, EFNA2, CD47, CAPS, ENPP1, LRIG2, SOST, PRKCA, ITGA11, SHC3, RAB41, BEST3, GPM6A, SLC12A1, SLC8A2, FFAR1, COL22A1, GNG10, OR2AP1, HCK, IGKV3-20, IGLV3-9, RRP22, IGLV3-32, SLC22A24, PLA2G4C, EMILIN1, TAS2R60, SLC6A2, SERPINB4, MCHR1, EPGN, PRNP, HEG1, TAB2, DPT, IGKV1-8, SHC2, HDAC6, PPP4C, TSPAN33, ABTB1, TOM1, PIP5K1B, WDR73, PRMT8, GPR15, CPNE1, TGFB1I1, AMIGO2, XPR1, DTNB, RGS11, PLD1, NCR3, KAZN, MYO1B, PRKCQ, PALLD, SLCO5A1, NOXA1, SV2A, PILRA, SPRY1, PCDH12, CYS1, MUC12, SLC6A17, DLGAP1, EMP2, ADAMTSL1, ANKS4B, ERBB4, SLC26A1, POPDC2, EDA2R, RASGEF1B, SLC19A1, TMEM204, SLC22A13, PRKAR2B, IGKV1-37, C4BPB, STON1, MUC6, FES, C8G, C17ORF58, FOLR2, CD34, PTGFR, G6PD, CHIC2, PALM |
| vesicle | PSMA8, UBE2G1, F8A1, RGL4, RPS27A, COPB1, SSR4, PSMB2, AGO2, MTMR11, PHLDA1, ENDOD1, ARMC8, S100A11, FER1L5, SNX4, CTSZ, ORMDL3, OR51E2, FCHO2, GIPC1, SH3GLB1, IL8RB, CD59, SLK, RAB2A, TSPAN4, COL4A2, TARM1, NLGN3, CPNE6, EFEMP2, PKLR, ARHGDIA, MARCKSL1, PCYOX1, RAB27A, SLC9A3R2, AP2B1, SLC9A3, MUC16, S100A9, PDPN, WASHC5, CD93, VASN, SEC16B, CD1B, CLIC6, ZPBP2, MYH9, CNST, NPC1L1, SNAP23, CAPZA1, BST1, LCP1, GIMAP5, AMY1A, COL1A2, RAB40A, SERPINB9, FCGR2A, GNAI2, IGHV3-23, IGLC2, FCER2, ARHGAP23, ITFG1, A1BG, NPY, GALNT15, KRT16, PRTN3, GLO1, WARS, TGM2, HEBP2, HP, SERPING1, MOXD1, TMED5, PGK2, CD177, GLUL, MLLT3, TUBB1, ATP6V1G2, DYSF, IQCF1, DDB1, VPS50, PIGT, RP2, CAPZB, HBS1L, ARFGEF2, TMX3, NMNAT2, JADE2, CSNK2B, ABI1, PSMD1, LYPLA2, PDCD6, TSSK1B, DNAJB1, NDRG1, BOK, SAMM50, EPN3, CAV3, LTBP2, KCNK9, SBF2, PROCR, YKT6, OMD, RAP1GAP, ECM1, ACKR3, CAPN11, OSBPL9, ANGPTL3, FN1, MTMR4, SEC24D, RAB8B, SLC15A3, AZIN2, TRAF3, MCEMP1, DNM2, MMRN2, TLR3, SLC9A6, SERPINE1, RAB3IP, TRPV2, LUZP1, RTN4R, SMAGP, AP1G1, ATP2B2, CHRNB4, BGN, OLFML3, SLC26A2, SPINK5, PATJ, DPEP3, TMEM230, ITGA1, RAB33B, CEACAM1, CRTAC1, KCNAB2, CD79B, GBP6, LDHC, TPT1, ATP13A5, SNX32, TRAF2, GSTT2B, OR11L1, GABRB3, HBB, DGKD, TIMP2, AP1S3, PLCD1, KRT33B, PAM, SLC25A3, SI, ACP2, DNPH1, KRT77, OPRM1, SERPINB6, PM20D1, ITM2B, TPI1, HSP90AA1, GCH1, AMBP, MREG, LGALS1, GOT1, HPRT1, STMN1, GSTA2, ADH5, CYB5R1, ACTR10, OTUB1, CLN3, TSSK4, RPS6KC1, MAL2, TRIM14, TBC1D16, WWP1, YWHAE, RNASET2, TP53INP2, STK31, MAPK15, YWHAG, GOLGA2, GNG2, VPS26A, NXPE4, NIPA2, UBQLN1, FGFRL1, PRKCH, AHSG, LMTK2, RAB37, IRAK4, KIDINS220, FLOT1, CLEC16A, VWA2, IRAK1, CD47, CAPS, LRIG2, PRKCA, GPM6A, SLC12A1, RBP4, DDAH1, HCK, IGKV3-20, DPYS, EMILIN1, LAP3, EPGN, PRNP, DPP7, PFAS, TAB2, KRT5, HDAC6, HIST1H2BE, TOM1, HPS4, PEF1, GPR15, CPNE1, NEURL1B, USO1, SYT5, PRDX6, PLD1, MYO1B, SV2A, PILRA, PCDH12, SLC6A17, EMP2, METRNL, RASGEF1B, SLC22A13, PRKAR2B, AGA, STON1, FES, C8G, IDH1, CDK2, TALDO1, G6PD, RPS16, CHIC2, SST, ACR, PALM, ACY1, CLEC18B |
| plasma membrane | SGCD, CIZ1, CCDC8, ALG10B, UTP20, KATNB1, RGL4, EXO1, PCDHGC4, RPS27A, CATSPER2, SLC25A14, COPB1, BBS4, HIC2, GPR148, SERINC3, PDZD11, TMIGD3, ATXN3, NKD1, PLET1, MYADM, FER1L5, SNX4, CCR7, RANGRF, NPBWR1, CYSLTR2, KCNMB1, CTSZ, ORMDL3, OR51E2, FCHO2, TRPM5, CYSLTR1, KCNMB3, TNFSF9, IL8RB, CD59, KCNAB3, KCNG4, OR2W3, HCRTR1, DLG3, CSF1R, SLC7A9, TSPAN4, TARM1, NLGN3, CPNE6, ARHGDIA, MYMK, MARCKSL1, LIMD2, KIF21A, SEMA4D, RAB27A, SLC9A3R2, AP2B1, SLC9A3, MUC16, CNR2, HRH4, S100A9, COL13A1, PDPN, RGS13, CD93, VASN, SOCS3, CD1B, SLC4A1AP, CLIC6, OXER1, MYH9, SLC10A2, MUC13, GJB3, CNST, SEMA6B, NPC1L1, SNAP23, SMPD4, BST1, PAK3, ARHGEF5, LCP1, CXCL16, CNR1, PREX2, LMBRD2, RAB40A, GYPB, OR52H1, HTR3A, FCGR2A, GNAI2, IGHV3-23, GRK5, OR2L13, IGLC2, FCER2, IL18RAP, PLCD3, TAS2R19, ITFG1, EVC, A1BG, RFTN2, TRBV12-5, KRT16, IL16, DYTN, POLE, SLC43A3, HTRA2, PRTN3, EFS, GLO1, TGM2, SLC6A1, ABCC10, ALDH3A1, APC, CD177, PFKM, POTED, TMBIM6, GLUL, CDC42SE1, DYSF, HAUS7, RP2, CPO, TMX3, KSR2, CSNK2B, SMURF2, NDRG1, ESYT1, SLC36A4, SDK1, DAAM1, EPN3, JPH4, CAV3, FERMT2, AKAP6, PTPRT, ASIC3, KCNK9, PROCR, YKT6, CXCL9, GPR6, ANO3, UNC5B, ACKR3, GUCY2C, FN1, PAG1, OSMR, RAB8B, ASPM, EPHA1, TRAF3, MCEMP1, DNM2, TLR3, FLRT3, SLC9A6, EXOC4, SERPINE1, TRPV2, EPHA5, RTN4R, SSH1, SMAGP, AP1G1, ATP2B2, RYR3, SPRY2, GPR34, CHRNB4, PLPPR4, GPR171, EFNB2, BGN, SLC9A2, IL17RE, RIC8A, IGF1R, CLDN12, RAPGEF4, CEACAM7, PCDH1, FCN3, SLC26A2, PATJ, DPEP3, ITGA1, JAM3, CACNA1I, CEACAM1, HCN2, KCNAB2, FADS2, PCDHA8, CD79B, OR56A5, IGHV6-1, HTR7, ATP13A5, CD163L1, OR6B3, IFITM10, TRAF2, OR2L8, OR11L1, ADRA2A, GABRB3, UGT1A1, IGHV3-48, CHRND, TRBJ2-3, TRDV2, TAS2R31, TRGV1, DGKD, DUOXA2, AP1S3, SLC43A1, CASP8, PLCD1, GFI1B, SPRR2G, XKR4, SLC17A4, CRTC1, SLC25A3, SI, PHEX, KRT77, OPRM1, PHB2, CTNNA2, SERPINB6, RNF31, MAGEA1, ITM2B, RABGGTB, CDIPT, HSP90AA1, AMBP, MREG, ENDOU, ARID4A, AOC2, IGKV1-27, CYB5R1, WARS2, ZDHHC4, TMEM65, CLN3, PTPN20, MAL2, PSD4, WWP1, NCOA1, YWHAE, KLHL7, SLC27A6, GNG2, NIPA2, UBQLN1, KCNE4, BEST4, FGFRL1, PRKCH, GPR142, ABCC9, GPR87, BTN3A3, GFRAL, LANCL1, GRB7, CDH26, RTP2, RAB37, FXYD6, SLC2A9, SIGLEC8, F2RL3, KCNQ4, IRAK4, FLOT1, ATP8B1, PXDNL, GFRA3, SLCO1B1, IRAK1, EFNA2, CD47, CAPS, ENPP1, LRIG2, PRKCA, ITGA11, SHC3, RAB41, BEST3, GPM6A, SLC12A1, SLC8A2, FFAR1, GNG10, OR2AP1, HCK, IGKV3-20, IGLV3-9, RRP22, IGLV3-32, SLC22A24, PLA2G4C, EMILIN1, TAS2R60, SLC6A2, SERPINB4, MCHR1, EPGN, PRNP, HEG1, TAB2, IGKV1-8, SHC2, HDAC6, PPP4C, TSPAN33, ABTB1, TOM1, PIP5K1B, WDR73, PRMT8, GPR15, CPNE1, AMIGO2, XPR1, DTNB, RGS11, PLD1, NCR3, KAZN, MYO1B, PRKCQ, PALLD, SLCO5A1, NOXA1, SV2A, PILRA, SPRY1, PCDH12, CYS1, MUC12, SLC6A17, DLGAP1, EMP2, ADAMTSL1, ANKS4B, ERBB4, SLC26A1, POPDC2, EDA2R, RASGEF1B, SLC19A1, TMEM204, SLC22A13, PRKAR2B, IGKV1-37, C4BPB, STON1, MUC6, FES, C8G, FOLR2, CD34, PTGFR, G6PD, CHIC2, PALM |
| Biosynthesis of amino acids | GOT1L1, PYCR3, PKLR, PGK2, PFKM, GLUL, RPEL1, PRPS1, RPIA, TPI1, NAGS, GOT1, IDH1, TALDO1, ACY1 |
| Factor: MOVO-B; motif: GNGGGGG | CHD1, SSRP1, CCDC8, LUC7L2, TEX15, EIF4E3, NPHP3, SPEF2, MROH1, NSFL1C, UBE2G1, KATNB1, STARD9, F8A1, LARP1B, CDADC1, SLAIN1, UCK1, ATF1, MCM7, NACAD, KLHL18, CHEK1, GRSF1, RPS27A, ZNHIT2, LRRC42, CATSPER2, ASB7, SRGAP2B, TOP1MT, KBTBD4, TUBB2B, KIFC2, PRPF19, HOXC12, YY1AP1, UVSSA, USP47, FBXO38, SERTAD1, SSR4, BEX4, WDR47, BBS4, CHDH, ASXL2, SDHAF2, PYCR3, KPNA3, HIC2, PPP2R5D, AGO2, FKBP9, SRSF9, PHLDA1, ENDOD1, AACS, ARL2BP, ZFAND4, ATXN3, NPAS4, DPP9, ARMC8, S100A11, NKD1, MIA2, IGSF10, SPATA6, YAF2, CUEDC1, MTFP1, SOX3, SVBP, SNX4, S100A1, RANGRF, NEU2, KCNMB1, CTSZ, ORMDL3, PPP1R3G, FCHO2, TRPM5, GIPC1, KCNMB3, SH3GLB1, IL8RB, SFSWAP, KCNAB3, KCNG4, SLK, HCRTR1, DLG3, CSF1R, TSPAN4, PITPNM2, COL4A2, NLGN3, CPNE6, EFEMP2, TSLP, ARHGDIA, TMEM215, PKDCC, MARCKSL1, LIMD2, ACBD7, RAB27A, SLC9A3R2, AP2B1, SLC9A3, NXN, SERF1A, PDPN, WASHC5, CD93, MYO9A, OTOS, VASN, SOCS3, WFDC12, LGI2, CLIC6, ZPBP2, MYH9, GJB3, SEMA6B, NPC1L1, SMPD4, CAPZA1, PAK3, ARHGEF5, CXCL16, CNR1, DOCK6, PREX2, KANK2, ZBTB24, ANKRD36C, LMBRD2, COL1A2, FGGY, SERPINB9, ESRRA, CHST13, CYP27C1, CCDC92, FCGR2A, ZNF140, AHR, BHLHA9, GRK5, SBSPON, LRRC2, PLCD3, ARHGAP23, NFKBIB, ST18, GREB1L, DMAC2, ITFG1, RTL5, EVC, ERICH2, A1BG, NPY, IREB2, GALNT15, INTS5, MXD3, POLE, DNAJC22, SELENOF, HELLS, NR1D2, SLC43A3, AIDA, HTRA2, SECISBP2, TRIO, MT-CO1, NPTX2, EFS, WARS, NES, TGM2, BRD1, GADD45B, HEBP2, METTL7B, WDR83OS, FBXL20, SERPING1, SLC6A1, UPRT, PIN1, MOXD1, ZNF580, ALDH3A1, AKR1E2, CARS, FAM89B, JRK, APC, TMED5, KRT83, PGK2, RHBDD3, SP5, AFMID, TMBIM6, LATS1, NDUFB3, PTMS, NEIL2, ZDHHC18, RASSF4, UBFD1, DCP1B, C9ORF50, KBTBD11, POLDIP3, DRAP1, CDC42SE1, MLLT3, BAZ1B, INTS1, THAP7, DZIP1, ANKRD16, UBE2E1, E2F8, CAPRIN1, GAMT, MARF1, RASSF7, DYSF, CREB3L2, PNPLA7, HAGH, DCAF11, MRPL39, RANBP3, DCAF4, POLR2C, MBD6, ACTR8, TASOR, AGO1, NSMCE3, DDB1, BEX5, POMP, HAUS7, URB1, USB1, CKAP2L, SUPT3H, CAPZB, ST7, RANBP1, CCNG1, PIH1D2, TPST2, POLR3K, C1D, PFDN2, CEP192, AGO4, C2CD3, DDX25, MLLT6, CLPSL2, HSF5, LCORL, ARFGEF2, POLD2, TMX3, ARMCX3, MTG1, SWI5, RECQL, SPOPL, IPO13, EPB41L4A, EIF5A2, JADE2, KSR2, CSNK2B, CCDC68, RBM3, MCTS1, PRMT7, DDA1, IRX1, DENND2A, UBE3A, ELOVL1, FXR1, SMURF2, LYPLA2, BAG1, PDCD6, GOLT1B, SEC61G, PPM1H, TSSK1B, ATG16L2, ZIC2, MOSPD3, MOV10, KHSRP, DAP3, TRIM8, SESN3, DNAJB1, DACT1, NDRG1, BIRC7, ESYT1, SSR2, BOK, SLC36A4, ARHGEF10L, RBM46, HOXA4, CCDC170, SDK1, DAAM1, EPN3, JPH4, LTBP2, ARAP2, GCC2, FERMT2, PTPRT, KCNK9, INKA2, SBF2, FGF16, PDE8B, UNC5B, RAP1GAP, INSYN1, TRIL, MYOM3, FN1, MACO1, OSMR, CPLX2, APLN, RAB8B, EPHA1, RSPO3, SLC15A3, AZIN2, FSCN2, DNM2, STARD6, PDGFRL, CRMP1, SLC9A6, SERPINE1, RAB3IP, LUZP1, PIP5KL1, EPHA5, RTN4R, SMAGP, AP1G1, ATP2B2, MFAP4, RYR3, SPRY2, GPR34, CHRNB4, PLPPR4, EFNB2, BGN, SLC9A2, THBS3, WNT16, RIC8A, NDST2, IGF1R, CLDN12, WFIKKN1, RAPGEF4, HOOK1, SCARF2, PCDH1, OLFML3, FCN3, PATJ, DPEP3, JAM3, KIF5A, LOXL2, MYO9B, NUS1, ZDHHC8, HCN2, IFRD2, CRTAC1, FADS2, FOXL2, CGB5, GBP6, CGB8, C5ORF15, WSCD1, LDHC, TPT1, KANSL1L, UCP1, SCML1, HOMEZ, SNX32, THOP1, FBXO33, TRAF2, GSTT2B, CSRNP1, CMC4, ADRA2A, PRSS53, IKZF4, GABRB3, MANSC1, C6ORF47, UGT1A1, CHRND, GDNF, FKBP1B, ZNF215, C14ORF93, SLC25A52, PLEKHA6, CASD1, TIMP2, AP1S3, NBEAL1, SLC43A1, PLCD1, MBOAT1, CCDC122, LEKR1, GFI1B, LONRF2, XKR4, DIP2C, CXXC1, CRTC1, OAT, RETREG2, CFAP97, ITPA, ZNF799, PRMT1, NT5C1A, SCYL3, IKBKG, CTNNA2, SERPINB6, PM20D1, BDH1, RNF31, HDAC1, PARP9, ZNF148, STRN4, RPIA, ETF1, ITM2B, TPI1, CUTC, GPX1, PAPOLB, HSP90AA1, GCH1, SPIC, IRF1, NAGS, CTF1, ZNF628, BIK, TSTD1, LGALS1, GOT1, HPRT1, C1ORF56, MYRF, TMEM200B, STMN1, ADH5, DHX9, ARID4A, SCO2, DDX3Y, ZNF707, CDX2, LENG9, SNRNP25, KLF12, CCND2, ZNF652, C11ORF96, CAB39L, GLI4, HECTD4, DHRSX, AVEN, TIMM23, GTF2A1, PHF21A, DOLK, YBX2, CYB5R1, TAF7, ZNF277, RAD50, MRNIP, SFPQ, NXNL2, ALG11, PPP4R2, TEX2, TMEM65, ACTR10, CLN3, CCNE2, ZMYND15, MEIOC, NEIL3, MRTO4, MAL2, ENTPD5, SRSF3, TBC1D16, ZNFX1, FOXB2, HDGF, ARID4B, TFAP2E, WWP1, NCOA1, C4ORF46, YWHAE, KLHL7, ZNF467, RNASET2, B3GALT1, TP53INP2, MAP3K6, RRP15, NCOA5, C14ORF28, MAPK15, SSX4, YWHAG, NMNAT3, GOLGA2, SLC27A6, ADAMTS12, VPS26A, TMTC1, NXPE4, UBQLN1, KCNE4, FGFRL1, PRKCH, ABCC9, ASB10, LMTK2, PLXDC2, LANCL1, CXCL14, RAB37, FXYD6, SIGLEC8, F2RL3, KCNQ4, IRAK4, FLOT1, ATP8B1, CLEC16A, GFRA3, LGI4, IRAK1, EFNA2, FGF11, CD47, TXLNA, CAPS, ENPP1, LRIG2, PRKCA, ITGA11, SHC3, PRICKLE1, RAB41, GPM6A, SLC12A1, SLC8A2, TMTC2, RBP4, DDAH1, COL22A1, GNG10, MOGAT1, PLA2G12A, CYP21A2, MAFA, HCK, PNMT, RRP22, DPYS, TBX6, PHYKPL, PUDP, ZNF536, PCYT1B, RPUSD1, ZNF69, PYY, EMILIN1, NR0B1, SLC6A2, ESRRB, LAP3, AS3MT, SCT, RBM5, ELL, PANO1, PRNP, MAFF, EIF3K, HEG1, DPP7, PFAS, PBX3, TAB2, KLF3, FGF4, SHC2, NRL, ATOH1, HAUS3, KRT5, ALDH5A1, PPP4C, TSPAN33, PIAS1, ABTB2, DLEU1-AS1, NAP1L4, PUM1, TMEM214, ABTB1, SMARCD2, KRTAP5-5, NUDT17, PIP5K1B, CCDC110, FASTKD3, RBM15B, PCYT1A, PIMREG, WDR73, B4GALT7, RBMX2, FBXL3, SERTAD2, PEF1, CSNK2A3, DTX4, PPP2R2A, USP25, PELI1, CARNS1, PRMT8, TFEB, PTPN9, CPNE1, NEURL1B, TGFB1I1, USO1, LRATD1, XPR1, ARFRP1, DTNB, RFNG, SYT5, PRDX6, RGS11, PLPPR3, KCNK12, MYO1B, PRKCQ, SLCO5A1, NOXA1, SV2A, LMCD1, PLPPR2, SPRY1, FGF19, CYS1, SLC6A17, ERBB4, ERLEC1, SLC26A1, POPDC2, CERS5, EDA2R, SLC19A1, TMEM204, SLC22A13, FAM171A2, FAM120B, PRKAR2B, LCN9, TDRD10, CCDC148, CEBPA, STON1, MUC6, C8G, TBX15, CD34, ST6GALNAC3, FKBPL, IDH1, ALKBH3, CDK2, TALDO1, SAMD8, ASPG, PTGFR, NR2F6, G6PD, RPS16, FUOM, ZEB2, PALM, HMCES, OLIG2, ACY1, TNNC1, C12ORF42, TMEM220, NCBP2AS2, POU2F1, LRRC10B |
